# Rules of nature’s *Formula Run*: Muscle mechanics during late stance is the key to explaining maximum running speed

**DOI:** 10.1101/2020.10.29.361089

**Authors:** Michael Günther, Robert Rockenfeller, Tom Weihmann, Daniel F. B. Haeufle, Thomas Götz, Syn Schmitt

## Abstract

The maximum running speed of legged animals is one evident factor for evolutionary selection—for predators and prey. Therefore, it has been studied across the entire size range of animals, from the smallest mites to the largest elephants, and even beyond to extinct dinosaurs. A recent analysis of the relation between animal mass (size) and maximum running speed showed that there seems to be an optimal range of body masses in which the highest terrestrial running speeds occur. However, the conclusion drawn from that analysis—namely, that maximum speed is limited by the fatigue of white muscle fibres in the acceleration of the body mass to some theoretically possible maximum speed—was based on coarse reasoning on metabolic grounds, which neglected important biomechanical factors and basic muscle-metabolic parameters. Here, we propose a generic biomechanical model to investigate the allometry of the maximum speed of legged running. The model incorporates biomechanically important concepts: the ground reaction force being counteracted by air drag, the leg with its gearing of both a muscle into a leg length change and the muscle into the ground reaction force, as well as the maximum muscle contraction velocity, which includes muscle-tendon dynamics, and the muscle inertia—with all of them scaling with body mass. Put together, these concepts’ characteristics and their interactions provide a mechanistic explanation for the allometry of maximum legged running speed. This accompanies the offering of an explanation for the empirically found, overall maximum in speed: In animals bigger than a cheetah or pronghorn, the time that any leg-extending muscle needs to settle, starting from being isometric at about midstance, at the concentric contraction speed required for running at highest speeds becomes too long to be attainable within the time period of a leg moving from midstance to lift-off. Based on our biomechanical model we, thus, suggest considering the overall speed maximum to indicate muscle inertia being functionally significant in animal locomotion. Furthermore, the model renders possible insights into biological design principles such as differences in the leg concept between cats and spiders, and the relevance of multi-leg (mammals: four, insects: six, spiders: eight) body designs and emerging gaits. Moreover, we expose a completely new consideration regarding the muscles’ metabolic energy consumption, both during acceleration to maximum speed and in steady-state locomotion.

## 1. Introduction

Saying, in the twenty-first century of the Gregorian calendar, that the evolution of living organisms is driven by stochastically induced genetic mutations interplaying with the physical properties and constraints that form and encompass the organisms seems close to trivial. Yet, questions remain, at least regarding *which* exact properties and constraints (or, simply, *rules*) are the essential ones—that is, those which can *causally explain* the phenomenon of life in focus. This paper addresses the particular question which rules—meaning first principles, laws of nature, design criteria, and structural and material properties (characteristics, force laws)—most sensitively determine the maximum speed that an animal can achieve in legged terrestrial locomotion.

Our paper has been inspired by a recent publication [1]. In it, Hirt and colleagues have provided an extensive data compilation of 458 measured values of maximum running speeds, gathered from 199 animal species, which they plotted double-logarithmically against an animal’s body size. Such plots have long been used as the visualising counterpart of mathematically capturing, by means of a power law equation [2, 3], the effect of body size on any biological property—that is, the property’s allometric scaling: “…the structural and functional consequences of a change in size or in scale…” [4]. This power law approach is a suitable means of searching for criteria that drive body design, because different rules being predominat usually predict different power law exponents for a given measured quantity that is probed for size-dependency. A few design criteria have already been suggested as being effective in shaping the body of a running species [5]. Amongst these criteria, the most prominent are ‘geometric similarity’ [6], ‘elastic similarity’ [7], and ‘dynamic similarity’ [8].

The first and main issue of this paper is to causally explain—by means of a mechanistic, reductionist biomechanical model—the allometry of the maximum speed of legged running or hopping (legged speed allometry).

A salient feature of the legged speed allometry is its overall speed maximum occurring in species— pronghorns, antelopes, and felides, in particular [9]—that are far from being the largest on earth. This feature, being in accordance with the early data compilation provided by [9] and a recent one by [10], was the main object of interest in [1]. The authors of [1] assumed, first of all, that the amount of metabolic energy of ATP that is stored in white muscle fibres, and which is available for doing the mechanical work necessary to accelerate the body, is limited. Second, they suggested that the time until depletion of this ATP storage increases less with body size than the time that is required to accelerate to a likewise size-dependent theoretical speed maximum, with the latter being determined by other factors. Based on this line of reasoning, they introduced a speed-limiting multiplier to their assumed power law description of the legged speed allometry. This multiplier, of which the crucial parameter is the size-dependent time until depletion of the white fibres during acceleration, allowed them to calculate a critical body size with a corresponding overall speed maximum (limit).

As a second issue of this paper, we offer an alternative, mechanistic explanation for this overall speed limit, which comes from concentric muscle contraction mechanics during a leg’s contact phase with the ground rather than from the accumulative effects of muscle metabolism across consecutive steps. At this, we predict, from a model idea, the settle time that it takes the muscle to accelerate its own mass, after a change in force demand, to the very contraction velocity that corresponds to the new force level demanded. The relation of this settle time for muscle acceleration to the duration of the respective leg contacting the ground can explain the overall running speed limit in the very range where it occurs, as per the data compilations [1, 9, 10].

As a third issue, we use our biomechanical model to further explore which anatomical, geometrical, mechanical, and muscular properties can predict—and thus causally explain—species-specific deviations from the (mean) legged speed allometry.

We start by addressing our first and main issue: deriving the legged speed allometry from these properties.

## 2. Model formulation

In terrestrial, legged locomotion, the body of a moving animal experiences inevitable mechanical energy loss (dissipation) in the early phase of the stance duration of any single leg that has collided with a substrate surface (in running, usually the ground) at the instance of its touch-down (TD). Energy is dissipated during the dynamics in the wake of TD, which are usually simply termed ‘impact response’. The earliest response is a high-frequency acceleration peak of parts of the body mass (leg bones or cuticula) immediately after TD. Energy dissipation occurs due to material deformation, primarily within the distal leg masses [11]. First of all, the most distal pads are compressed [12, 13], which induces shock-waves that propagate through the leg [14] and, certainly, at least in humans, the whole body up to the head [15, 16]. Further energy loss originates [17, 18] from the short-range deformation of, and fluid flow in, soft tissue like leg muscles [19, 20], fatty pads, and organs as a consequence of the impact decelerating the skeleton to which the soft tissue and fluids are connected. To counteract the inevitable leg shortening after TD, more energy is dissipated by cross-bridges [21, 22] and tendons (hystereses: [23]) during longer-range, lower-frequency, eccentric contractions of leg muscles, as is by hemolymph (e.g., in spiders [24, sect. 1.2]) or other fluid flows. These lower-frequency deformations are also inevitable, because the vertical momentum of the body must be reversed by some repulsion dynamics acting against gravity. Backed by basic mechanics, others [25] have used exactly this line of reasoning to calculate speeds that minimise work per distance travelled. The latter minimisation is particularly useful when an animal is moving larger distances at cruising speed (roaming or migration). However, this criterion can certainly not be the major determinant in locomotory situations that demand maximum speed or acceleration (hunt or escape).

If only one leg is in contact with the ground, the forward (horizontal) component of the running velocity of a moving animal’s centre of mass (COM) must go through a minimum during the leg’s stance duration. In particular, this is because at least the lower-frequency dynamics come with an initially braking ground reaction force (GRF) component in the forward direction. An event such as a minimum in running velocity would then occur nearby midstance (MS), i.e., approximately half 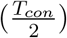 of the stance duration *T_con_* after TD. This transient minimum in running velocity around MS must be compensated for during late stance by the muscles’ re-feeding mechanical work in order to further accelerate the COM or finally maintain maximum running speed v_*max*_. Note here that, throughout the remainder of this paper, the term ‘muscle’, in fact, means a whole muscle-tendon complex if ‘muscle’ is used without any qualifier. In the mechanical steady-state of maintaining v_*max*_, the net mechanical energy balance of work done by the legs and energy losses by body-internal friction plus external drag forces equilibrates to zero during a gait cycle. In gaits with strongly synchronised legs, including one-leg ground contact phases, such a steady-state equilibration may also emerge within a single stance phase. If several legs are in contact with the ground, energy loss in one leg may be compensated for by work in another leg, and fluctuations around a speed steady-state can be maintained by means of a finite number of consecutive and cooperative legs’ stance phases. Yet, due to the general inevitability of material deformation after any collision, compensation will always occur by concentric muscle contractions during a single leg’s late stance.

According to this line of reasoning, our model represents the mechanical situation at the end of stance, that is, we start with the idea that we look at a situation close before a *single* leg’s lift-off (LO). A reductionist model approach always requires abstraction. Our object of investigation asks for apprehending the essence of the dynamics that are inherent to usually *several* legs interacting with the ground at the same time, as in a trot, alternating tripodal gaits, or metachronal as well as other multi-legged wave patterns.

Thus, our presentation of the problem ‘calculate maximum running speed’ so far asks for a conceptual step. In a straight forward way, this is introducing the top speed state (TOPSS), defined as the state in which (i) the re-fed amount of muscle work is just so sufficient to compensate for all dissipative losses in the wake of a leg’s TD, and which (ii) incorporates that the animal runs at its highest speed possible facing the correspondingly maximised drag force, when all forward acceleration ceases. TOPSS must occur after MS of a single leg, close to LO, and is the very mechanical state of a running animal that, per definition, *represents* v_*max*_. TOPSS is an instantaneous state of an intermediate speed maximum that represents a low amplitude fluctuation around a mean steady-state value of maximum running speed. Therefore, a v_*max*_ value predicted by a biomechanical model at TOPSS is well representative of an animal’s maximum running speed as usually measured in experiments.

Following up on the rationale of compensatory mechanical work, the key mechanical event that occurs before TOPSS is, then, the (crucial or representative) muscle under consideration being in the isometric state (ISOMS), which, indeed, occurs close to MS in any spring-mass-like locomotion [26, 27, 28, 29, 30]— see, e.g., the times of maximum joint flexion in human running [31, table 4] and sprinting [32, fig. 2], [33, fig. 1], and even in the long jump [34, 35]. As we keep in mind in the remainder of this paper that, for a start, TOPSS regularly occurs before, but close to, a single leg’s LO, we also memorise that the event of ISOMS occurs close to MS. Immediately connected to the TOPSS definition, ‘the leg’ of our model may in fact be a lumped one, which consists of more than one anatomical leg. Notwithstanding, a certain muscle in a certain anatomical leg may be identified that is the crucial one to contribute work at an instant of TOPSS as just defined by coupled muscle-COM kinetics.

**Figure 1:**
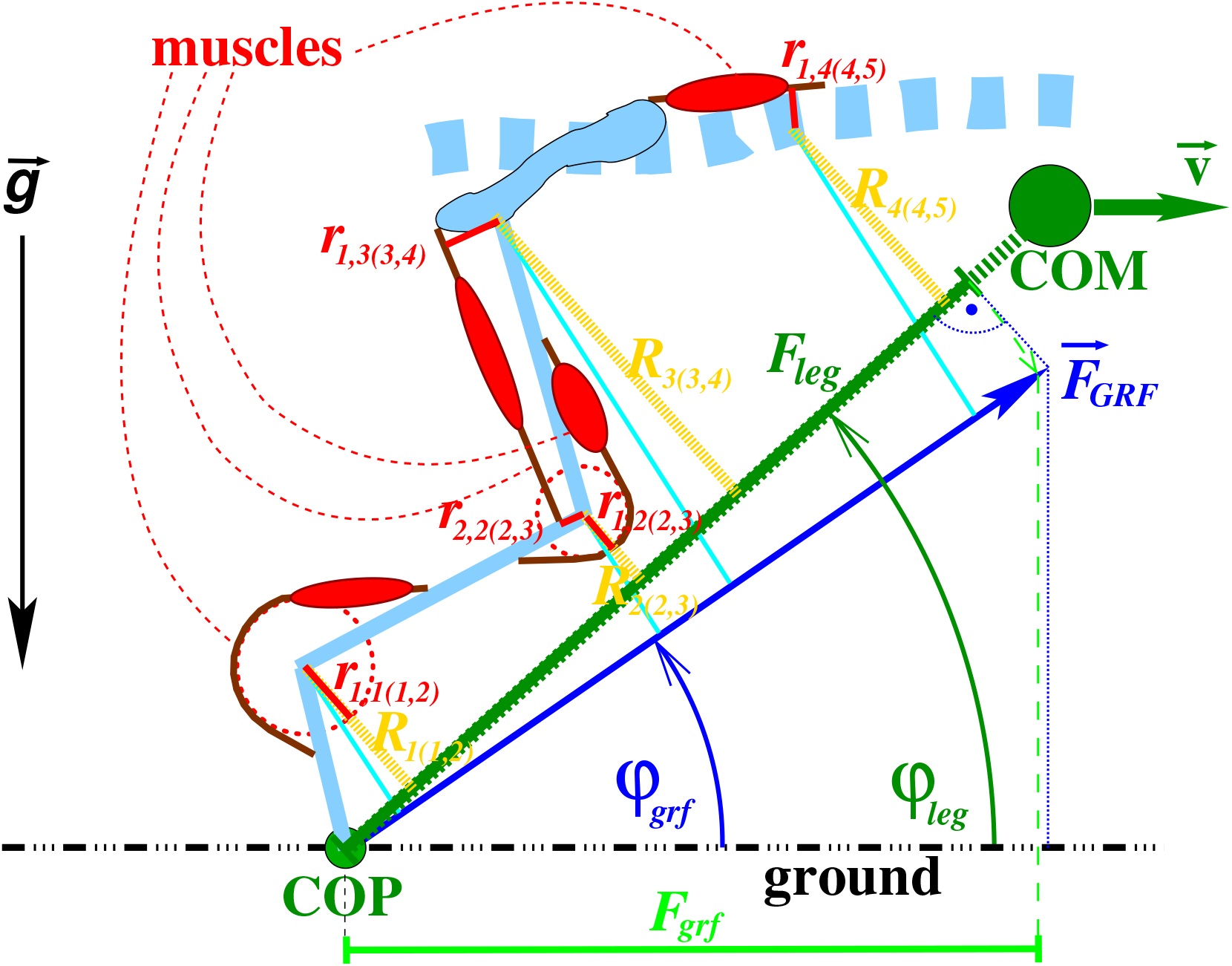
Model geometry, and its relation to running direction 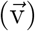 and ground reaction force (GRF) vector 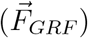. In the model plane, which is spanned by the vectors of gravitational acceleration 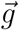 (perpendicular to the ground) and running direction (defined by 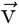), a single anatomical leg is depicted for clarity. A lumped leg may then consist of more than one anatomical leg. Here, this single anatomical leg even entirely equals a functional leg because only its anatomical tip, rather than a foot with finite contact area, touches ground: Its tip and the centre of pressure (COP) are equal. Generally, the instantaneous axis of a functional leg, be it a single or lumped one, is the line (thick dashed dark green) that connects the leg’s instantaneous location of its COP to the position of the body centre of mass (COM). The COP-COM distance is the (functional) leg length (Eq. (J.6)) *L_leg_* in our model. The instantaneous velocity 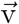 of the body COM is depicted by a horizontally oriented (dark green) arrow, which also represents the running speed, with v symbolising its magnitude. The COP of a net (lumped) leg would be located somewhere within the polygon that arises from the ground contact positions of all those anatomical legs that are assumed to form the lumped leg. The effective mechanical advantage (EMA) of each muscle *m* crossing the joint DOF *j*(*i, k*) in a functional leg is 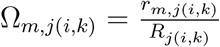, with *i* and *k* indicating the leg segments connected by the joint DOF *j*. The symbols *R_j_*_(*i,k*)_ (golden) and *r_m,j_*_(*i,k*)_ (red) denote the joint DOFs’ and muscles’ lever arms, respectively. If *j*(*i, k*) is an angular joint DOF, then the (kinematic) *R_j_*_(*i,k*)_ is the perpendicular distance between its axis of rotation and the axis of its respective functional leg, which is in contact with the ground. The DOFs’ lever arms for kinematic (thick dashed golden lines) and force (cyan lines) gearing differ if 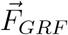 (blue arrow) and leg axis misalign, the axial leg force *F_leg_* (dark green line with caps) is the projection of 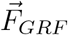 onto the leg axis (Eq. (11)). A muscle lever arm *r_m,j_*_(*i,k*)_ measures how much the length of muscle *m* changes if the coordinate of a joint DOF *j*(*i, k*) is changed. For a rotational joint DOF (angle), it is the perpendicular distance (red lines) between the muscle’s line of action and the axis of rotation. The instantaneous angles of the leg vector—the leg axis—and the respective GRF vector 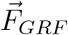 in the model plane, i.e., cos *φ_leg_* and cos *φ_grf_*, respectively, are also drawn, as well as the projection *F_grf_* onto the running direction (green line with caps; see Eq. (2)) of the vector with magnitude *F_leg_* that points in GRF direction (Eq. (16) from Eq. (15) with Eq. (12)). Our initial guesses of both cosines as functions of body size (mass) *M* (App. D.3) are plotted in Fig. 6. In our model, multiple joints and muscles are lumped into a net, size-dependent function Ω(*M*) (App. D.2, Fig. 5).

**Figure 2:**
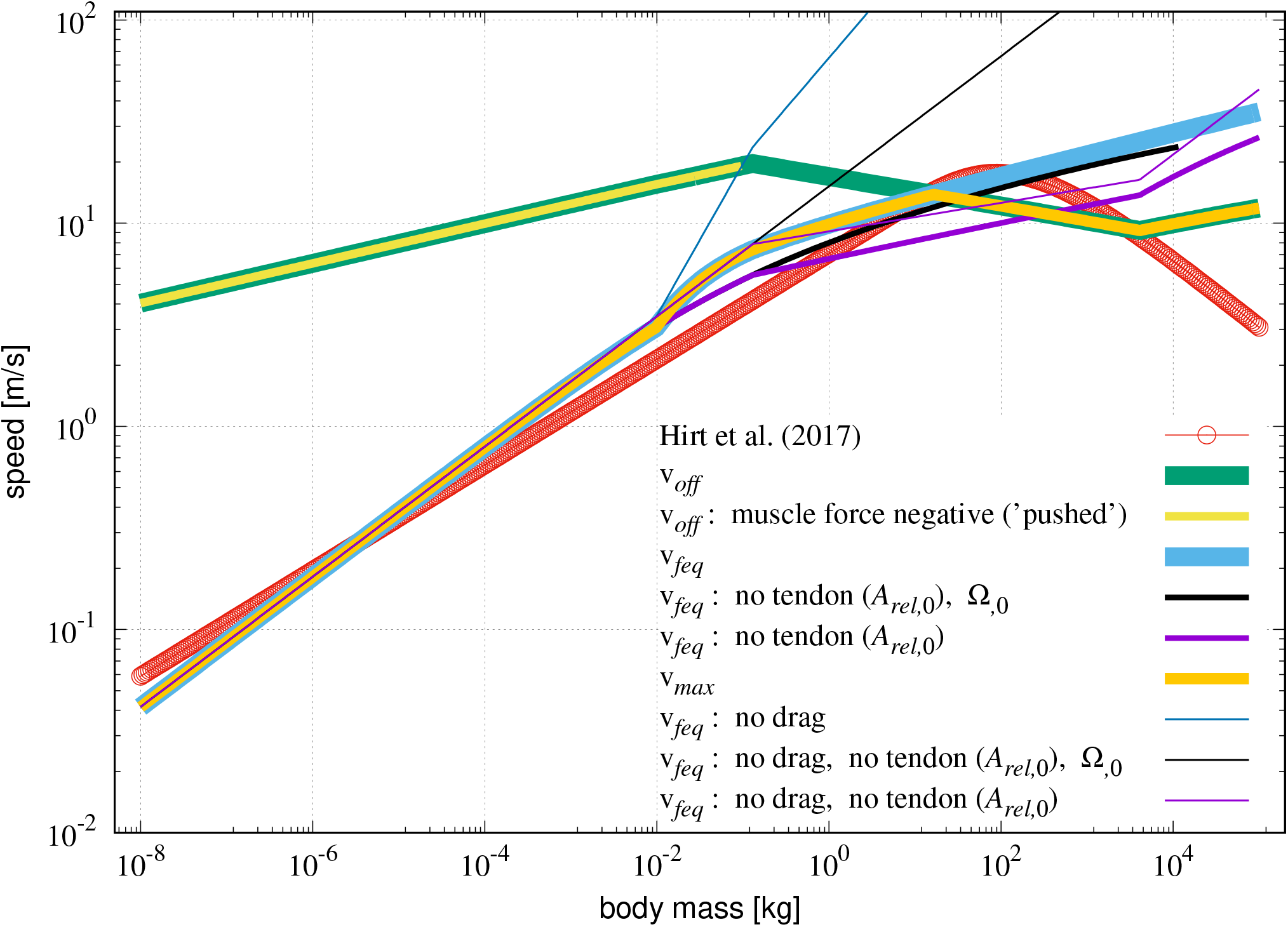
Model solutions (output): Maximum running speed v_*max*_ (thick orange line) vs. body mass *M*, using our initial guess (Table 2, third row) of parameter values. Two conditions or constraints, respectively, are suggested to determine v_*max*_ at any body mass value *M*: the force equilibrium (Eq. (2)) between (transformed) muscular driving force and external air drag (solution v_*feq*_: Eq. (H.4); very thick blue line) and the equality (Eq. (18)) between half the stance duration and finite available settle time for accelerating the muscle to high concentric muscle contraction velocities (solution v_*off*_: Eq. (H.7); very thick green line). As both speed-constitutive conditions depend on exactly the same property, the muscle’s force-velocity relation, the most limiting condition—i.e., the one with the lowest speed value predicted (Eq. (21))—determines the solution v_*max*_ that can be realised in nature (thick orange line). With our initial parameter guess, the model predicts that the solution changes from v_*feq*_ to v_*off*_ (kink in the thick orange line) for an average animal with body mass 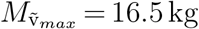 running at a speed of 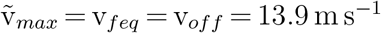. At any body size above this crossing point of both alternative solutions, muscle inertia is predicted to lead to premature termination of muscle work, because the leg is forced to lift off the ground before the muscle has accelerated to the concentric contraction velocity that would just so balance air drag. For a comparison, the curve fitted by [1] through 458 experimental data points, gathered from 199 species, is also plotted (thin red line with circles: overall 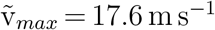 at 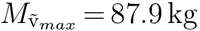). Note that the kink (at *M* = 0.1 kg) in the solution v_*off*_ to the duration constraint Eq. (18) is due to the EMA Ω commencing to change at this body size. Note also that it is just by coincidence that the solution v_*feq*_ to the force equilibrium Eq. (2) assuming *no air drag* (‘no drag’: thin dark blue line) crosses very close to this kink. The latter solution is the muscle’s *V_m,max_* transformed to v by the factor 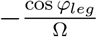 (Eq. (8)): Without air drag, running speed would be solely determined by muscle-internal damping, nonetheless limited by the duration constraint. Further model solutions for other limit cases are likewise plotted and explained in App. L, e.g., the kink in v_*off*_ solution at about 4 · 10^3^ kg.

A second abstraction is required, yet possible because the re-feeding of work is done by muscles. The leg force—and, thus, the forward-driving GRF component—is generated by a concert of multiple muscles even within a single leg. Here, again, the lumping concept of ‘one muscle’ helps. For one thing, the same applies as for the concept of ‘one leg’: either some blur due to lumping may be accepted for a start or the case of ‘the last one standing’ may be considered, that is, a certain (crucial) muscle still doing accelerating work on the COM at an instant of TOPSS. For another thing, lumping of multiple muscles crossing multiple joints in a leg can be particularly justified because skeletal muscles share a common property across species, muscle types, and even dynamic contraction conditions [36, 37, 38]: the relation between the force of a muscle and its concentric contraction velocity is a hyperbola, the ‘Hill relation’. This biomechanical muscle characteristic is named after A. V. Hill, who had first formulated it [39] on the basis of measurements of concentric steady-state contractions of dissected frog muscle fibres. We suggest, by use of the reductionist biomechanical leg-and-body model here, that the Hill relation, extended to non-steady states [36, 37, 38] of muscles with even tendons [40], plays a crucial role in explaining maximum running speed.

As a third abstraction, the concept of a ‘centre of pressure’ (COP, see e.g., [41, 42, 43]) that describes the mechanical interaction of two bodies—here, the animal’s body and the earth—at a substrate surface (e.g., the ground) is generally applicable. A *functional* leg’s axis can be defined as the difference vector from the instantaneous COP location at the distal end of an anatomical leg to the instantaneous position of the whole body’s COM, with the body COM thus constituting any functional leg’s proximal endpoint (Fig. 1). The magnitude of this vector is the (functional) leg length *L_leg_*. The COP concept still applies consistently when one is lumping together an arbitrary number of legs that are in contact with the ground at the same time [43]: The overall COP of the lumped leg is the weighted sum of the COP locations of all single leg’s making the lumped leg, with the weight being the respective single leg’s GRF component normal to the plane [41, 42, 44].

All these introductory remarks facilitate punctuating the basic idea behind our model: Whatever the balance of mechanical energy contributions during a leg’s stance phase, those leg muscles that can, by contracting concentrically, still release non-vanishing mechanical power *F_m_* · v_*m*_ at late stance must be the cause of the positive mechanical work

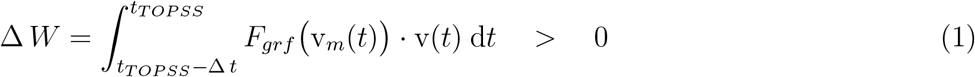

done to accelerate the COM in the short but finite period Δ *t* of time *t* before a maximum in COM velocity. Here, the symbols v_*m*_ and *F_m_* are a muscle’s contraction velocity and force, respectively, and *F_grf_* (v_*m*_(*t*)) and v(t) are the instantaneous components of GRF and COM velocity in the running direction. If no other event interferes, the instant *t* = *t_TOPSS_* of ceasing COM acceleration occurs when countering drag force *F_d_* (v(*t*)) equals *F_grf_* (v_*m*_(*t*)) (see Eq. (2) below), while positive (accelerating) power *F_grf_* (v_*m*_(*t*))·v(*t*) is still provided by muscles (see the inequality sign in Eq. (1)). The concentrically contracting, finishing muscles must, therefore, be crucial co-authors of the rules that determine the running state achieved, be it for further acceleration or in a steady state across steps. The latter is what we are primarily asking for in the present model approach—namely, finding TOPSS, as defined by Eq. (2) rooted in Eq. (1). In TOPSS, the COM velocity v is at its maximally possible steady-state value, that is, the body at maximum running speed v_*max*_. There is a further necessary condition for making it possible for these muscles—and, hence, the leg—to do work: The kinematic constraint between COM velocity and axial leg lengthening must be kept up. In other words, without the proximal end of the leg following the COM, any leg contact with the ground at its distal end would immediately cease. This is already suggestive of a muscle’s maximum contraction velocity being a determining factor for maximum running speed.

As said above, the instantaneous COM velocity v(*t*) is at its minimum during leg stance some time around an instant of MS or ISOMS, respectively, and at its maximum at an instant of TOPSS. This TOPSS definition reflects a *force equilibrium between non-zero drive and drag forces* (zero forward acceleration). This force equilibrium can generally be derived from the definition of mechanical work (Δ *W*: Eq. (1), left equality) by the steady-state rationale that the instantaneous mechanical powers generated (the integrand *F_grf_* (*t*) · v(*t*) in Eq. (1)) and dissipated (*F_d_*(*t*) · v(*t*)) equal at a finite v(*t*) value. All modes of legged locomotion (gaits) can be comprehensively examined by the use of the more abstractly defined running system states ISOMS and TOPSS. These states may shine through more distinctly in alternating gaits, which deploy a single leg or a strongly synchronised set of legs [26, 27, 45, 46, 47, 48, 49, 50], respectively, or may be more difficult to discern in multiple-leg gaits with strongly overlapping ground contact phases [51, 52, 53, 54, 55, 56].

In our concrete leg-body model, which we introduce now, the forward component of the GRF is assumed to be opposed solely by a drag force while the animal is running in a surrounding medium (air). All model properties (parameters) are assumed to either scale with size (body mass *M*) or be constants.

### 2.1. Force equilibrium, maximum running speed, and body size

According to the considerations above, the forward components of body-external, movement-resisting drag force (symbol *F_d_*) and driving GRF (symbol *F_grf_*: the body-internally generated leg force transmitted to the COM via leg-ground interaction) equilibrate at TOPSS:

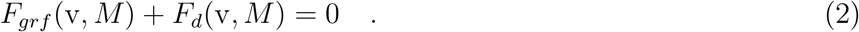

At this force equilibrium, the net body COM acceleration in running direction vanishes. The solution to Eq. (2) for the only free variable v provides

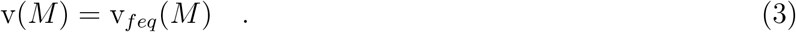

Appendix G explains in more detail how the force equilibrium Eq. (2) writes explicitly (Eq. (G.1)) in terms of our model parameters and running speed v. In particular, Eq. (G.1) turns out to be a cubic equation (Eq. (G.2)) for speed (*z* = v), which can be solved (Eq. (H.4)) in closed form. If no other constraint interferes, the solution Eq. (3) of the force equilibrium mathematically represented by Eq. (2) is, thus, our primary model prediction of maximum running speed:

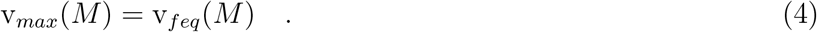

As will be explained in Sect. 2.7 further below, there is, indeed, a second condition or constraint (Eq. (18)), respectively, beyond the force equilibrium Eq. (2), which can limit maximum running speed v_*max*_ and which arises from the same muscular force-velocity properties that determine not only concentric contractions but also muscle mass acceleration. Above a certain body size, this constraint (Eq. (18)) becomes more speed-limiting than the force equilibrium (Eq. (2)), and, thus, interferes with it in fact.

In the first instance, however, we assume that the maximum speed a running animal can achieve is given by the physical and physiological properties that determine the opponents in the force equilibrium Eq. (2): *F_grf_*, i.e., the body-internal driving and dragging (visco-elastic) forces in the muscles, and *F_d_*, i.e., the body-external drag forces imposed by the environment (here, air drag with both friction and pressure contributions). To that end, we assume, in our model approach, that the two forces *F_grf_* (v, *M*) and *F_d_*(v, *M*) depend on running velocity v as well as on a set of parameters representing physical and physiological properties, with part of these parameters, in turn, depending on body size as represented by body mass *M*, like in others’ approaches [1, 57, 58]. Accordingly, if we found, in the literature, empirical indications of a significant size-dependency of any one model parameter, we treated such a potentially size-dependent parameter as a function of body mass *M* (a mathematical power law, see App. A), which is explicitly indicated by a parameter’s functional dependency on *M* (in parenthesis) in our model description. We apply our model calculations to body mass values that range from *M* = 10^−8^ kg (mites [59]) to *M* = 10^5^ kg (heaviest dinosaurs [60]).

### 2.2. Body mass and characteristic length

As emphasised in the introduction, our first main issue is to determine how maximum running speed v_*max*_ depends on body size. That is, we aim to derive, symbolically, a mathematical formulation of the dependency of v_*max*_ on physical and structural parameters, as well as the sole independent variable ‘body mass’ *M* in our model: the relation v_*max*_ = v_*max*_(*M*). For this, we further suggest specifying the characteristic body length *L_bod_* as the trunk’s dimension, i.e., its longest dimension in particular. As an alternative, *L_bod_* might be more abstractly defined, for example, as twice the longest half-axis of the ellipsoid representing the inertia tensor of all body mass. In our model, we relate body mass and characteristic body length by

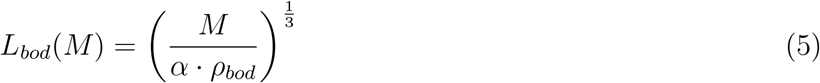

in which *ρ_bod_* is the body’s mass density (here, water: *ρ_bod_* = 10^3^ kg m^−3^). Our initial guess *α* = 0.12 was estimated using human anthropometric databases [61, 62] by means of in-house software *calcman* [63, 64], which calculates body segmental masses and dimensions from the given body mass *M*, body height, and sex.

### 2.3. Forces, kinematics, and model parametrisation

Modelling air drag *F_d_*(v, *M*) requires five parameters (see App. B for details). Modelling the relation between forward-driving GRF component *F_grf_* (v, *M*) and body-internal driving (muscle) forces requires some more sophistication, in particular, the formulation of the gearing between muscle force (*F_m_*) and COM dynamics (*F_grf_*), which is mediated by muscle and leg anatomy and geometry. For this, we use the concepts of *effective mechanical advantage* [58] (EMA; definition in App. C; mathematical symbol Ω) and *axial leg force* (*F_leg_*). The crucial leg properties in legged locomotion are its skeletal structure (bones and joints) and the properties of its actuators (muscles). In the following, we explain the main transformational steps that relate muscle contraction to COM movement while leaving some details about model parametrisation and choice of values to the Appendix. Any initial guesses of model parameter values not immediately mentioned in their introductory text passages can be found in App. D. All model parameters are listed in Table 1, together with a short explanation, and also in Table 2, together with their initially guessed values. The relation between rate of length change (velocity v_*m*_) of a muscle and its contractile force *F_m_* is modelled here, in a well-nigh classical way, by Hill’s [39] hyperbolic force-velocity relation (Eq. (I.1), Eq. (I.2), Eq. (I.3), or Eq. (I.4), respectively) formulated as Eq. (E.1) in our model-specific parametrisation *F_m_* = *F_m_*(v_*m*_, *M*) (see App. E).

**List of symbols and model parameters**

**Table 1:**
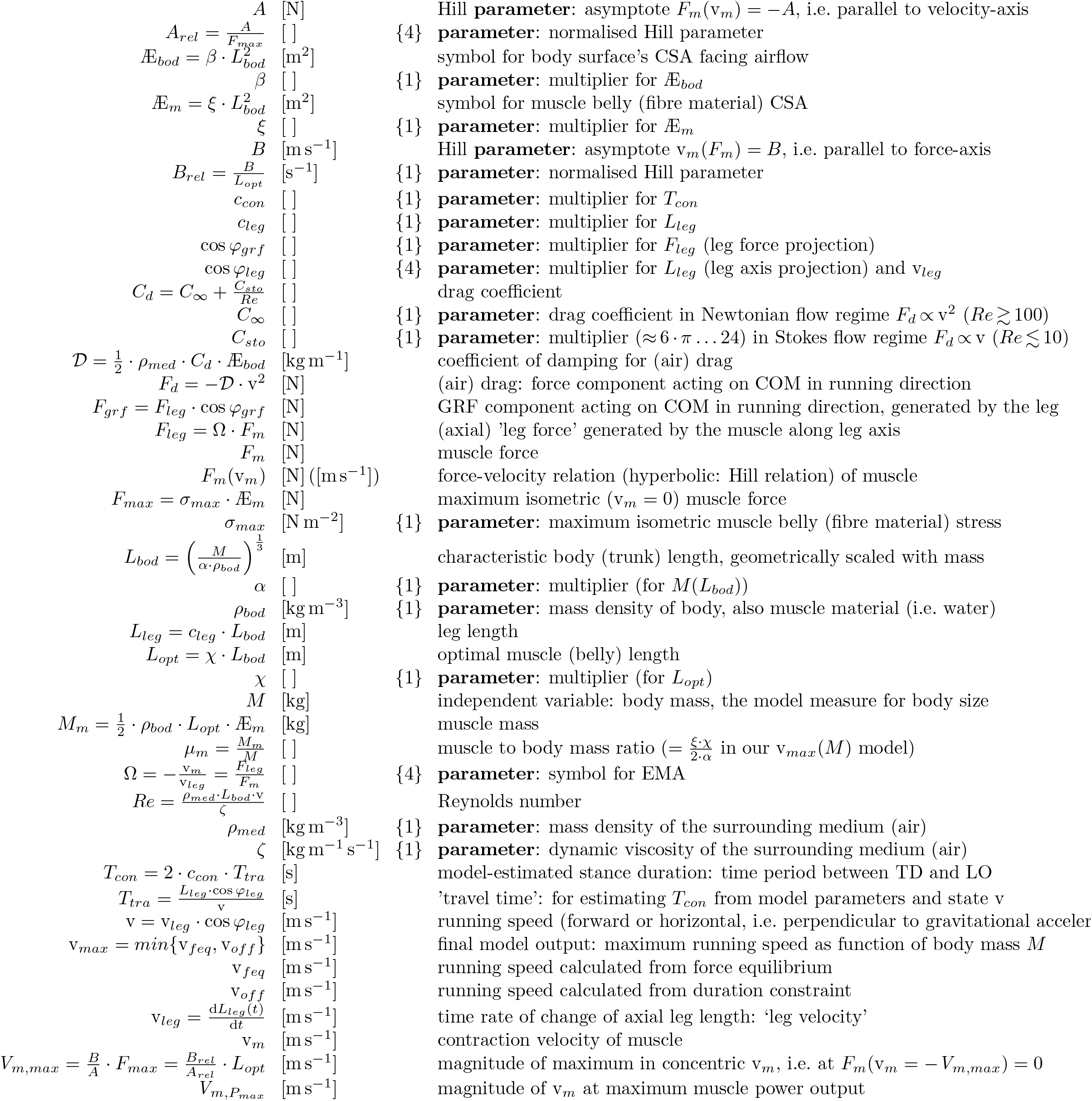
Parameters of the biomechanical running model. Units are given in the second column. For any free biomechanical model **parameter** *p_n_* (indicated by *n*), the third column reports in curly brackets the actually required number of parameters to mathematically describe its allometry. If the number is 1, the parameter value does not depend on body size as represented by body mass *M*. Otherwise, the parameter *p_n_*(*M*) is considered dependent on *M*, in terms of a power law (see App. A). Describing a power law requires two parameters 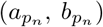. If applicable, a third and fourth parameter may be introduced by fixing a lower 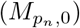 and an upper 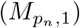 boundary, respectively, of power law applicability, while then implying the values *p_n,_*0 and *p_n,_*1 of *p_n_*(*M*) at their respective boundaries to also apply beyond (*p_n_*(*M*) taking *p*_*n*,0_ or *p*_*n*,1_, respectively). Note that the term ‘muscle’ means a whole muscle-tendon complex if ‘muscle’ is used without any qualifier.

|  |  |  |
| --- | --- | --- |
| $A$ | [N] | Hill **parameter**: asymptote $F_m(v_m) = -A$, i.e. parallel to velocity-axis |
| $A_{rel} = \frac{A}{F_{max}}$ | [ ] | {4} **parameter**: normalised Hill parameter |
| $\mathcal{A}_{bod} = \beta \cdot L_{bod}^2$ | [m<sup>2</sup>] | symbol for body surface’s CSA facing airflow |
| $\beta$ | [ ] | {1} **parameter**: multiplier for $\mathcal{A}_{bod}$ |
| $\mathcal{A}_m = \xi \cdot L_{bod}^2$ | [m<sup>2</sup>] | symbol for muscle belly (fibre material) CSA |
| $\xi$ | [ ] | {1} **parameter**: multiplier for $\mathcal{A}_m$ |
| $B$ | [m s<sup>-1</sup>] | Hill **parameter**: asymptote $v_m(F_m) = B$, i.e. parallel to force-axis |
| $B_{rel} = \frac{B}{L_{opt}}$ | [s<sup>-1</sup>] | {1} **parameter**: normalised Hill parameter |
| $c_{con}$ | [ ] | {1} **parameter**: multiplier for $T_{con}$ |
| $c_{leg}$ | [ ] | {1} **parameter**: multiplier for $L_{leg}$ |
| $\cos \varphi_{grf}$ | [ ] | {1} **parameter**: multiplier for $F_{leg}$ (leg force projection) |
| $\cos \varphi_{leg}$ | [ ] | {4} **parameter**: multiplier for $L_{leg}$ (leg axis projection) and $v_{leg}$ |
| $C_d = C_\infty + \frac{C_{sto}}{Re}$ | [ ] | drag coefficient |
| $C_\infty$ | [ ] | {1} **parameter**: drag coefficient in Newtonian flow regime $F_d \propto v^2$ ($Re \gtrsim 100$) |
| $C_{sto}$ | [ ] | {1} **parameter**: multiplier ($\approx 6 \cdot \pi \dots 24$) in Stokes flow regime $F_d \propto v$ ($Re \lesssim 10$) |
| $\mathcal{D} = \frac{1}{2} \cdot \rho_{med} \cdot C_d \cdot \mathcal{A}_{bod}$ | [kg m<sup>-1</sup>] | coefficient of damping for (air) drag |
| $F_d = -\mathcal{D} \cdot v^2$ | [N] | (air) drag: force component acting on COM in running direction |
| $F_{grf} = F_{leg} \cdot \cos \varphi_{grf}$ | [N] | GRF component acting on COM in running direction, generated by the leg |
| $F_{leg} = \Omega \cdot F_m$ | [N] | (axial) ‘leg force’ generated by the muscle along leg axis |
| $F_m$ | [N] | muscle force |
| $F_m(v_m)$ | [N] ([m s<sup>-1</sup>]) | force-velocity relation (hyperbolic: Hill relation) of muscle |
| $F_{max} = \sigma_{max} \cdot \mathcal{A}_m$ | [N] | maximum isometric ($v_m = 0$) muscle force |
| $\sigma_{max}$ | [N m<sup>-2</sup>] | {1} **parameter**: maximum isometric muscle belly (fibre material) stress |
| $L_{bod} = \left(\frac{M}{\alpha \cdot \rho_{bod}}\right)^{\frac{1}{3}}$ | [m] | characteristic body (trunk) length, geometrically scaled with mass |
| $\alpha$ | [ ] | {1} **parameter**: multiplier (for $M(L_{bod})$) |
| $\rho_{bod}$ | [kg m<sup>-3</sup>] | {1} **parameter**: mass density of body, also muscle material (i.e. water) |
| $L_{leg} = c_{leg} \cdot L_{bod}$ | [m] | leg length |
| $L_{opt} = \chi \cdot L_{bod}$ | [m] | optimal muscle (belly) length |
| $\chi$ | [ ] | {1} **parameter**: multiplier (for $L_{opt}$) |
| $M$ | [kg] | independent variable: body mass, the model measure for body size |
| $M_m = \frac{1}{2} \cdot \rho_{bod} \cdot L_{opt} \cdot \mathcal{A}_m$ | [kg] | muscle mass |
| $\mu_m = \frac{M_m}{M}$ | [ ] | muscle to body mass ratio ($= \frac{\xi \cdot \chi}{2 \cdot \alpha}$ in our $v_{max}(M)$ model) |
| $\Omega = -\frac{v_m}{v_{leg}} = \frac{F_{leg}}{F_m}$ | [ ] | {4} **parameter**: symbol for EMA |
| $Re = \frac{\rho_{med} \cdot L_{bod} \cdot v}{\zeta}$ | [ ] | Reynolds number |
| $\rho_{med}$ | [kg m<sup>-3</sup>] | {1} **parameter**: mass density of the surrounding medium (air) |
| $\zeta$ | [kg m<sup>-1</sup> s<sup>-1</sup>] | {1} **parameter**: dynamic viscosity of the surrounding medium (air) |
| $T_{con} = 2 \cdot c_{con} \cdot T_{tra}$ | [s] | model-estimated stance duration: time period between TD and LO |
| $T_{tra} = \frac{L_{leg} \cdot \cos \varphi_{leg}}{v}$ | [s] | ‘travel time’: for estimating $T_{con}$ from model parameters and state $v$ |
| $v = v_{leg} \cdot \cos \varphi_{leg}$ | [m s<sup>-1</sup>] | running speed (forward or horizontal, i.e. perpendicular to gravitational acceleration) |
| $v_{max} = \min\{v_{feq}, v_{off}\}$ | [m s<sup>-1</sup>] | final model output: maximum running speed as function of body mass $M$ |
| $v_{feq}$ | [m s<sup>-1</sup>] | running speed calculated from force equilibrium |
| $v_{off}$ | [m s<sup>-1</sup>] | running speed calculated from duration constraint |
| $v_{leg} = \frac{dL_{leg}(t)}{dt}$ | [m s<sup>-1</sup>] | time rate of change of axial leg length: ‘leg velocity’ |
| $v_m$ | [m s<sup>-1</sup>] | contraction velocity of muscle |
| $V_{m,max} = \frac{B}{A} \cdot F_{max} = \frac{B_{rel}}{A_{rel}} \cdot L_{opt}$ | [m s<sup>-1</sup>] | magnitude of maximum in concentric $v_m$, i.e. at $F_m(v_m = -V_{m,max}) = 0$ |
| $V_{m,P_{max}}$ | [m s<sup>-1</sup>] | magnitude of $v_m$ at maximum muscle power output |

**Table 2:**
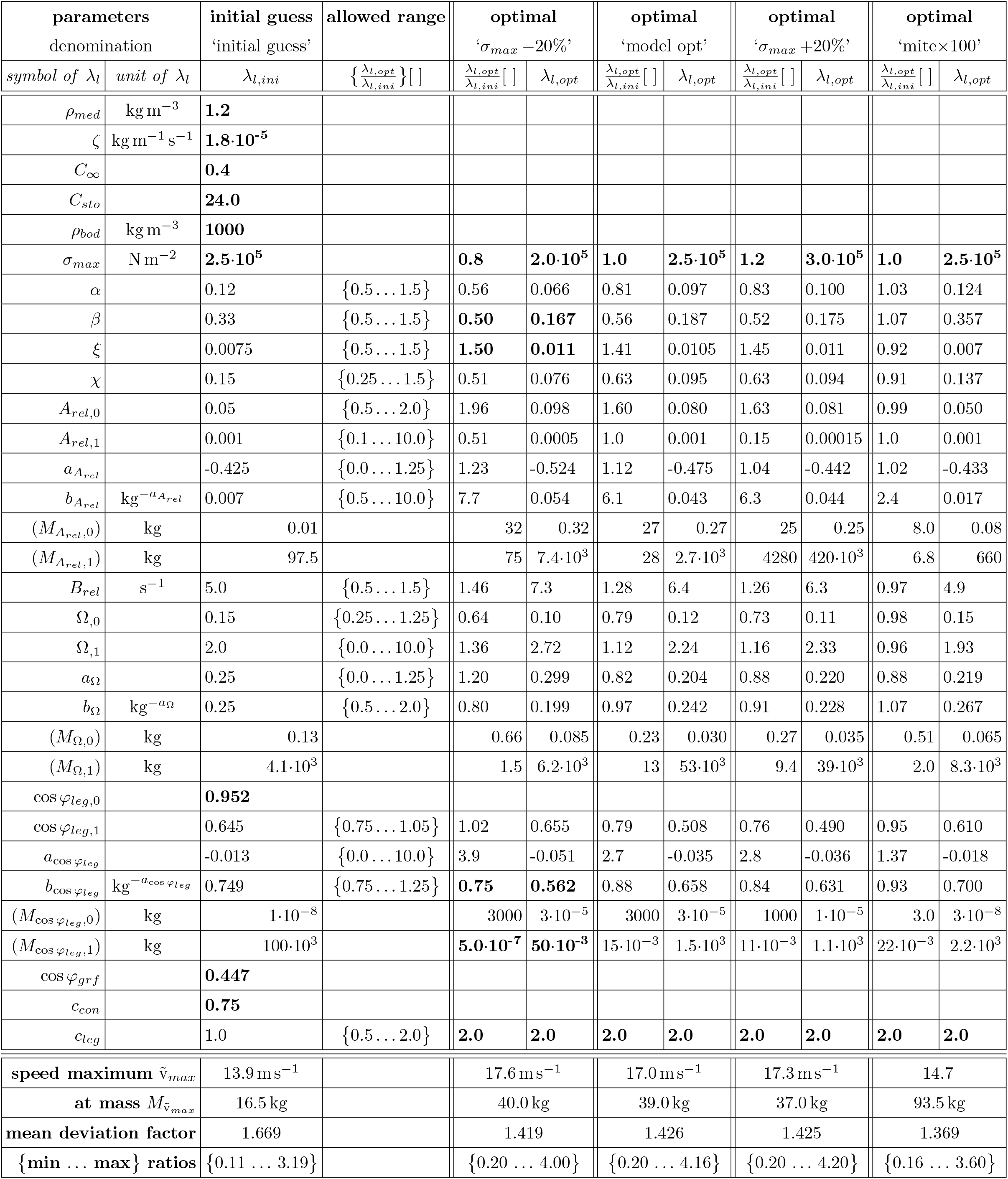
Compilation of model parameter values (as a synopsis, see Table 1: **List of symbols and model parameters**). In case a parameter *p_n_* is assumed to be *M*-dependent (*p_n_*(*M*)), the corresponding power law parameters 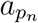 and 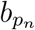 (see Eq.(A.2)) are given. If applicable, we likewise provide its boundary value(s) (*p_n,_*_0_, *p_n,_*_1_) and, as more illustrative, the body mass values at the lower 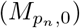 and upper 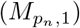 boundaries, though we do so in parentheses because this is redundant parameter information: 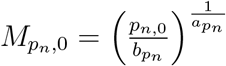 and 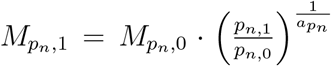. Column three reports our initial guess of model the parameter values extracted from the literature. Within the curly brackets, the fourth column reports the boundaries for the multiplier to the initial guess of a parameter value within which the optimising algorithm was allowed to select the multiplier value during least-square minimisation of the deviation of the model solution from the experimental data points as compiled by [1]. The fifth and sixth columns, respectively, contain the selected optimal values of multipliers and parameters for the maximum isometric muscle stress fixed to *σ_max_* = 2.0 · 10^5^ N m^−2^ (see App. E). For *σ_max_* fixed to our initial guess *σ_max_* = 2.5 · 10^5^ N m^−2^ (Fig. 4), optimal values are given in the seventh and eighth columns, respectively. In columns nine and ten, values are reported for fixed *σ_max_* = 3.0 · 10^5^ N m^−2^. In columns eleven and twelve, optimised values are reported again for the case *σ_max_* = 2.5 · 10^5^ N m^−2^, now with the four data points of the smallest animals (mites) weighted a hundredfold; this emulates a situation in which approximately the same amount of experimental data is available for very small animals as for larger ones [1] (197 species providing *N_exp_* = 455 points, see again Fig. 4). Values are reported in bold font if they were either fixed a priori or found by the optimising algorithm to be within less than one percent near the boundaries of the range allowed to the optimising algorithm. The leftmost kink in the legged speed allometry v_*max*_(*M*) (Figs. 2,5,4) indicates our finding of an *overall maximum in running speed* 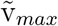 (corresponding critical mass: 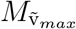) across all sizes. We quantified the deviation of an experimentally determined data point (index ‘*i*’: *M_i_*, v_*exp,i*_) from the respective point on the model solution (v (*M*)) by calculating the 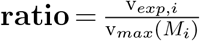. To quantify the overall deviation of a model solution from all experimental data points (**mean deviation factor**), we calculated the arithme)tic mean of all *N_exp_* data points’ **factor**_*i*_ = (**ratio**_*i*_) if v_*exp,i*_ >= v_*max*_(*M_i_*) or 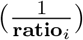 if 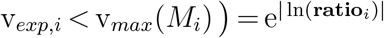. For comparison, the fit by Hirt et al. [1] (Fig. 4) provides an overall **speed maximum** 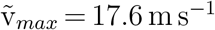 at mass 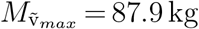, with a **mean deviation factor** of 1.417 and respective {**min**…**max**} **ratios** of {0.19…3.68}.

| parameters<br>denomination | | initial guess<br>'initial guess' | allowed range | optimal<br>'$\sigma_{max}-20\%$' | | optimal<br>'model opt' | | optimal<br>'$\sigma_{max}+20\%$' | | optimal<br>'mite$\times 100$' | |
| --- | --- | --- | --- | --- | --- | --- | --- | --- | --- | --- | --- |
| *symbol of $\lambda_l$* | *unit of $\lambda_l$* | $\lambda_{l,ini}$ | $\left\{\frac{\lambda_{l,opt}}{\lambda_{l,ini}}\right\}[]$ | $\frac{\lambda_{l,opt}}{\lambda_{l,ini}}[]$ | $\lambda_{l,opt}$ | $\frac{\lambda_{l,opt}}{\lambda_{l,ini}}[]$ | $\lambda_{l,opt}$ | $\frac{\lambda_{l,opt}}{\lambda_{l,ini}}[]$ | $\lambda_{l,opt}$ | $\frac{\lambda_{l,opt}}{\lambda_{l,ini}}[]$ | $\lambda_{l,opt}$ |
| $\rho_{med}$ | $\text{kg m}^{-3}$ | **1.2** | | | | | | | | | |
| $\zeta$ | $\text{kg m}^{-1} \text{s}^{-1}$ | **1.8$\cdot 10^{-5}$** | | | | | | | | | |
| $C_\infty$ | | **0.4** | | | | | | | | | |
| $C_{sto}$ | | **24.0** | | | | | | | | | |
| $\rho_{bod}$ | $\text{kg m}^{-3}$ | **1000** | | | | | | | | | |
| $\sigma_{max}$ | $\text{N m}^{-2}$ | **2.5$\cdot 10^5$** | | **0.8** | **2.0$\cdot 10^5$** | **1.0** | **2.5$\cdot 10^5$** | **1.2** | **3.0$\cdot 10^5$** | **1.0** | **2.5$\cdot 10^5$** |
| $\alpha$ | | 0.12 | $\{0.5 \dots 1.5\}$ | 0.56 | 0.066 | 0.81 | 0.097 | 0.83 | 0.100 | 1.03 | 0.124 |
| $\beta$ | | 0.33 | $\{0.5 \dots 1.5\}$ | **0.50** | **0.167** | 0.56 | 0.187 | 0.52 | 0.175 | 1.07 | 0.357 |
| $\xi$ | | 0.0075 | $\{0.5 \dots 1.5\}$ | **1.50** | **0.011** | 1.41 | 0.0105 | 1.45 | 0.011 | 0.92 | 0.007 |
| $\chi$ | | 0.15 | $\{0.25 \dots 1.5\}$ | 0.51 | 0.076 | 0.63 | 0.095 | 0.63 | 0.094 | 0.91 | 0.137 |
| $A_{rel,0}$ | | 0.05 | $\{0.5 \dots 2.0\}$ | 1.96 | 0.098 | 1.60 | 0.080 | 1.63 | 0.081 | 0.99 | 0.050 |
| $A_{rel,1}$ | | 0.001 | $\{0.1 \dots 10.0\}$ | 0.51 | 0.0005 | 1.0 | 0.001 | 0.15 | 0.00015 | 1.0 | 0.001 |
| $a_{A_{rel}}$ | | -0.425 | $\{0.0 \dots 1.25\}$ | 1.23 | -0.524 | 1.12 | -0.475 | 1.04 | -0.442 | 1.02 | -0.433 |
| $b_{A_{rel}}$ | $\text{kg}^{-a_{A_{rel}}}$ | 0.007 | $\{0.5 \dots 10.0\}$ | 7.7 | 0.054 | 6.1 | 0.043 | 6.3 | 0.044 | 2.4 | 0.017 |
| $(M_{A_{rel},0})$ | kg | 0.01 | | 32 | 0.32 | 27 | 0.27 | 25 | 0.25 | 8.0 | 0.08 |
| $(M_{A_{rel},1})$ | kg | 97.5 | | 75 | 7.4$\cdot 10^3$ | 28 | 2.7$\cdot 10^3$ | 4280 | 420$\cdot 10^3$ | 6.8 | 660 |
| $B_{rel}$ | $\text{s}^{-1}$ | 5.0 | $\{0.5 \dots 1.5\}$ | 1.46 | 7.3 | 1.28 | 6.4 | 1.26 | 6.3 | 0.97 | 4.9 |
| $\Omega_{,0}$ | | 0.15 | $\{0.25 \dots 1.25\}$ | 0.64 | 0.10 | 0.79 | 0.12 | 0.73 | 0.11 | 0.98 | 0.15 |
| $\Omega_{,1}$ | | 2.0 | $\{0.0 \dots 10.0\}$ | 1.36 | 2.72 | 1.12 | 2.24 | 1.16 | 2.33 | 0.96 | 1.93 |
| $a_\Omega$ | | 0.25 | $\{0.0 \dots 1.25\}$ | 1.20 | 0.299 | 0.82 | 0.204 | 0.88 | 0.220 | 0.88 | 0.219 |
| $b_\Omega$ | $\text{kg}^{-a_\Omega}$ | 0.25 | $\{0.5 \dots 2.0\}$ | 0.80 | 0.199 | 0.97 | 0.242 | 0.91 | 0.228 | 1.07 | 0.267 |
| $(M_{\Omega,0})$ | kg | 0.13 | | 0.66 | 0.085 | 0.23 | 0.030 | 0.27 | 0.035 | 0.51 | 0.065 |
| $(M_{\Omega,1})$ | kg | 4.1$\cdot 10^3$ | | 1.5 | 6.2$\cdot 10^3$ | 13 | 53$\cdot 10^3$ | 9.4 | 39$\cdot 10^3$ | 2.0 | 8.3$\cdot 10^3$ |
| $\cos \varphi_{leg,0}$ | | **0.952** | | | | | | | | | |
| $\cos \varphi_{leg,1}$ | | 0.645 | $\{0.75 \dots 1.05\}$ | 1.02 | 0.655 | 0.79 | 0.508 | 0.76 | 0.490 | 0.95 | 0.610 |
| $a_{\cos \varphi_{leg}}$ | | -0.013 | $\{0.0 \dots 10.0\}$ | 3.9 | -0.051 | 2.7 | -0.035 | 2.8 | -0.036 | 1.37 | -0.018 |
| $b_{\cos \varphi_{leg}}$ | $\text{kg}^{-a_{\cos \varphi_{leg}}}$ | 0.749 | $\{0.75 \dots 1.25\}$ | **0.75** | **0.562** | 0.88 | 0.658 | 0.84 | 0.631 | 0.93 | 0.700 |
| $(M_{\cos \varphi_{leg},0})$ | kg | 1$\cdot 10^{-8}$ | | 3000 | 3$\cdot 10^{-5}$ | 3000 | 3$\cdot 10^{-5}$ | 1000 | 1$\cdot 10^{-5}$ | 3.0 | 3$\cdot 10^{-8}$ |
| $(M_{\cos \varphi_{leg},1})$ | kg | 100$\cdot 10^3$ | | **5.0$\cdot 10^{-7}$** | **50$\cdot 10^{-3}$** | 15$\cdot 10^{-3}$ | 1.5$\cdot 10^3$ | 11$\cdot 10^{-3}$ | 1.1$\cdot 10^3$ | 22$\cdot 10^{-3}$ | 2.2$\cdot 10^3$ |
| $\cos \varphi_{grf}$ | | **0.447** | | | | | | | | | |
| $c_{con}$ | | **0.75** | | | | | | | | | |
| $c_{leg}$ | | 1.0 | $\{0.5 \dots 2.0\}$ | **2.0** | **2.0** | **2.0** | **2.0** | **2.0** | **2.0** | **2.0** | **2.0** |
| speed maximum $\tilde{v}_{max}$ | | 13.9 $\text{ms}^{-1}$ | | 17.6 $\text{ms}^{-1}$ | | 17.0 $\text{ms}^{-1}$ | | 17.3 $\text{ms}^{-1}$ | | 14.7 | |
| at mass $M_{\tilde{v}_{max}}$ | | 16.5 kg | | 40.0 kg | | 39.0 kg | | 37.0 kg | | 93.5 kg | |
| mean deviation factor |  | 1.669 |  | 1.419 |  | 1.426 |  | 1.425 |  | 1.369 |  |
| {min ... max} ratios |  | {0.11 ... 3.19} |  | {0.20 ... 4.00} |  | {0.20 ... 4.16} |  | {0.20 ... 4.20} |  | {0.16 ... 3.60} |  |

We will now outline the transformations that relate, by the geometry linking COM to COP, muscle contraction velocity v_*m*_ to COM (running) speed v, as well as muscle force *F_m_* to the GRF component *F_grf_* in running direction. These relations arise from applying the basic principle of work to the very same geometry and, further, introducing a mechanics-based extension to parametrise the GRF vector 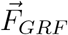 for which the magnitude 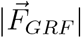 can be estimated by the axial leg force *F_leg_*, that is, the geared muscle force *F_m_*.

### 2.4. EMA Ω and leg projection cos φ_leg_: kinematic constraints linking muscle contraction velocity (v_m_) to axial leg length rate (v_leg_) and running speed (v)

The angle *φ_leg_* (Fig. 1) of the leg axis with respect to the running (forward) direction transforms the axial length rate of change along the leg axis (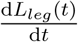; d*t* is the time differential) into the COM velocity in forward direction, that is, (axial) leg velocity 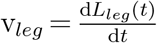 into running speed v and vice versa:

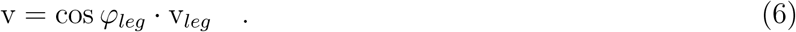

Further on, the EMA Ω (see App. C) transforms muscle contraction velocity 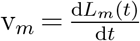 into leg velocity v_*leg*_, and vice versa:

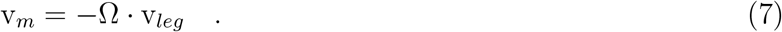

Note the minus sign in Eq. (7), which is due to the shortening (v_*m*_ < 0) of a leg-extending muscle being linked to leg lengthening (v_*leg*_ > 0). From Eqs. (6,7), the transformation between muscle contraction velocity v_*m*_ and running speed v, thus, follows as

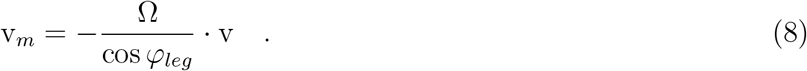

### 2.5. EMA Ω: gearing muscle force (F_m_) into axial leg force (F_leg_)

As a general property of biological locomotion, muscle forces are the sources (or drains) of mechanical work within the animal’s body. A muscle’s force acts with some leverage on all those terrestrial runner’s bones or segments, respectively, that it passes between its origin and insertion. The muscle’s work is directly done at the skeleton. If, at any point in time, work transmission between muscles and those kinetic energy fractions of body segments that are stored in rotations around their respective segmental COMs is negligible—as compared with the instantaneous transmission to and from their linear kinetic energy fractions—, then basic mechanics tells us that *one* necessary requirement is fulfilled for the net work of all muscles being entirely and immediately geared, by the (functional) leg geometry, into the linear COM acceleration of the whole body, which is reflected by the GRF. The *second* (almost) necessary requirement for letting ‘entirely’ exactly apply would be that GRF and functional leg axis align. In this case, the net angular momentum around whole-body COM due to the linear momenta of all functional leg’s segments would not change at that moment. Accordingly, the muscle work would not either change the net rotational energy associated with these segments’ linear momenta, but manage to entirely and exclusively change the linear COM momentum. Misalignment of a *single leg’s* GRF vector and the leg’s axis, therefore, indicates that the instantaneous muscle power generated within this leg may not be entirely transmitted to linear COM acceleration at that point in time. This is nonetheless possible if the instantaneous sum of all acting leg’s GRF vectors does not change the overall angular momentum around whole-body COM.

This work transmission by functional leg gearing (Fig. 1) can be expressed as a torque equality that applies to a joint. The joint torque component (perpendicular to the model plane, Fig. 1)

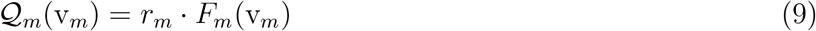

is caused by the force *F_m_* of a single muscle *m* contracting, and exerted through a lever mechanism with opposite signs to both segments that are adjacent to the joint. The muscle’s lever arm *r_m_* with respect to the joint is the perpendicular (shortest) distance between the muscle’s line of action and the joint axis.

There are two basic mechanisms that cause misalignment of GRF vector and leg axis (Fig. 1): (i) inertia forces and torques within the leg, which are particularly significant in leg impact situations [65, 66], and (ii) torques that are applied by proximal muscles [67] originating externally to the anatomical leg, namely, close to the whole-body COM. By contrast, the external torque mechanism due to finite contact areas with the substrate at the leg’s distal end (foot) are fully covered by the COP concept.

If we now assume that inertia forces and torques are insignificant (quasi-static conditions) and, thus, practically negligible at late stance, then *Q_m_* approximately equals the corresponding torque component

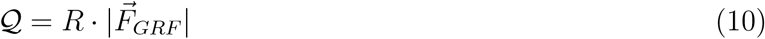

that is exerted on the adjacent segments around this very joint axis due to the GRF, with 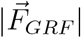 symbolising the magnitude of the GRF vector 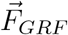, and *R* being the perpendicular (shortest) distance between the joint axis and 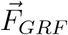 (see blue lines in Fig. 1 and, e.g. [68, fig. 1]). The projection of 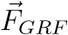 onto the running direction is cos *φ_grf_* (Fig. 1). The projection of 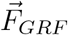 onto the leg axis can be termed the ‘leg force’

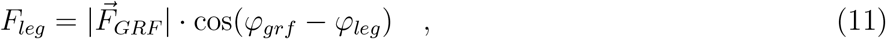

which, in the quasi-static condition, exactly equals 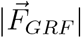 if, in addition, torques by proximal muscles are insignificant (*φ_grf_* = *φ_leg_*). As the deviation of 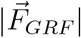 and *F_leg_* is about 30% at maximum (in the smallest animals, see Fig. 6 below: cos(*φ_grf_* − *φ_leg_*) > 0.7), we can set

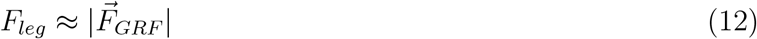

and, thus,

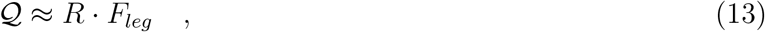

instead of Eq. (10) for simplifying the model.

In Eq. (13), *R* is the (shortest) distance between joint and leg axes (Fig. 1). Both approximate equalities, namely, of 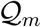 (Eq. (9)) and 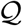 (Eq. (13)) as well as of 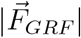 and *F_leg_* (Eq. (12)), imply that the very same geometrically determined EMA 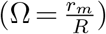 lever mechanism (see App. C) that transmits—via an angular change d*φ_j_* in one joint degree of freedom (DOF) *j*—a muscle’s (v_*m*_) to a corresponding leg’s (v_*leg*_) *length change* (Eq. (7)) likewise dictates the transmission of muscle (*F_m_*) to axial leg *force*

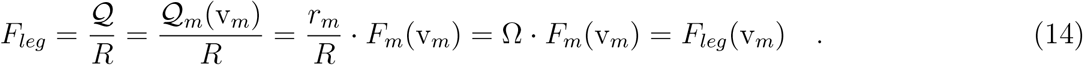

Here, we do not aim to develop a physically and mathematically rigorous generalisation of the EMA concept to the multi-muscle case, let alone to a multi-joint leg structure. The model formulated here relies on (i) the concept of one lumped or crucial muscle for which all equations introduced strictly apply, and (ii) the smooth transferability of geometry and mechanical principles, which together make possible to interpret *F_m_* in Eq. (14) as a generally good approximation of either some single, or net, or mean muscle force value. This value arises from the concentric action of either a key muscle or the concert of a number of muscles and causes the axial driving force *F_leg_*, with an effective Ω value likewise arising from the multiple-muscle, multiple-joint cooperation.

### 2.6. cos φ_grf_: transforming axial leg force (F_leg_) into the forward-driving GRF component (F_grf_)

The projection

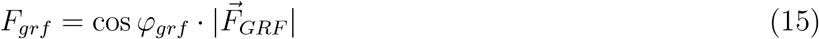

of the GRF onto the running direction, which, according to Eq. (12), can be approximated as

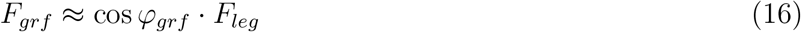

in terms of the axial leg force *F_leg_*, accelerates the COM in this direction (see App. F). With *F_leg_* according to the rightmost equality sign in Eq. (14) being substituted into the approximate equality of Eq. (16), the corresponding transformation between forward-driving GRF component and muscle force writes:

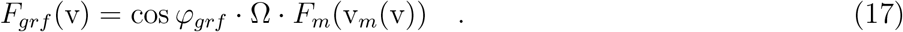

In Eq. (17), we have at once noted by writing functional dependencies that the muscle force is here modelled by a hyperbolic force-velocity relation *F_m_*(v_*m*_) (Eq. (E.1)) and the muscle contraction velocity v_*m*_ directly relates to running speed v (Eq. (8)).

### 2.7. Premature work termination due to finite stance duration and muscle inertia

In this section, we explain a physical constraint that can impose lower values of maximum running speed than those predicted (Eq. (3)) from the force equilibrium (Eq. (2)) between the geared muscle force and air drag.

When an animal is running, at some instant during a legs’ stance phase, which occurs around MS or ISOMS, respectively, the leg is at its minimum length [28, 29, 30]. In late stance (at LO or TOPSS, respectively), the leg has become longer (extended) again. Several muscles crossing leg joints must have shortened accordingly, starting from the isometric condition around MS and then actively increasing their shortening velocities during concentric contractions. These muscles are, therefore, ‘leg extensors’. While most muscle-tendon complexes in the leg are approximately isometric around MS, the muscle fibres of the extensors are very likely still in eccentric [35] conditions at MS and switch to the concentric mode later—towards TOPSS—than the whole complex. In any case, we have suggested so far in this paper, that the hyperbolic force-velocity relation (Eq. (E.1) or Eq. (I.1), respectively) of a lumped or crucial leg extensor muscle is the very muscle-mechanical characteristic that determines leg work during late stance (towards TOPSS).

The very same hyperbolic force-velocity relation that crucially determines the prediction (Eq. (3)) of maximum running speed from the force equilibrium (Eq. (2)) likewise enters into a constraint that can impose a running speed below this force-based prediction. To explain this, we first recall that the outlined boundary conditions of the leg during running require an acceleration of leg masses in space towards TOPSS. At the same time, the mass of any working extensor muscle is accelerated along the muscle’s line of action. Second, because the hyperbolic force-velocity relation represents active muscle fibres contracting against *external loads*, the structural properties that underlie this characteristic are, likewise, inherent to the dynamics of driving the muscle material itself through velocity changes (acceleration) while the *muscle’s inertia* (*mass*; see Eq. (J.1)) *counteracts*. Based hereupon, we have calculated, in App. I, the mechanical dynamics of a muscle accelerating its own mass distribution along its line of action in response to changing contraction force demands. The main result of App. I is the time *τ_m_* (Eq. (I.19)) required for a muscle to settle from an initially generated force value to a newly demanded force value.

The functional consequence of muscle inertia, expressed in terms of *τ_m_*, for running speed v is that the estimated time *T_tra_* (Eq. (J.4)) of the whole-body COM to move at v from ISOMS to TOPSS, which we assume to be equal to nearly half of the stance duration *T_con_* (see Eq. (J.3)), can become *shorter* than the time *τ_m_* that the inert muscle requires to increase its contraction velocity v_*m*_ from ISOMS to TOPSS, i.e., to settle to the TOPSS force level. For any body mass *M*, we, thus, ask for the critical condition

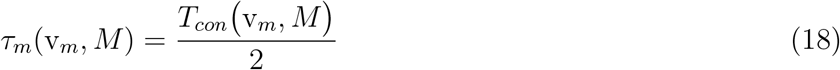

of equality of both durations. The particular muscle contraction velocity

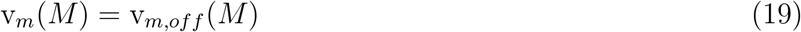

that fulfils Eq. (18) is the answer to this question. In App. J, more detailed explanations of the derivation of condition Eq. (18) are given, as are explanations of how it is solved. Most notably, Eq. (18) is a cubic equation (Eq. (G.2)) like the force equilibrium Eq. (2) (i.e., Eq. (G.1) explicitly), though, here in terms of *z* = v_*m*_ rather than *z* = v. Accordingly, Eq. (18) can, like Eq. (2) (solution: (Eq. (H.4))), be solved (Eq. (H.6)) in closed form.

Through the inversion of Eq. (8), we can finally calculate the running speed

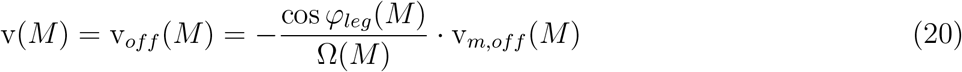

that corresponds to the found (Eq. (19)) contraction velocity v_*m,off*_ (*M*). The speed v_*off*_ is a secondary model prediction of maximum running speed v_*max*_—an alternative to v_*feq*_ fulfilling Eq. (3). In particular, it turns out that v_*off*_(*M*) (Eq. (20)) is predicted to be lower than v_*feq*_(*M*) (Eq. (3)) above a critical body mass 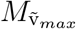. Therefore, we term the condition Eq. (18) the *duration constraint*. According to our calculation, animals larger than represented by the critical body mass 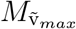 cannot run as fast as the force equilibrium (Eq. (2)) would allow (v_*feq*_(*M*)), because the duration constraint, which in terms of *T_con_* represents time-critical mechanisms *other* than forward drive by muscles, enforces LO before having attained TOPSS (Eq. (2)) and, with it, induces the premature termination of work release by the leg, thus, of whole-body COM acceleration.

## 3. Results

### 3.1. Maximum running speed limited by premature work termination

The minimum value

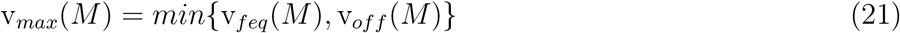

of the two solutions Eq. (3) (i.e., Eq. (H.4)) and Eq. (20) (i.e., Eq. (H.7) with Eq. (H.6)) for running speed v as calculated from Eq. (2) (i.e., specifically Eq. (G.1)) and Eq. (18), respectively, is our model prediction of the maximum running speed for the whole range of the body masses of legged animals found on earth to date: the legged speed allometry v_*max*_(*M*). The overall characteristic calculated this way is meant to cover the gross scaling tendencies of maximum running speed: a mechanistic explanation for the allometric ‘main road’. For an examination of the body designs of specific species by means of our model, we refer to the discussion in Sect. 4.1.

In Fig. 2, we have plotted this characteristic as arising from an initial guess of all parameter values taken from the literature (column three in Table 2): v_*feq*_(*M*) is the very thick light blue line and v_*off*_(*M*) is the very thick green line partly overlaid by the yellow one, while the overall predicted legged speed allometry v_*max*_(*M*) according to Eq. (21) is the thick orange line. For comparison, the data fit by Hirt et al. [1] is plotted using a thin red line with circles. The other solution variants plotted in Fig. 2 are explained further below. The solution v_*max*_(*M*) changes from v_*feq*_(*M*) to v_*off*_ (*M*) with a kink—where thus v_*max*_ = v_*feq*_ = v_*off*_ is fulfilled—in the thick orange line at a (critical) mass 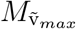 of 16.5 kg and a corresponding speed 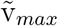 of 13.9 m s^−1^. Generally, the kink at 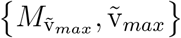 is our prediction of the overall maximum in the legged speed allometry. The second kink back to an upward tendency in v_*max*_(*M*) at *M* ≈ 4 · 10^3^ kg is due to the only physical constraint (Eq. (18)) considered in our present running model in addition to Eq. (2), in combination with a plain assumption for Ω(*M*) at sizes where no EMA data are currently available (further explanations in App. L). Therefore, any v_*max*_(*M*) prediction based on our model for animals bigger than indicated by this upward kink is unreliable.

Note, first, that this v_*max*_(*M*) prediction with its specific kink values is based on our initial parameter guess. Our v_*max*_(*M*) findings are only moderately changed by parameter optimisation (see below and Table 2). Further, note that our model predicts, in general, that any animal larger than indicated by the kink at 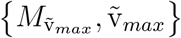 achieves (only) a maximum running speed determined by the premature termination of muscle work (for more details, see App. J). By ‘premature’, we mean that within (about half of) the leg’s finite stance duration *T_con_*, which is likely to be governed by vertical COM repulsion dynamics (e.g., [26]) and/or limited stance lengths, the muscle is too inert (as quantified by Eq. (18) in terms of its settle time *τ_m_*) to accelerate to the very contraction speed that would be possible in accordance with the force equilibrium Eq. (2).

The violet line in Fig. 3 shows the estimated halved stance duration 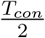 (Eq. (J.7)) as a function of body mass *M*, while assuming that the animal runs at speed v = v_*feq*_(*M*) as predicted by the specific formulation Eq. (G.1) of the force equilibrium Eq. (2). The light blue and brown lines in Fig. 3 show two specific settle times (Eq. (I.19)) of the muscle: *τ_m,max_*(*M*) and *τ_m,min_*(*M*), which arethe two boundaries of the durations required for contractions to settle tozero-load concentric v_*m*_(*M*) = −*V_m,max_*(*M*) orisometric v_*m*_(*M*) = 0 conditions,respectively. Generally, settle times *τ_m_*(v_*m*_(v, *M*), *M*) increase steeply with *M* as compared (in Eq. (18)) with halved stance duration 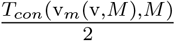. The steep increase in *τ_m_*(v_*m*_(v, *M*), *M*) (Eq. (I.19)) is due *primarily* to muscle mass *M_m_* increasing linearly with *M* (exponent: 1; Eq. (J.1) with inserting Eq. (J.2),Eqs. (E.3,E.4), and Eq. (5)), and *secondarily* to TOPSS (Eq. (G.1)) shifting to operating points on the force-velocity relation with slopes 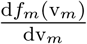 that probably decrease with body mass. The net *M*-exponent of *τ_m_*(v_*m*_(v, *M*), *M*) is about 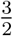 for the slowest predicted concentric settling (at *F_m_* = 0: *τ_m,max_*(*M*), the light blue line in Fig. 3) and about 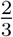 for the fastest concentric settling predicted to be possible (around the isometric state *F_m_* = *F_max_*: *τ_m,min_*(*M*), the light brown line in Fig. 3).

**Figure 3:**
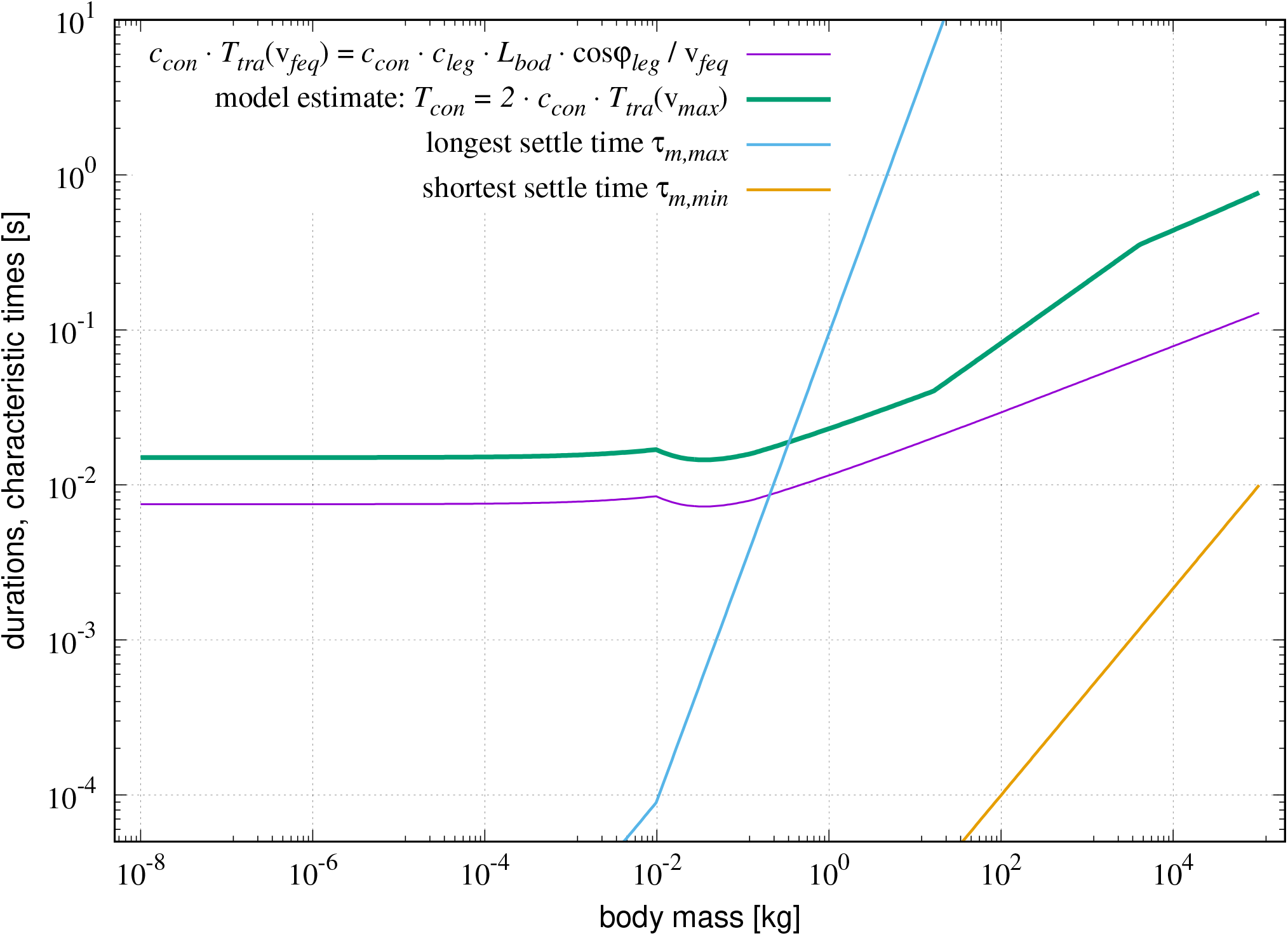
Model predictions of characteristic times vs. body mass *M*. (i), violet line: Model-estimation of half the stance duration 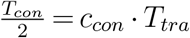 (Eq. (J.3)) at (maximum) speed v_*feq*_(*M*) being *model-calculated by solving the force equilibrium* (Eq. (2)), with an empirical factor *c_con_* = 0.75 (see Table 2 and text below Eq. (J.8)) that approximates stance durations in both mite running and human sprinting. (ii), green line: Corresponding stance duration estimate *T_con_* = 2 · *c_con_* · *T_tra_* at the model-predicted maximum speed v_*max*_(*M*) (Eq. (21)), with here v_*max*_(*M*) = v_*off*_ (*M*) < v_*feq*_(*M*) for *M* > 16.5 kg *due to the duration constraint* (Eq. (18); see Fig. 2): Along the ordinate, the green line is two times the violet one for *M* < 16.5 kg, with the v_*max*_-restriction by v_*off*_ (*M*) superposed for *M* > 16.5 kg (two additional kinks: see below); *T_con_*(*M*) well matches data [30, fig. 5] measured in the range *M* = 10^−1^…10^2^ kg. (iii), light blue line: According to the muscle mass being accelerated by a muscle force modelled as Hill’s hyperbolic force-velocity relation (Eq. (I.1), the time *τ_m,max_* (Eq. (I.19) for settling of muscular contraction velocity v_*m*_ → −*V_m,max_* after a change to zero muscle force demand *F_m_* = 0 (slowest settling). (iv), light brown line: Settle time *τ_m,min_* like in (iii), however, for a change in force demand to nearby the isometric force *F_m_* = *F_max_*, i.e., v_*m*_ 0 (fastest settling). Thus, lines (iii) and (iv) depict the extreme values ((iii): the longest, (iv): the shortest) of the settle time that our model muscle needs, as a function of body size, to contract from ISOMS to TOPSS. At *any* body size, the force equilibrium (Eq. (G.1: TOPSS) can be fulfilled in our model by a muscle contracting concentrically at the force 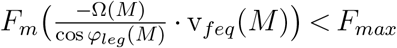. *Below* the critical body mass 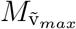 (bottom of Table 2; here: note the kink in green *T_con_* line at 16.5 kg) to which the overall running speed maximum 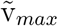 corresponds, the settle time *τ_m_*(*M*) that corresponds to the operating point at 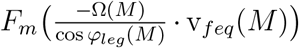 is *shorter* than half the stance duration 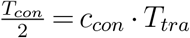 (Eq. (J.8), with then muscle inertia predictably *not interfering with* maximum running speed. Exactly the critical point 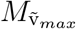, 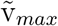, the condition 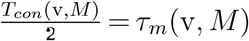 is fulfilled. In our model, the latter equality (Eq. (18)) is then used to calculate the solutions v_*max*_(*M*) = v_*off*_ (*M*) for all 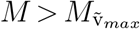, i.e., where muscle inertia interferes.

Somewhere between the light blue and brown lines in Fig. 3 (i.e., in the region *τ_m,min_*(*M*) < *τ_m_*(v_*m*_, *M*) < *τ_m,max_*(*M*)), at body masses higher than a (critical) value 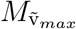, the v_*off*_ (*M*) solution actually undergoes the solution v_*feq*_(*M*) from the force equilibrium (Eq. (G.1)); see, again, Fig. 2. Here, the muscle force values that correspond to v_*off*_ (*M*) are always positive. For animals larger than this critical size, v_*max*_(*M*) is, thus, determined by v_*off*_ (*M*) (see Eq. (21). The predicted kink (overall maximum) in the legged speed allometry v_*max*_(*M*) (the thick orange line in Fig. 2) at 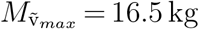 is also discernible in the course of the predicted stance duration *T_con_*(*M*) (the green line in Fig. 3).

### 3.2. Synopsis of parameter optimisation

In Table 2, a compilation of all 26 non-vanishing model parameters (column one) and five sets of their values are given: the set containing our initial guess from the literature (column three: ‘initial guess’ *λ_l,ini_*) as well as four sets that were determined by minimising the deviation of the v_*max*_(*M*) solution from the experimental data cloud provided by Hirt et al. [1]. For this, we employed the non-linear optimisation (least-square minimisation), trust-region-reflective algorithm *lsqnonlin* implemented in MATLAB (version R2019a, The MathWorks, Natick, MA, USA). The parameter values of the four optimised v_*max*_(*M*) calculations are reported in columns six, eight, ten, and twelve of Table 2.

The physical parameters listed in the four top rows of Table 2, which model air drag, were generally assigned fixed values to, as were to another five model parameters: *ρ_bod_*, *σ_max_*, cos *φ_leg,_*_0_, cos *φ_grf_*, and *c_con_* (their values printed in bold font in Table 2). Thus, seventeen parameter values were available to the optimisation algorithm for variation. The allowed ranges of variation are given in column four in terms of the minimum and maximum factors to the initial guess value. In the event that the algorithm assigned the final (optimal) parameter value to a boundary value, the optimal value listed in Table 2 is printed in bold font. Parameter values given in parentheses (mass values) are given for transparency, though they are simply transformed representations of the corresponding set of four parameters *a_pn_*, *b_pn_*, *p_n,_*_0_, and *p_n,_*_1_ that fix a power law *p_n_*(*M*) of a size-dependency in the model.

In our main optimisation case (‘model opt’: columns seven and eight), maximum isometric muscle stress was fixed at the (default) value *σ_max_* = 2.5 · 10^5^ N m^−2^ as an initial guess (column three). Another two optimisations, with −20% (columns five and six) and +20% (columns nine and ten) of this default *σ_max_* value, were run. Again, using the default *σ_max_*, the fourth optimisation case (‘mite×100’: columns eleven and twelve) emulated having the same number of measured data points available at very small sizes, i.e., in the spider and insect range, as for vertebrates: the four data points for the smallest animals (mites) were simply weighted a hundredfold. In this hypothesised case, the body size at which the size-scaling of ‘aligning the joints’ (Ω(*M*): *M*_Ω,0_ = 65 g) and ‘introducing tendons’ (*A_rel_*(*M*): 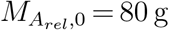) set in, would be located close to each other. In all other optimised cases (‘model opt’, ‘ ±20%’), ‘introducing tendons’ is calculated to set in at higher values of about M = 300 g. These three optimisations (c.f. Fig. 4) distinguished only by *σ_max_* variation calculate, on the allometric ‘main road’, an overall running speed maximum of 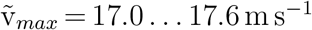 occurring at 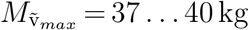. When one more highly weights (case ‘mite×100’) small animals without tendons, the overall speed maximum would be calculated to occur in larger animals 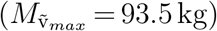 at a lower speed value 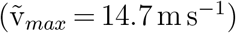.

**Figure 4:**
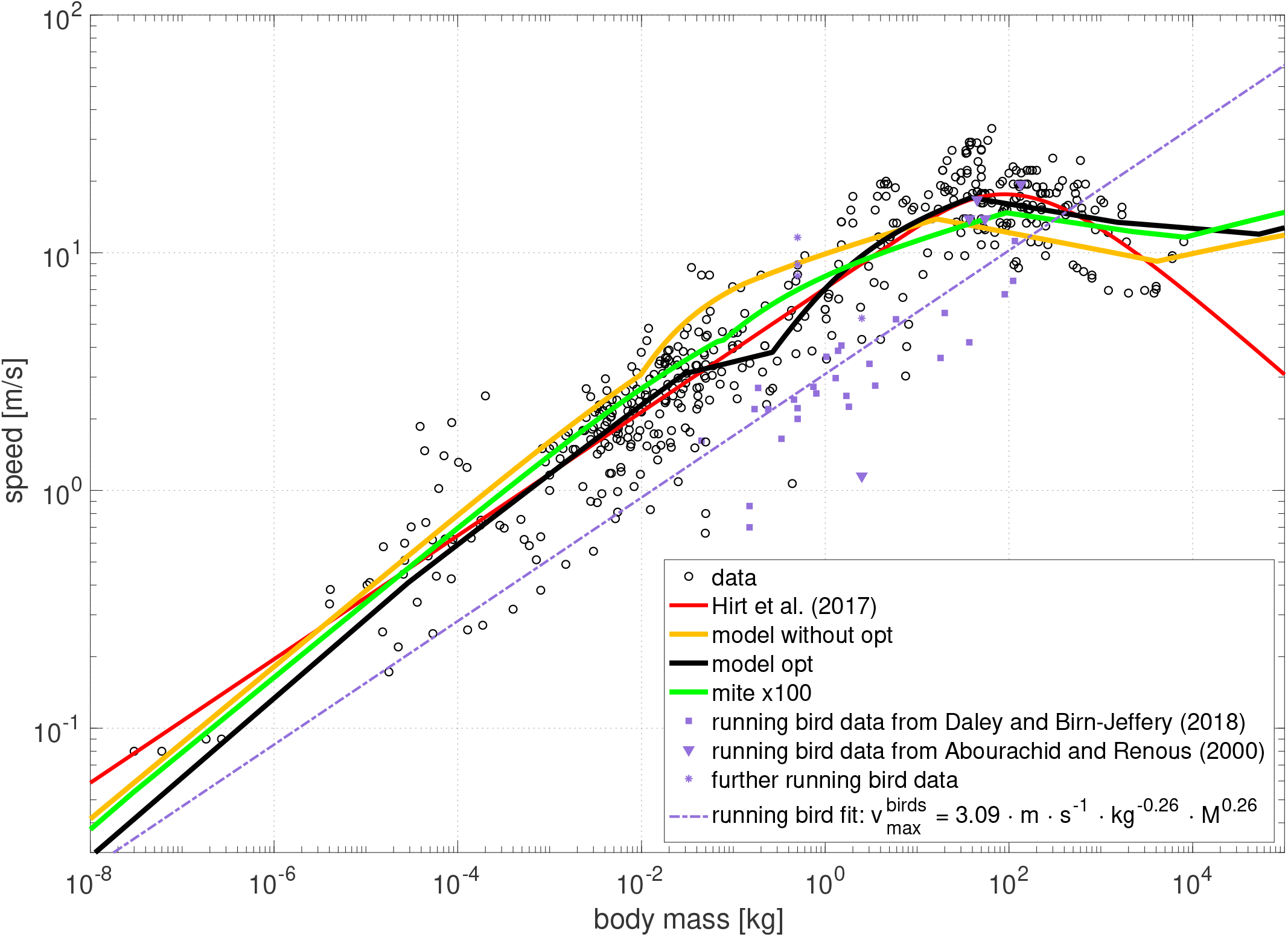
Experimental data, empirical function fit thereto, and three model solutions of the maximum running speed v_*max*_ vs. body mass *M*. Plotted are the compilation of experimental data points (black circles: ‘data’) provided by [1], their fitting function (red line: ‘Hirt et al. (2017)’), as well as three of our model solutions: The one (orange line: ‘model without opt’, the same as the thick orange line in Fig. 2) calculated with initial guesses of parameter values extracted from literature (third column in Table 2; see also Fig. 2), the one (black line: ‘model opt’) with parameters minimising the residuum with respect to the experimental data points while assuming *σ_max_* = 2.5 · 10^5^ N m^−2^ (eighth column in Table 2), and the one (green line: ‘mite x 100’) with the same *σ_max_* value, but each of the four data points for the smallest animals (mites) weighted a hundredfold (twelfth column in Table 2). The coordinates of the overall maximum in running speed 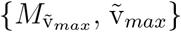 are: {87.9 kg, 17.6 m s^−1^} for ‘Hirt et al.’, {16.5 kg, 13.9 m s^−1^} for ‘model without opt’, {39.0 kg, 17.0 m s^−1^} for ‘model opt’, and {93.5 kg, 14.7 m s^−1^} for ‘mite 100’. The four experimental data points at the lowest body masses (from three mite species of which two are part of the data set by [1] providing three points) have been made comparable with conditions found in mammals, i.e., body temperatures of about 37° C. Further explanations of the data plotted are given in App. M.

For our default value of maximum muscle stress (*σ_max_* = 2.5·10^5^ N m^−2^) and if it were fixed to a 20% higher value, the optimised parameter values settled generally well into the boundaries that we had assessed as being somehow reasonable, except for the parameter *c_leg_* determining (functional: Fig. 1) leg length *L_leg_* (Eq. (J.6)). However, leg length is *solely* significant in the duration constraint (Eq. (18) with Eq. (J.8): *c_leg_*-dependent) and *not* for fulfilling the force equilibrium (Eq. (G.1): *c_leg_*-independent). Therefore, the model solution v_*feq*_(*M*) for body sizes below the overall speed maximum does *not* depend on leg and stance length. However, due to the duration constraint (Eq. (18)), the condition of ‘premature work termination’ is shifted to higher body sizes if the leg and, with it, the stance length become longer. This is because increasing leg length (Eq. (J.6)) increases stance duration (Eq. (J.7)) by means of increased stance length (Eq. (J.5)) for a given running speed v. With this, any muscle is given more time 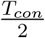(*M*) (Eq. (J.7)) for accelerating itself within *τ_m_*(*M*) (Eq. (I.19)) to the high contraction velocity at TOPSS (Eq. (2)).

In many cases, our model assumes structural properties for which no experimental data are available. Thus, often, quite wide boundaries were allowed. In the case of leg length, the optimised model results— particularly, the location of the overall speed maximum (the kink)—are, thus, based on the assumption that the functional leg is twice as long as the characteristic (trunk) length *L_bod_*. However, this may be a result of optimising on the basis of a measured data cloud that is scattered, around the location of the overall speed maximum, across a wide range of sizes (1…500 kg: Fig. 4) and body designs. The value *c_leg_* = 2 may apply to a cheetah, though not to a human or rhino. Also note that in the model presented here, just the product *c_con_ · c_leg_* counts (Eq. (J.8)); geometric and temporal properties are indistinguishable as, currently, no mutual coupling with a mechanical model of repulsion dynamics is taken into account. Thus, to avoid redundancy in optimisations while confining model elaboration, we left the parameter *c_con_*, which represents mechanisms that govern stance duration other than by variation of leg length, always fixed to the rough initial guess value.

The ratio 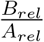 of the two Hill-parameters represents the multiple of optimal fibre length *L_opt_* per second (i.e., the strain rate) at which the muscle contracts with zero load (see *V_m,max_*: Eq. (I.5)). In our model, the ratio 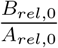 represents the maximum strain rate of *sole fibre material* (muscle belly), as this number applies to animals too small to be equipped with tendons. The numbers are (Table 2) 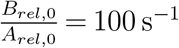 for the ‘initial guess’, 98 s^−1^ for the ‘mite×100’ case, 75 s^−1^ and 78 s^−1^ for the ‘±20%’ cases, and 80 s^−1^ for the ‘model opt’ case. These values are much higher than those measured in steady-state contractions. *Yet,* the model muscle, evaluated at TOPSS, definitely represents a *non-steady*, concentric contraction state: a snapshot of running dynamics to which the muscle (particularly the fibre mass) has accelerated itself, starting from ISOMS. Steady-state values have been measured, for example, in small mammalian [69] and spider [70] muscle fibres. Extrapolating—for comparison with all other species—their 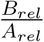 ratios to a body temperature of 37° C by a *Q*_10_-factor of 1.7 [71, p. 147] that applies to 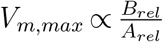, we find steady-state values of 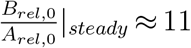 in both species. Now, the non-steady to steady speed ratio is about eight [36]; and TOPSS is, muscle-wise, a non-steady state. First, it can, thus, be concluded that our optimised 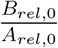 ratios (78…98) are well matched by extrapolated experimental data for non-steady state contractions 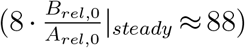. Second, it can be noted that the lumped serial elasticity of a piglet’s (*M* = 1.5 kg) main ankle extensor muscles, which accelerated—in quick-release experiments—various masses that had been applied as external loads, amplified the *V_m,max_* value of the whole muscle(-tendon complex) as compared with sole fibre material (the “contractile element” in [40]) by a factor of 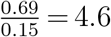 [40, fig. 5]. Based on this observation, an increasing significance of tendons in terms of an increase in the ratio 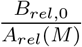 is modelled by an assumed decrease of *A_rel_*(*M*) with body mass *M*, which starts at 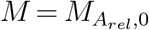 (see, e.g, Fig. 5 capturing our ‘initial guess’ parameter set). To summarise, the 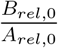 values found by optimisation, as well as our initial guess, and, likewise, the simple allometric scaling ansatz for *A_rel_*(*M*), are echoed by experimental data.

**Figure 5:**
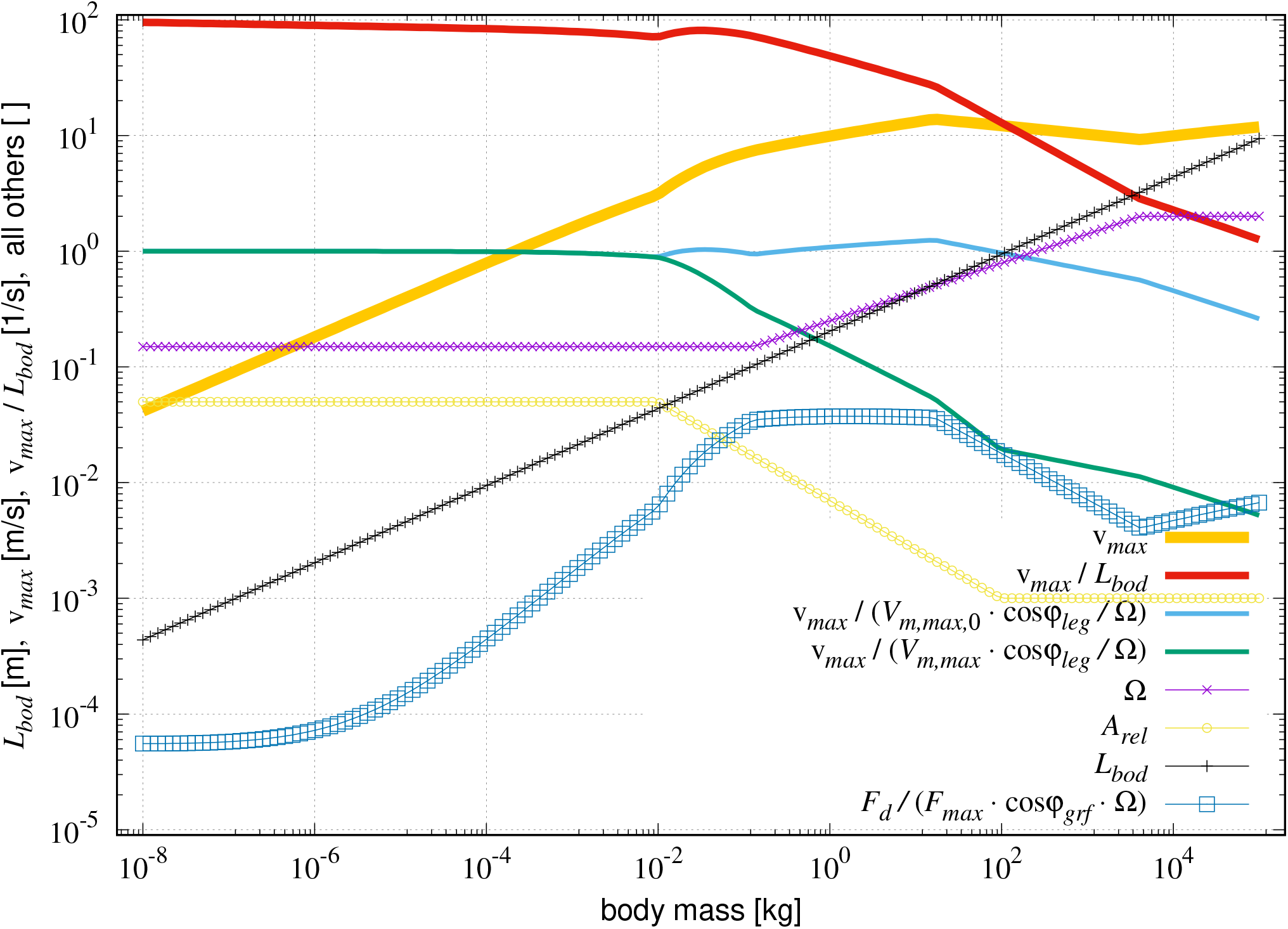
Diverse model input and output vs. body mass *M*. (i; output), thick orange line: Maximum running speed v_*max*_ (Eq. (21)). (ii; output), thick red line: Maximum running speed v_*max*_ normalised to characteristic body (trunk) length *L_bod_*. (iii; output), light blue line: v_*max*_ normalised to the running speed that corresponds to our model estimate of the *muscle fibres’* maximum concentric contraction velocity *V_m,max,_*_0_ (Eq. (I.5) with constant *A_rel,_*_0_ termed ‘without tendon’), which is transformed by EMA Ω to axial leg lengthening rate and then projected by cos *φ_leg_* onto running direction (Eq. (8), Fig. 1). (iv; output), green line: Same as in (iii) except v_*max*_ normalised to the running speed that corresponds to the transformed and projected (Eq. (8)) *whole muscle’s* (including potential tendon: *A_rel_*(*M*)) maximum concentric contraction velocity *V_m,max_* (Eq. (I.5)). (v; input), violet line with crosses: Ω (Eq. (C.1), App. D.2). (vi; input), yellow line with circles: ‘Hill parameter’ *A_rel_* (Eq. (I.6), App. D.1). (vii; input), thin black line with plus signs: Characteristic length *L_bod_* (Eq. (5). (viii; output), dark blue line with squares: Drag force *F_d_* (Eq. (B.1)) normalised to the isometric muscle force *F_max_* (Eq. (E.2), which is transformed by Ω to the corresponding axial leg force *F_leg_* (Eq. (14)) and then projected onto running direction by cos *φ_grf_*, i.e., normalised to the maximally possible forward-driving GRF component (the value for zero speed).

### 3.3. The muscle running against both itself and the air

According to our model, running speed increases generally with the rate of length change v_*m*_ (velocity) of contracting muscle fibres: Inverting Eq. (8) provides the relation 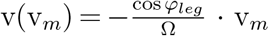. Thus, it is not surprising that the main factor in the increase in maximum running speed with body size on the allometric ‘main road’ v_*max*_(*M*) (Fig. 2: thick orange line) is the muscle’s maximum concentric contraction velocity *V_m,max_* increasing linearly with muscle belly (fibre) length *L_opt_* (right equality sign in Eq. (I.5)); then again, *L_opt_* scales linearly with characteristic body length *L_bod_*(*M*) (Eq. (E.4)); therefore, v_*max*_(*M*) eventually scales with the third root of body mass *M* (exponent: 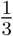; Eq. (5)). Such linear (i.e., geometrical) scaling of v_*max*_ with *L_bod_*(*M*) can be observed in its pure form in small animals, as these are, first of all, practically not hindered by air drag. Accordingly, in Fig. 2, the thin ‘no drag’ lines are within the thick orange line below about *M* = 10 g. Here, air drag *F_d_* is less than 1% of the maximally possible driving GRF (Fig. 5: blue line with squares). Second, elasticity in series to the fibres (e.g., tendons) very likely does not play a significant role in small animals. Accordingly, in our model with optimised parameter values, *A_rel_*(*M*) remains constant below 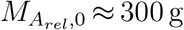 (Table 2; initial guess: 10 g, see Fig. 5).

Adding elastic material, e.g., tendons, in series to muscle fibres has the crucial functional effect of boosting the value of a muscle’s maximum concentric contraction velocity *V_m,max_* [40, fig. 5]. By allowing a change in the value of the ‘Hill parameter’ *A_rel_* (see Eq. (I.5)) as a function of *M* (see again Fig. 5), we have mathematically implemented the option to find such a boost effect in our model. In fact, we found, in our optimisation calculations, a reduction of *A_rel_*(*M*) with increasing body size (Table 2: 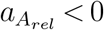), which deviates only in numerical detail from our initial guess. Accordingly, nature has evidently made use of this strategy to boost *V_m,max_*(*M*) beyond its geometrical fibre scaling with *L_bod_*(*M*): Our optimised model calculations, which find the body size of *A_rel_*(*M*) change onset at 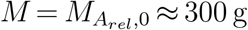, confirm a statement in [72] that, due to Young’s modulus of tendon material, the storage of elastic energy in tendons has become mechanically significant only in the running of animals larger than a kangaroo rat [73], i.e., for *M* ≳ 100 g.

The two main scaling tendencies, *L_bod_*(*M*) and *A_rel_*(*M*), that induce an increase in v_*max*_(*M*) with body size are countered by three major mechanisms:

i. The coefficient of damping 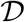 (Eq. (B.2)) for air drag *F_d_* (Eq. (B.1)) increases with the body’s cross-sectional area (CSA) Æ_*bod*_ (Eq. (B.3)), which itself increases with body mass (exponent: 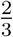). In the first instance, this seems to be just about compensated for by the driving muscle force increasing with the same exponent, along with the muscle’s (fibre material) CSA Æ_*m*_ (Eq. (E.3)). *Moreover*, air drag increases with speed. Figure 5 demonstrates (see the dark blue line with squares) that air drag *F_d_*, when normalised to the maximally possible driving GRF, indeed increases with body size up to the overall maximum in running speed, and then decreases again due to decreasing speed.
ii. Second, any increase in v_*max*_(*M*) with size by v_*m*_ increasing with *V_m,max_*(*M*) may be countered by the erection of legs (more upright) likewise increasing: See again 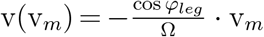 (Eq. (8)) and note that cos *φ_leg_*(*M*) decreases with size (Fig. 6). However, such a decrease in cos *φ_leg_*(*M*) is assumed to only subtly occur across the entire body size range (low magnitude of the exponent: −0.02 0.05; see Table 2). Note: There seems to be term-wise confusion in literature. We use the term *erection* to express the notion that a leg is brought to a more *upright* position, i.e., to make it more aligned with either gravitational acceleration or the normal vector of the leg’s contact surface. This is different from *aligning* a leg, which means increasing EMA (Ω), i.e., aligning the joints to the leg axis. In two key papers [58, 74] about EMA, the terming was the same as what we adopted here. Unfortunately, the term “erect” in the meaning of “aligned” was then used in a later review [75].
iii. Third, the effect of the *V_m,max_*(*M*) increase with size on v_*max*_(*M*) is significantly countered (in particular, compare the thin black and violet lines in Fig. 2) by an increasing alignment of joints to the leg axis: an increase in Ω(*M*). Such an Ω(*M*) increase has been found to occur in mammals with body masses of above approximately 100 g [58]. Our model calculations almost perfectly confirm this finding, as values of *M*_Ω,0_ in the range of 30…85 g (Table 2) are calculated in all optimised cases. Furthermore, optimised values in the range 0.20…0.30 for the exponent *a*_Ω_ match the experimentally found values [74] (initial guess: 0.25) similarly well. Notably, the v_*max*_-countering Ω increase starts approximately at body sizes for which tendons also become significant.

**Figure 6:**
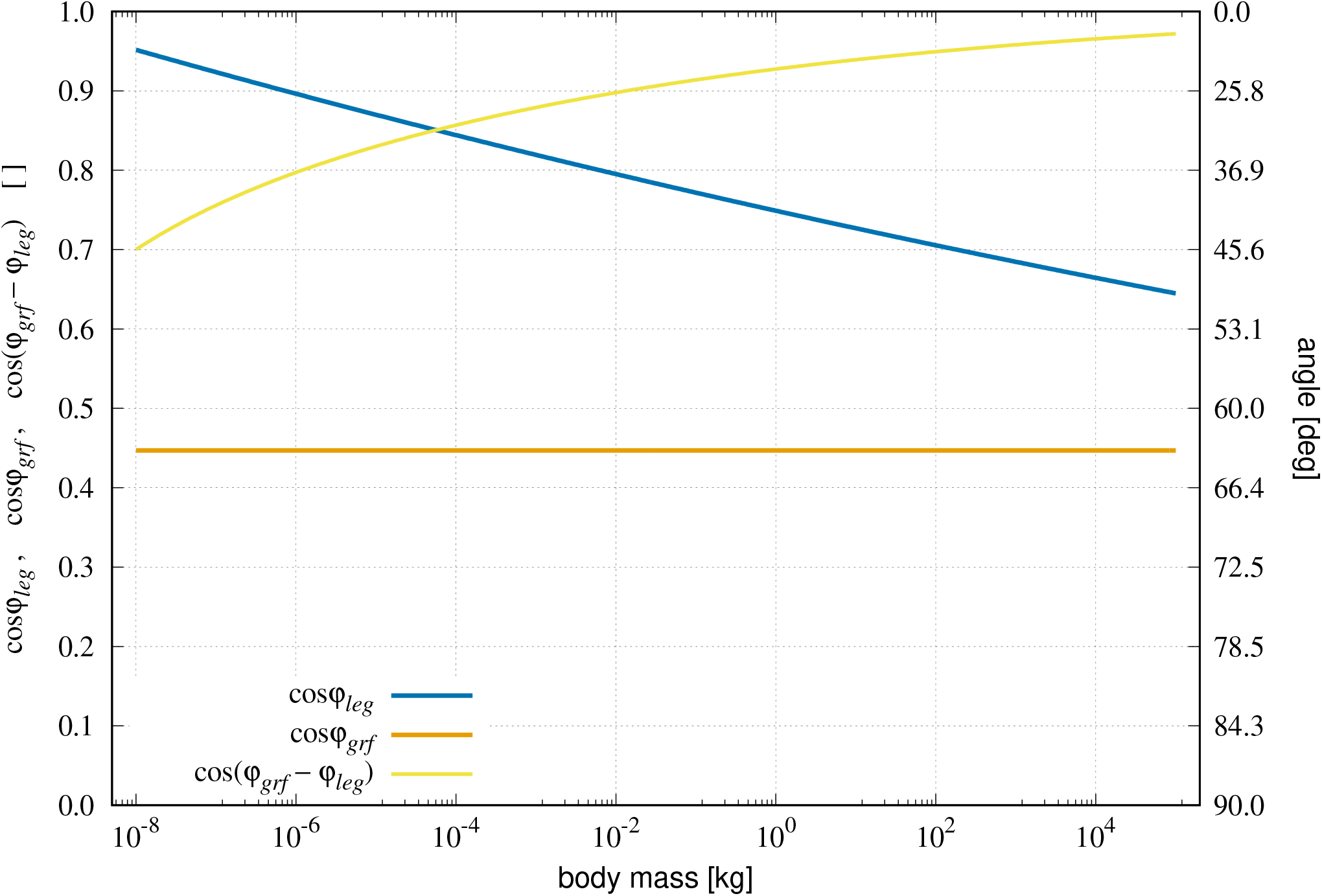
Model input: Leg and GRF angles (Fig. 1) vs. body mass *M*. The ordinate scale on the right side indicates the degree value of the angle *ϕ* that corresponds to cos *ϕ* given on the left side. (i): Model parameter cos *φ_leg_* (Eq. (6), App. D.3) representing the leg angle, i.e., the projection of the leg axis onto the running direction. (ii): Model parameter cos *φ_grf_* (Eq. (F.1), App. D.3) representing the GRF angle, i.e., the projection of the GRF vector onto the running direction. (iii): Their difference represented by cos(*φ_grf_* − *φ_leg_*), with the leg axis and the GRF vector being aligned for cos(*φ_grf_* − *φ_leg_*) = 1.

The predicted allometry of air drag and flow is conflated into Fig. 7. The smallest animals (mites with *M* ≈ 0.01 mg) run at a Reynolds number of *Re* ≈ 1, i.e., surrounded by stationary laminar airflow, well in the Stokes flow regime. Then, up to a body mass of about 1 mg, an animal experiences periodically laminar flow (*Re* ≈ 1…40) with regular, stationary vortices attached behind its body. Above *M* ≈ 1 mg, the Newtonian flow regime replaces the Stokes regime. First, the regular vortices of size of about the body dimensions detach into a ‘Kármán wake’. Then, above about 10 mg…100 mg, with *Re* ≈ 100…400, the regular vortices are increasingly accompanied and superposed by small-scale turbulences with increasing *Re* values. The drag coefficient is approximately constant (*C_d_ ≈ C*_∞_) in the Newtonian flow regime, i.e., at all body sizes above 10 mg…100 mg. When *Re* = *Re_c_* ≈ 3…5 · 10^5^ is reached, i.e., at *M* = 5…10 kg, the wake behind the body has narrowed to a fluidised bed that consists solely of small-scale turbulences.

**Figure 7:**
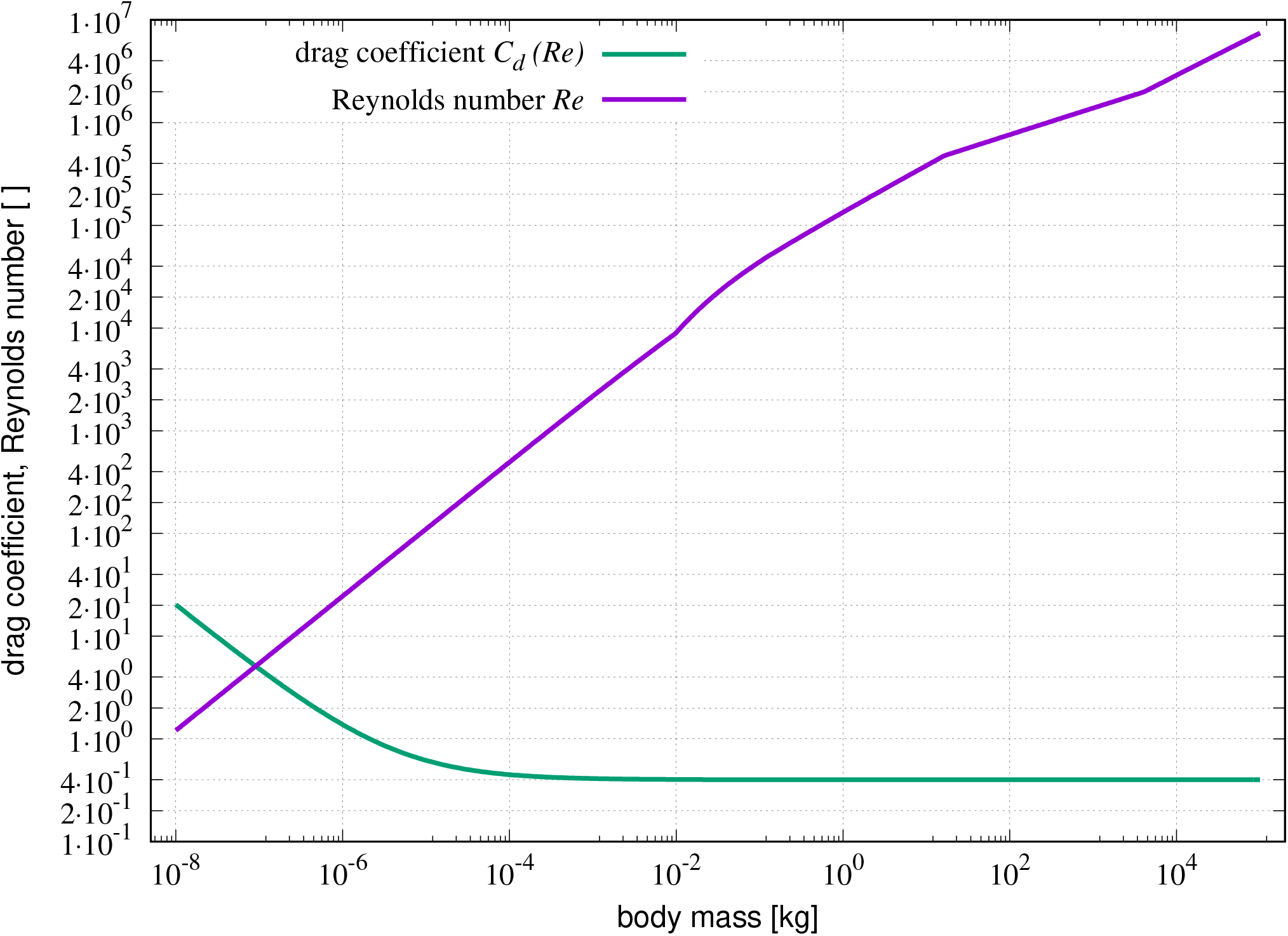
Model predictions of air drag variables vs. body mass *M*. A very brief abstract of air drag physics is provided in App. B. (i): Drag coefficient *C_d_*(*Re*) (Eq. (B.4)) and (ii): Reynolds number *Re* (Eq. (B.5)) at v = v_*max*_. Above approximately *M* = 10^−6^ kg, air drag starts to change from the Stokes towards the Newtonian flow regime. The critical *Re* value *Re_c_* ≈ 3…5 · 10^5^ is reached at *M* ≈ 10 kg.

### 3.4. Sensitivity analysis

There is high variance in the measured data [1, fig. 2c] of maximum running speed v_*max*_ across the whole mass range. For example, at *M* ≈ 50 kg, the ratio between approximately v_*max*_ = 32 m s^−1^ for cheetahs (*Acinonyx jubatus*)—measured 29 m s^−1^ [76], maximum very likely higher [77]—and measured values from heavier top sprinters [78] downscaled to about 10 m s^−1^ for 50 kg humans (*Homo sapiens*) is about three. At *M* ≈ 0.5 kg, the ratio between v_*max*_ = 8.0 m s^−1^ [79] for roadrunners (*Geococcyx californianus*) and v_*max*_ = 1.05 m s^−1^ [80] for common blue-tongued skinks (*Tiliqua scincoides*) is actually almost eight. Our mechanistic model enables us to offer a causal explanation not only for the allometric ‘main road’ of maximum running speed, that is, the legged speed allometry v_*max*_(*M*), but also for observed variances.

To calculate the allometric ‘main road’ v_*max*_(*M*), we simply have to fix all body-, leg-, and muscle-specific parameter values as either constants or parametrised functions of body mass *M* (Table 2). Beyond such calculations of the legged speed allometry, we can also use the model to symbolically calculate the impact of any model parameter on any solution v_*max*_(*M*)—that is, calculate the parameters’ percentage sensitivities of v_*max*_(*M*) with respect to their percentage changes in value, in the vicinity of the solution. Rigorously and systematically calculating locally selective sensitivities enables us to grasp the impact of physical and physiological body design on maximum running speed. Section 4.1 then applies and discusses a common global version of sensitivity analysis—finite parameter value variation—to bring this impact illustratively alive. There, we demonstrate how realistically chosen *finite* changes in parameter values determine maximum speed *aside the main road* of the (mean) legged speed allometry v_*max*_(*M*), at a *given body size* (*M*).

We now briefly explain, and then conduct, a local sensitivity analysis [81] to rigorously calculate the (local, differential) effect of all model parameters on v_*max*_. In prior studies, the dynamics of muscle activation [82] and isometric contractions [37] had already been investigated, using a similar method. Here, we can apply a simplified version because our model enables us to calculate all solutions (Fig. 2) from an algebraic equation (Eq. (G.2)) in closed form (Eqs. (H.4,H.7)), rather than from integrating ordinary differential equations [37, 82]. Given the above-defined model for v_*max*_ = v_*max*_(*M,* Λ) (see Eq. (21) and again Eqs. (H.4,H.7); for the set of parameters Λ defined in Eq. (A.3) see Tables 1,2), we define the *local first-order sensitivity* 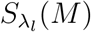 of our calculated model solution v_*max*_(*M*) with respect to any model parameter *λ_l_* ∈ Λ as

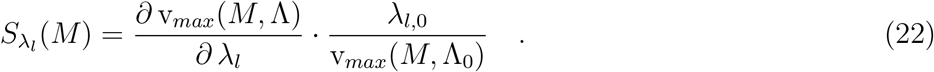

Here, Λ_0_ denotes a particular set of parameter values *λ_l,_*_0_ ∈ Λ_0_ that serves as a *local reference*. Some restrictions are inherent to the significance of local sensitivity analysis [83] and there are several global approaches [84, 85, 86] to overcoming them. In any event, the major benefit of local sensitivity analysis consists in gaining a breezing, quantitative overview of how any parameter effects the model output (solutions): At each particular mass value, 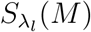 is the percentage change in v_*max*_(*M*) per percentage change in *λ_l_*. The direct comparability of parameter sensitivities 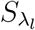 is due to normalising in Eq. (22) all the changes in solution (*∂* v_*max*_(*M,* Λ)) and any single parameter (*∂ λ_l_*) to each a respective reference value (indicated by ‘_0_’).

For the solutions v_*feq*_(*M*) and v_*off*_ (*M*) (Fig. 2) predicted by our initial parameter guess (column three: ‘initial guess’ *λ_l,ini_* in Table 2), the calculated sensitivity functions 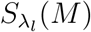 for v_*max*_(*M*) ∈ {v_*feq*_(*M*), v_*off*_ (*M*)} that exceed the value 0.1 are displayed in Fig. 8. An exemplary sensitivity of 0.5 (or −0.5) at given mass *M* indicates that a parameter increase of 30 % would result in an increase (or decrease, respectively) of 15 % in v_*max*_ at this mass. The sensitivity functions of parameters that occur solely in products, such as *ρ_med_* and *C*_∞_, are congruent and, thus, consolidated in the key of Fig. 8. Sensitivities for v_*max*_ are obtained by reading the function values for the solution v_*feq*_ below *M* = 16.5 kg and otherwise reading values from the v_*off*_ solution.

**Figure 8:**
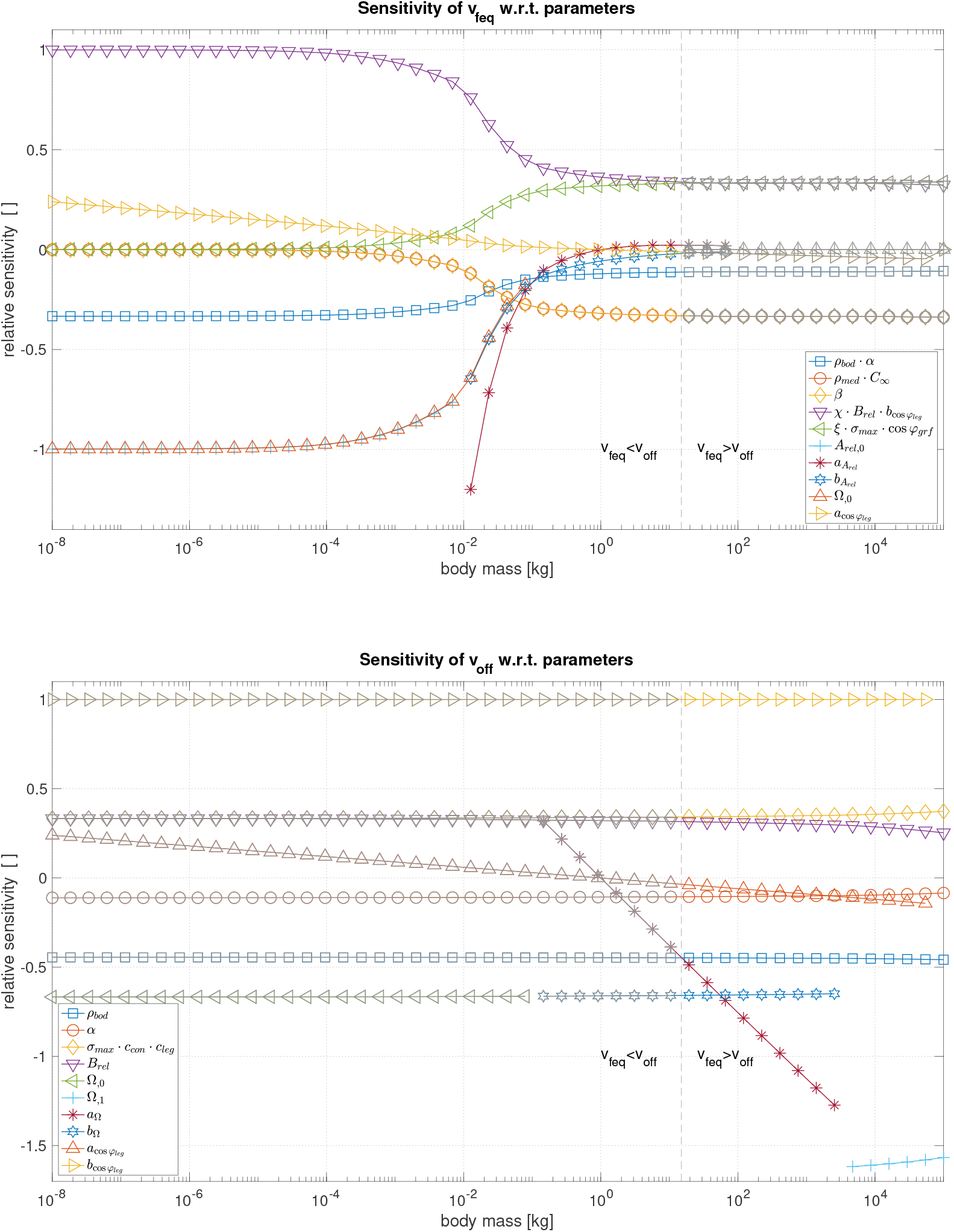
Parameter sensitivities vs. body mass *M*. Plotted are the normalised sensitivities (Eq. (22)) with respect to the model parameters (see Table 2) for the values of maximum running speed v_*max*_ as calculated from (top) the force equilibrium (Eq. (2): v_*feq*_) and (bottom) the constraint of finite available settle time (duration constraint) for accelerating the muscle to high concentric muscle contraction velocities, i.e., premature termination of muscle work (Eq. (18): v_*off*_). v_*max*_ = v_*feq*_ applies to *M* ≲ 16.5 kg, v_*max*_ = v_*off*_ = 13.9 m s^−1^ to *M* ≳ 16.5 kg (see Fig. 2; parameter values: initial guess). Data are plotted for all those parameters with the magnitudes of their respective sensitivities exceeding 0.1 within the considered range of body sizes. A normalised sensitivity value of 1.0 means: v_*max*_ doubles if the respective parameter value doubles.

Overall, most sensitivity functions are approximately constant below 10^−2^ kg and above 10 kg. We recall that, in our initial guess, we have assumed that *A_rel_* and Ω change with size between about 10^−2^ kg and 10^2^ kg, as well as 10^−1^ kg and 4 · 10^3^ kg, respectively, (Fig. 5, Table 2). Exceptions are the sensitivities of 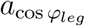 for v_*max*_ ∈ {v_*feq*_, v_*off*_}, as well as *a_ccon_* and *a*_Ω_ for v_*max*_ = v_*off*_. For v_*feq*_, the overall most sensitive parameters are the muscle-to-body-length ratio *χ*, the Hill parameter *B_rel_* (assumed as size-independent in our model), the pre-factor 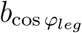 (positive), the basic Hill parameter value *A_rel,_*_0_ or its pre-factor 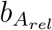, respectively, and eventually the basic EMA value Ω_,0_ (negative). For v_*off*_, the overall most sensitive parameters are, again, 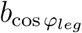 (positive) as well as Ω_,0_ or its pre-factor *b*_Ω_ (negative), respectively.

Further, note that (i) the sensitivities of the parameters *β* (scaling body CSA Æ_*bod*_) and *ρ_med_* (air density) are the same, (ii) the sensitivity of *β* is exactly the negative of the *ξ* (scaling muscle CSA Æ_*m*_), *σ_max_* (maximum isometric muscle stress), and 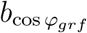 (GRF direction) sensitivities, and (iii) the sensitivities of *A_rel,_*_0_ and *b_Arel_* sum up for the Ω_,0_ sensitivity.

## 4. Discussion

Hirt et al. [1] have published a comprehensive data set of measured maximum speeds of 199 terrestrial species running or hopping with legs. It contains an obvious structure (increase, maximum, decrease)— namely, a curve that is characteristic of evolution-implemented biological design: the *legged speed allometry*. While Hirt and colleagues indicated a relation to limitations of the muscular metabolism, there is currently no causal explanation for this characteristic on the basis of mechanistic laws that govern the design of universal biomechanical constructs for terrestrial locomotion: *mass* (aggregation of body material), *leg*, and *muscle*. Here, we have demonstrated that only two biomechanical ideas are sufficient to causally explain the legged speed allometry—at least up to elephant size (Fig. 4: upward kinks in v_*max*_(*M*))—by physical laws applied to these three constructs.

The first idea is that an *equilibrium* (TOPSS) *between geared muscular driving force and external drag force* determines maximum running speed—at least, for *smaller animals* (below *M* ≈ 40 kg). Herein, the muscular drive itself has an intrinsic, dissipation-caused maximum contraction velocity due to Hill’s force-velocity relation, which in turn limits, geared by leg geometry, the velocity of the COM movement away from the COP (i.e., running speed), even if air drag is negligible (in very small animals).

The second idea is that, in each single stance phase, only *limited time is available for the leg-muscle system to re-feed mechanical work* to the COM for forward running. Herein, the most basic limitation of stance duration comes from a stance length being limited on its part by a finite leg length, which necessarily entails a limited range of leg motion during stance. Given the leg length, any increase in speed shortens the time available for a muscle to achieve, from being isometric (at ISOMS), the demanded high contraction velocity (at TOPSS). This basic limitation of stance duration is part of the duration constraint Eq. (18), which manifests, in our running model, mechanisms other than the force equilibrium Eq. (2) that can induce a leg’s LO to occur *before* achieving TOPSS (therefore, ‘premature’). Regularly, after the running system having achieved the force equilibrium Eq. (2) (TOPSS), the corresponding leg would lose ground contact by decreasing leg force, in combination with the drag overbalancing, the vertical GRF component decreasing, and the sticking tip of the leg starting to slip. As a consequence of the steep dependence of the muscle settle time *τ_m_*(*M*) on body size (Fig. 3), however, increases in size make things worse: The larger an animal, the more likely it is that an LO will be enforced within *τ_m_*(*M*) by another mechanism than as a mechanical consequence of regularly achieving TOPSS. Above a critical body size that corresponds to a mass of ~40 kg (*larger animals*), a muscle’s run-up from ISOMS to settling at TOPSS becomes so slow that the force equilibrium (TOPSS) can not be achieved anymore; then, the muscle’s settle time is predicted to instead determine maximum running speed. The stance duration may be further narrowed due to mechanisms that superpose the leg’s striding function, such as repulsion dynamics against gravity or swing leg inertia.

According to our running model, maximum running speed is, thus, determined by muscle mechanics, particularly during late stance.

### 4.1. Cats and spiders: aside the main road

To explain the allometric ‘main road’ of maximum running speed v_*max*_, i.e., the overall legged speed allometry v_*max*_(*M*), we have represented all environment-, body-, leg-, and muscle-specific model properties by *M*-independent parameters or parametrised functions of body mass *M*. All parameter values used in this study are compiled in Table 2. Here, we would like to illustrate how realistically chosen finite *changes* in parameter values determine maximum speed *aside the main road* of the (mean) legged speed allometry. That is, we examine *variations* in the values of selected parameters (body designs) at a *given body size* (*M*).

Some parameters in our model imply a substantial level of abstraction. The leg geometry in running is characterised by axial joint alignment and muscular gearing (EMA Ω). The geometry of the leg-body interaction—kinematics and kinetics—is characterised by the directions of the leg axis (*φ_leg_*) and the GRF (*φ_grf_*). The maximum contraction speed of a muscle is characterised by the combined visco-elasticity of its fibres and compliant material in series to them (*A_rel_*): e.g., tendon, aponeurosis, or skeletal parts.

We illustrate the effect of design variations on v_*max*_ by comparing a (modelled) human with a body mass of *M* = 100 kg with two (modelled) predators that have the same body mass and that, therefore, have the potential to feed on humans: an invented one (giant spider: similar to the creature ‘Shelob’ [87]) and one that has always been a real-world competitor for meat (big cat: similar to a leopard or cougar). Such a model representation is termed ‘replica’ in the following. Solely, the values of the four highlighted parameters Ω, *φ_leg_*, *φ_grf_*, and *A_rel_* are varied. With this, we both demonstrate their quantitative influence on performance and make graspable the core physical meaning contained in their abstraction. These four parameter values are chosen to be specifically representative of those three designs (Table 3). All other parameters, such as, for example, the optimal length of the bulk of active muscle fibres (*L_opt_*), its CSA (Æ_*m*_) and, thus, both the maximum isometric muscle force (*F_max_*) and muscle mass (*M_m_*), the trunk and leg lengths (*L_bod_* and *L_leg_* = *L_bod_*, i.e., *c_leg_* = 1), and the body’s CSA (Æ_*bod*_) are assumed to be the same in the three designs.

**Table 3:** Replica-specific model parameter values and predicted maximum running speeds. Body mass *M* = 100 kg and all other model parameters are completely identical in the three species-like replicae (third column in Table 2: ‘initial guess’), including, e.g., the muscle fibre length and its maximum isometric force— thus, also its mass. The accordingly predicted maximum speed on the ‘main road’, i.e., for a 100-kg animal replica taken from the initial-guess solution (Fig. 4: ‘model without opt’) is v_*max*_ = v_*off*_ = 12.2 m s^−1^ (v_*feq*_ = 17.0 m s^−1^). Data for the humanoid replica are taken from the biomechanical literature on humans, as given below. The giant spider replica is assumed to have no elastic structure in series to the muscle fibres, nor a tendon nor any other significant compliance such as, e.g., flexible legs. *φ_leg_* is steeper in humans [92] than in the average animal at 100 kg (*φ_leg_* = 45°) and in spiders (*φ_leg_* = 35°). In the big cat replica, it is assumed to be as flat as it is in spiders with *M* 3 g (see [67] for a rough estimate during acceleration). The cat-like value was estimated from sketches of a sequence of running cheetah outlines [93, fig. 1] as well as the leg configuration data of a cheetah at LO [94, fig. 7,top] and associated length data [95, fig. 2a]. Also, due to the fact that cats have claws, which likely enhances the coefficient of friction to *ς* ≈ 2 [77] at LO, we assumed that cats may be able to realise at least *φ_grf_* = 45° (i.e., *ς* = 1), in contrast to humans and spiders. In the spider and cat replicae, maximum running speed v_*max*_ is predicted to be determined by the force equilibrium (Eq. (2): v_*feq*_), whereas v_*max*_ is limited in the humanoid replica by the stance duration constraint (Eq. (18): v_*off*_).

| replica | $v_{max}$ [m s<sup>-1</sup>] | $\Omega$ [] | $A_{rel}$ [] | $\varphi_{grf}$ [°] | $\varphi_{leg}$ [°] |
| --- | --- | --- | --- | --- | --- |
| humanoid | 10.7 (= $v_{off}$; $v_{feq} = 15.9$) | 0.70 | 0.001 | 63.4 | 55 |
| big cat | 20.6 (= $v_{feq}$) | 0.35 | 0.001 | 45 | 35 |
| giant spider | 15.7 (= $v_{feq}$) | 0.15 | 0.05 | 63.4 | 35 |

At a body mass of 100 kg, all model parameter values of the average animal replica ‘built’ by our initial parameter guess from literature data (Table 2, third column) approximate, in fact, a human sprinter, except for the leg angle, which is assumed to be *φ_leg_* = 45° on average, but *φ_leg_* = 55°, i.e., more upright, in humans (*Homo sapiens*). Thus, the calculated maximum running speed v_*max*_ = 10.7 m s^−1^ (labelled as ‘humanoid’ in Table 3) is a first promising indicator of the validity of our model: For exceptional male athletes, values in the range 11.9 m s^−1^ [78] to 12.5 m s^−1^ [88, 89] have been measured. Remarkably, our running model predicts that the maximum speed of running with a human-like body-leg design is generally limited by the stance duration constraint (Eq. (18)) for which a crucial parameter is leg length (see Eqs. (J.6,J.7) and Sect. (3.2)): A too-short stance duration induces premature termination (LO) of the acceleration of the muscle’s own mass towards its contraction speed that corresponds to TOPSS (fulfilling the force equilibrium Eq. (2))—that is, the generation of muscle work is prematurely terminated. Thus, the humanoid maximum running speed is always located on the ‘descending branch’ (see Fig. 2: solution v_*off*_ (*M*) of Eq. (18) bearing Eq. (8) in mind) of the legged speed allometry v_*max*_(*M*). If muscle inertia were neglected, our model would predict (see Table 3) that humanoid 100 kg-sprinters would be able to run at v_*feq*_ = 15.9 m s^−1^ (solution of Eq. (2) applying to smaller animals).

We also find, not surprisingly, that the cat replica is much faster than the humanoid replica (Table 3). Three of our examined parameters add to this racing advantage: The more crouched leg [10] (cat-like Ω assumed as half the human-like value from [90]) enables a cat to approximately double the axial leg lengthening velocity (Eq. (7)) with an identical muscular design. Moreover, the flatter angles *φ_leg_* and *φ_grf_* foster the transmission of the axial lengthening velocity (Eq. (6)) and leg force (Eq. (17)), respectively, to forward direction. An anatomical property that enables a cat to implement a flat leg angle *φ_leg_* is discussed in Sect. 4.3: A cat is equipped with a flexible spine, which helps enlarge its stance length (Eq. (J.5)). Flat GRF angles *φ_grf_* are enabled by claws [77] (see further below). Combined, these effects endow the cat replica with almost twice the humanoid maximum running speed, despite the increased drag and halved axial leg force (due to Ω) of the cat-like design.

There are two main reasons, why, in the cat replica, the stance duration constraint (Eq. (18)) does not kick in, such that this replica can fully exploit its racing advantage over the humanoid. First, the significantly flatter leg angle allows a longer acceleration distance from ISOMS to TOPSS (stance length: Eq. (J.5)) and, thus, a longer stance duration (Eq. (J.7)). Second, *at the same running speed* v, the muscle—identical in both designs—performs at a demanded contraction velocity v_*m*_ that is *lower* for reduced Ω (Eq. (8)). The muscle can, thus, generate a *higher* running speed, though at a higher muscle force demanded, which is closer to the isometric force demanded at ISOMS. Thus, less time *τ_m_* is required to accelerate the muscle’s own mass to achieve the muscle’s force value corresponding to the force equilibrium (Eq. (2)): *τ_m_*(v_*m*_) (Eq. (I.19)) decreases with steeper *f_m_*(v_*m*_) (Eq. (I.11)), i.e., with increasing 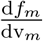 (Eq. (I.13)). Admittedly, the contraction speed advantage by lower Ω comes at the cost of an associated decrease in axial leg force (Eq. (14))—thus, driving GRF (Eq. (17)). Yet, a mechanism that effectively over-compensates for such a loss in driving force due to leg transmission is immanent in Hill’s [39] hyperbolic force-velocity characteristic of the muscle (Eq. (I.1)): Any reduction in the demand for muscle contraction velocity due to a (linear) decrease in Ω (Eq. (8)) is linked to a more-than-linear increase in muscle force (Eq. (I.2)). In addition, the flatter GRF angle *φ_grf_* in cats (Table 3) provides some other compensation for an Ω-induced reduction in the axial leg force by making better use of the leg force for forward-driving GRF (*F_grf_*) generation (Eq. (16)). According to our model, all these properties, combined, equip cat-like creatures with a substantial racing advantage over humanoid designs.

The invented spider replica is also clearly faster than the humanoid one. From spider-like Ω being much lower than even that in cats, one might have expected that the giant spider could hunt down the big cat replica. Luckily for cats, spiders are not supposed to have elastic tendons, which is reflected by the spider replica’s fifty-times-higher *A_rel_* value (Table 3). Functionally, the maximum contraction velocity of a spider-like muscle is accordingly limited to solely muscle fibre output (non-steady-state [36, 37, 38]), as it is stripped of the velocity-amplifying capacity of a tendon. The probably steeper GRF angle (*φ_grf_*) also counteracts the EMA (Ω) benefit. There are indications [67] that accelerating spiders’ leg angles are even lower than our conservative initial guess *φ_leg_* = 35°. Equipped with such flatter legs in the TOPSS condition, a geometry-tuned spider replica would be able to partially close the remaining racing gap on a big-cat-like creature. Further conceiving that a giant spider replica incorporated additional serial elasticity (tendons or other elastic leg structures), such creatures would then be able to even hunt down cats of their own size.

The consequence of an approximately five-times-lower Ω in a spider-like leg design as compared with a human-like leg design is that, to slightly over-compensate bodyweight with the same maximum isometric force *F_max_* of the muscle available, our spider replica requires four legs on the ground (or three at minimum if significant eccentric muscle force enhancement [91] is present) at some point within an entire gait cycle. This is because the muscle parameters are chosen human-like. Thus, the humanoid leg generates one and a half times the bodyweight axially if the muscle acts isometrically (at *F_max_*: see Eqs. (E.2,E.3) and the chosen *ξ* value). With a spider-like leg geometry (Ω), such muscle parameter values result in significantly reduced maximum isometric leg forces: Axially, one leg of the spider replica can generate only thirty percent of the bodyweight.

Eventually, we examine what our model predicts for the commonly accepted ‘king of speed’—the cheetah (*Acinonyx jubatus*). The measured benchmark is 29 m s^−1^ [76], as an average over a 200 m distance, with, thus, some potential for even higher values [77]. Compared with other, less-speed-specialised big cats, a cheetah is more lightweight: we assume a 50 kg bodyweight, which may represent a very well-built specimen [77]. Assuming again, for a start, that all model parameter values represent the ‘main-road’ v_*max*_(*M*) (initial guess: Table 2, third column), and then superposing the four crucial ‘aside-the-main-road’ parameter values of our cat replica (Table 3), we try to further speed-optimise this cat-like design by adopting another five species-specific parameter values to make the cat replica more cheetah-like: The body is made more slender in breadth as compared with length (and certainly in height, particularly at late stance), which reduces the body’s CSA that faces airflow, by assuming *β* = 0.15 (compare all optimal parameter values in Table 2). The muscle fibres are made faster by assuming *B_rel_* = 10 s^−1^ and a little stronger by assuming a maximum isometric stress value of *σ_max_* = 3.0 · 10^5^ N m^−2^. Moreover, the use of ground friction in sandy or grassy savannah is pushed to the physical limits by means of claws [77], which enables a coefficient of friction at late stance of up to *ς* ≈ 2 and, thus, a GRF angle as flat as *φ_grf_* = 26.5°. With these parameter values, a maximum running speed of v_*max*_ = v_*feq*_ = 36.1 m s^−1^ is predicted for a cheetah *if* the leg length is additionally assumed to be double the trunk length (*c_leg_* = 2, the optimal values in Table 2), e.g., by combining a flexible spine plus space- and time-shifted deployment of two cooperating legs in a gallop (see Sect. 4.3). It is likewise predicted that the cheetah’s running speed would be limited to v_*max*_ = v_*off*_ = 29.4 m s^−1^ by muscle inertia and premature LO, i.e., the solution of the stance duration constraint (Eq. (18)), if the leg were as short as assumed in our initial guess (*c_leg_* = 1). By sticking to *c_leg_* = 2, yet again more conservative guesses for the friction coefficient (*ς* ≈ 1, i.e., *φ_grf_* = 45°) and the maximum isometric stress (*σ_max_* = 2.5 · 10^5^ N m^−2^), we would still predict unconstrained v_*max*_ = v_*feq*_ = 31.5 m s^−1^, and a short leg (*c_leg_* = 1) would then put the limit to v_*max*_ = v_*off*_ = 27.6 m s^−1^.

### 4.2. Reliability of maximum running speed data

The data set compiled by [1], with a few points of some species added and used here, provides measurements of the maximum running speeds of a wide range of legged terrestrial species. However, under laboratory conditions, it is notoriously difficult to spur specimens to perform with maximum effort. Under both laboratory and field conditions it is likewise difficult to assess the degree of effort an animal yields. Thus, most certainly, some species’ capabilities are significantly underrated. Moreover, many species simply do not have evolved body plans facilitating particularly high running speeds. In fact, the locomotor apparatuses of most of the compiled species trade off between a couple of demands [55], of which high speed locomotion is only one. Accordingly, one might argue that only the upper rim of the scattered data cloud in Fig. 4 reflects the capabilities of locomotor apparatuses that are optimised to some degree for maximum sprint speeds (see below). However, even the upper rim might contain some erroneous data points, as muscle power is significantly affected by the temperatures at which muscles operate, particularly in small poikilotherms such as arthropods. Therefore, ambient temperature can significantly affect running performance. In this light, not only the extraordinary running speeds measured for teneriffiid mites [96] but also those for Australian tiger beetles [97], which both seem to significantly exceed the calculated values, must be scrutinised. Spagna et al. [98] and Full & Tu [49] also did not provide temperature conditions for their experiments with apparently extremely fast spiders and cockroaches, respectively.

Interestingly, good data on fast-running bipedal species seem to be largely missing, with the only exception being ourselves, human beings. However, human running performance is in no way optimised for achieving particularly high maximum speeds. On the contrary, humans are rather on the slower side (see Tables 2,3) of the mean legged speed allometry, similar to penguins [99] or, let us say ducks [100], in order to avoid too harsh a comparison.

However, our approach was not at all aimed at speed optimisation for—or by—specific body plans. With our assumed and optimised parameter values (Tab. 2), the model reveals the principally attainable running speeds if one abides by the built-in scaling mechanisms (Fig. 4 and Tab. 2). Nevertheless and beyond, the model can be used to causally explain species-specific deviations from the (mean) legged speed allometry (see Sect.4.1).

### 4.3. Multiple legs on the ground and adaptions towards maximised work output

In this section, we will address the interplay between the different scaling parameters and the resulting speed-limiting effects for distinctly different body plans. For that, we first use the examples of birds. Then we compare bi-, quadru-, and polypedal species, focusing in particular on considerations of the relation between body plans, chosen gaits, and work maximisation.

#### Bipedal vs. quadrupedal designs

As Fig. 4 shows, the existing data for a range of bird species is consistently below the maximum speeds achieved by their quadrupedal fellow species (of the same size). From a mechanical perspective, the birds’ anatomically generally longer legs [101] (Fig. 9) should facilitate longer stance durations, given the same leg and GRF angles at LO or TOPSS. Moreover, bipedalism has the advantage of low to lacking interaction forces [102, 103] (GRF interferences) between the legs as well as resulting in simplified control [104]. This particularly holds at high running speeds, as the legs’ temporal overlap diminishes there. While it seems as though the birds’ longer legs cannot fully compensate for the lack of a flexible spine (see below), it might well be that the current relative scarcity of data on maximum running performance in birds is the reason for the apparent speed leeway. Moreover, particularly under field conditions, many bird species will certainly simply fly off when higher speeds are required and usually do not need to fully exhaust their terrestrial capabilities. The capacity to fly may also alter their behaviour under laboratory conditions.

**Figure 9:**
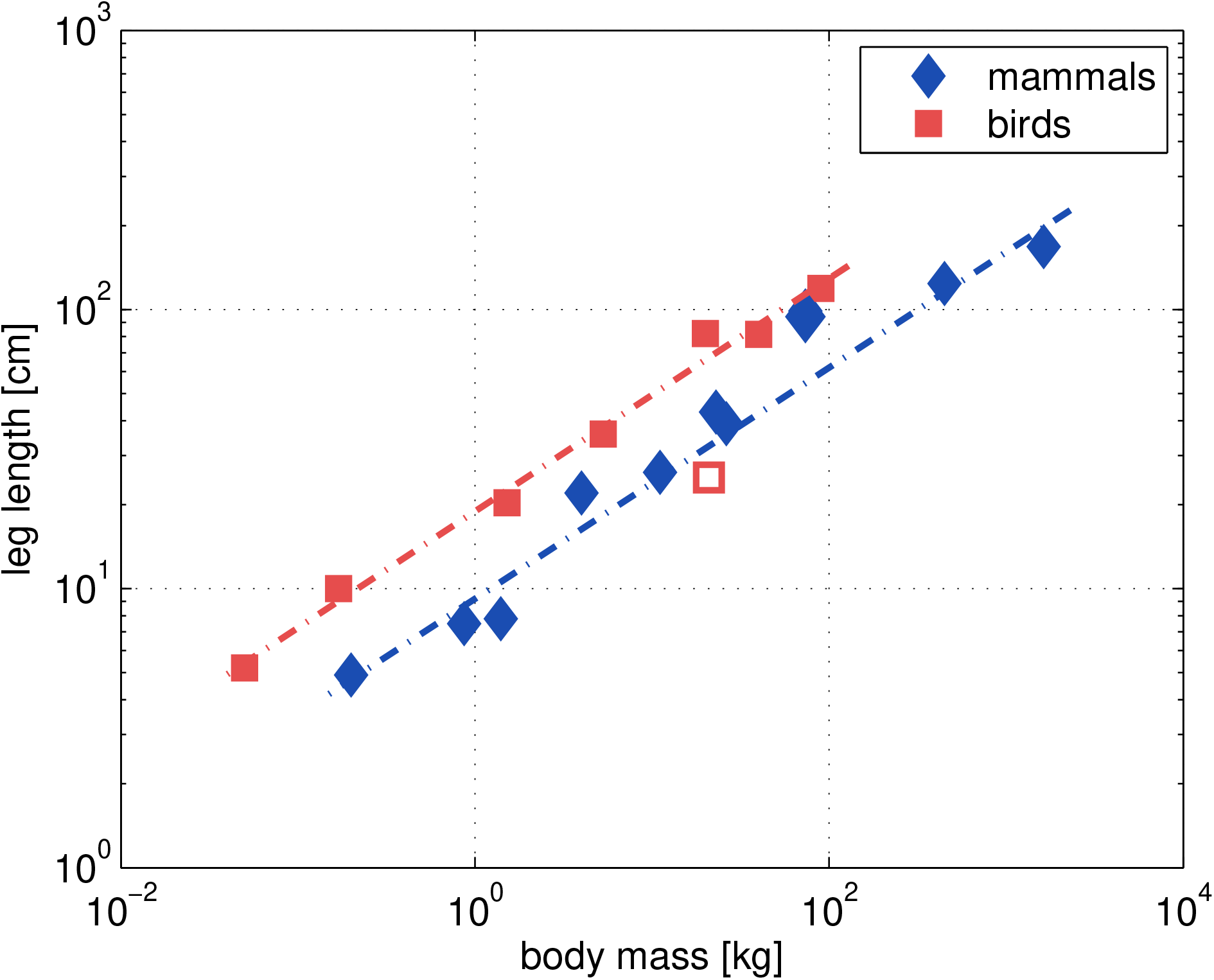
Experimental data taken from Pontzer [101]: *In situ* leg lengths of birds and mammals vs. body mass *M*. The lengths had been measured from the distal end to the hip joint in standing posture. The unfilled square represents the emperor penguin with comparably short legs and the typical waddling gait. Humans and reindeer had particularly long legs and similar body masses of about 73.5 kg, such that their markers superpose each other and are close to the bird regression line. The data points for the penguin and for humans with their deviating body plans were not considered for the calculations of the least-square fits (dash-dotted lines).

#### Scaling of the leg’s skeletal elements

The long bones of large quadrupedal mammals scale differently with size from those of small quadrupeds [105]. Interestingly, this seems valid for lineages with as different ecological adaptations and phylogenetic descendants as felids, bovids, and ceratomorphs. Therefore, similarly-sized species can reasonably be compared with each other.

In small quadrupedal species, the length *L_bone_* and diameter *D_bone_* of long bones scale almost linearly with each other: 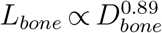 [75, 105]. Despite this observation of an almost geometrical allometry of bone dimensions, on increasing body mass, bone stresses can still be maintained at constant levels by means of aligning the joints with the (leg’s) GRF vector [74] above a critical value of approximately 0.1 kg: here, EMA increases with body mass [58] (Fig. 5, Table 2). While quantitative EMA data of species larger than horses are non-existent so far, it seems clear that alignment is fully exploited in the largest mammals, like elephants, rhinos, and large ungulates, such that the robustness of long bones must increase in order to keep peak stresses within safe levels [106]: The long bones of very large species then scale nearer [75, 105] to what a design by elastic similarity [7, 107] 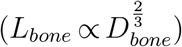 or even static stress similarity [7, 75] 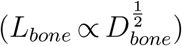 predicts. However, full EMA adjustment does not mean that limbs are completely straightened (aligned), particularly not at MS. In galloping species with otherwise distinctly aligned limbs, such as many large ungulates, the legs are still somewhat more crouched at MS, which attends a decrease in COM height and significant vertical deflection amplitudes when approaching their highest speeds (e.g., [108, 109]).

#### Gaits: the construct of a single leg and its concept passed on to a set of legs

In those many running species equipped with more than two legs, the exertion of a leg set’s total leg force on the ground depends not only on single leg properties but also on the leg coordination pattern an animal chooses at a specific speed. While, at maximum speeds, most bipedal animals alternate their legs (only kangaroos and some passerine [110] and rodent families prefer to hop), animals with more than one pair of legs can adopt a much higher number of leg coordination patterns by shifting the phase relations of the ipsilateral legs, i.e., by changing the temporal coordination of ipsilateral legs. Nevertheless, some of these coordination patterns result in largely equivalent body dynamics—and, accordingly, gaits. For example, in quadrupeds, pace and trot result in equivalent COM and energy fluctuations, though the activity of ipsilateral legs is in phase in the former and out of phase in the latter. Furthermore, both coordination patterns are dynamically equivalent to the tripodal coordination scheme of insects and bipedal running, with the kinetic and potential energy of the COM oscillating out of phase with each other, at exactly double the stride frequency. All four running schemes are found predominantly in species whose legs are equipped with anatomical features that make possbile the transient elastic energy storage and recovery, which facilitates energy-efficient locomotion [45, p. 255], [47, p. 479], [111], [112], [113, p. 197], [28, p. 299].

However, the more (walking) legs an animal’s body plan contains, the more difficult it is to attain sufficient synchronisation among the legs of the potential sets of legs. Yet, synchronisation is a requirement for an efficient deployment of elastic leg structures in the repulsion dynamics of the COM against gravity: A pooled absorption of elastic energy is indispensable for both decelerating the COM in its downward movement and, likewise, re-accelerating it upwards, attending smooth and pronounced GRF peaks [104] (spring-mass dynamics [26, 27, 114, 115]). The deployment of elastic structures for transient energy storage and recovery at high speeds, i.e., with relatively low duty factors, therefore, seems to be confined to body plans with three or fewer pairs of (walking) legs. In addition, the implied spring-mass dynamics in the sagittal plane [26, 27, 114] require vertical amplitudes of the COM during the stance phase increasing with running speed, along with the compression rate of the compliant leg. Accordingly, spring-mass dynamics and synchronised sets of legs can be deployed efficiently only (i) with sufficient clearance of the body above the substrate and (ii) if the leg’s compression rate neither significantly hampers repulsion dynamics nor makes the energetic balance inefficient through excessive dissipative losses. In many arthropods and in confined spaces this is apparently fulfilled at best up to intermediate speeds. At high and top running speeds, blaberid cockroaches, mites, and spiders [54, 55, 56], and probably many other fast polypedal organisms, change their leg coordination scheme and temporally dissociate the previously synchronised sets of legs. This makes vertical COM oscillation amplitudes decrease and attain absolute minima at ipsilateral phase shifts specific to a number of propulsive legs [104]. At these specific phase shifts, which were also found in behaving animals [55, 56, 116], body dynamics are no longer spring-mass-like; a single leg’s stance duration is then determined by its functional length as well as running speed rather than by anti-gravity repulsion dynamics. Such phase shifts also lead to the elimination of airborne phases as typical for running and trotting gaits.

Through a combination of our mechanical model, muscle energetics known from physiology, and empirical data on animal acceleration, we have found good arguments (see Sect. 4.4) supporting the key idea that TOPSS (Eq. (2)) is determined by the properties of ‘the last one (muscle) standing’ (Eq. (G.1)) at late stance of a single functional leg (Fig. 1). This key idea can immediately be passed on from a single-leg model consideration to real-world situations with multiple legs on the ground. In the latter, the time courses of force and work may be more subtle during a stride cycle, meaning that the force and energy fluctuations are low and occur at higher frequencies. Yet, any of the possibly multiple TOPSSs during one stride can be detected by fine-structured analysis of COM kinematics. If there is a maximum in the forward component of the COM velocity, this is a TOPSS, of which identifying ‘the last one standing’ would be an experimental challenge. Mechanically, one leg (or muscle) in a set supports the other by prolonging the available periods of doing work, be it for anti-gravity action or propulsion in the running direction. Multiple leg action is an alternative to designing longer legs: Both strategies increase the available time periods for doing work (at the same speed). The multiple-leg strategy decouples overall work-generation from the repulsion dynamics of a single leg. For this strategy to work, the running organism must apparently forego the deployment of efficiency-enhancing elasticity for reversing the vertical momentum.

#### Gait changes: the emerging advantages of the gallop

Many quadrupeds, in turn, are able to increase their stance (ground contact) lengths when running faster and faster, with accordingly approaching a lower limit in their stance durations [77, 117, 118]. Having arrived at their high-speed regime, these animals have changed to one of the various galloping gaits (canter, half-bound, bound, transverse gallop, and rotary gallop) [119, 120]. It is further known that horses as well as cheetahs [121], dogs [94], and small mammals [122, 123, 124, 125] extend their anatomical legs and flexible pelvis-lumbar-spine-complexes in combination when galloping. Therefore, by integrating their flexible spines in fast running movements, they effectively increase their leg length proximal to the hip joint (see Fig. 1: functional leg concept). The aforementioned galloping gaits are kinematically defined, and, therefore, often labelled ‘asymmetrical’ [126]. Asymmetrical gaits are characterised by diminishing phase shifts within each of the pairs of legs. Accordingly, the sequenced and overlapping ground contacts of the four legs substitute for a ground contact of synchronised sets of (diagonally opposite) legs in symmetrical gaits like trot and pace. Consequently, downwards-upwards redirection of the COM occurs only once per stride and not two times per stride as is typical for symmetrical gaits [127]. Because the redirection is now distributed in time over the sequenced ground contacts of the legs, galloping gaits offer relatively longer overall stance durations and a reduced cost of transport [128, 129], because the legs can be stiffer (more aligned: higher EMA). Therewith, compliant excursions are reduced: in galloping gaits, the forelegs act in a manner similar to that of the spokes of a rimless wheel [130, 131]. They mainly redirect the vertical movement of the COM, while the trailing hindleg, in particular, provides significant propulsion by means of leg lengthening [132, fig. 3], with muscles generating positive work. Therefore, at speeds close to the absolute maximum, the forelegs’ braking and re-accelerating forces neutralise each other, whereas particularly the trailing hind leg is refeeding the majority of the energy (see [77, fig. 5]) lost due to air drag and body-internal friction processes.

In species of distinctly different sizes [117, 133], the stance durations in galloping seem longer in the hindlegs than in the forelegs. Combining this with (i) the basic geometric fact that, for the running speed v and the angular range *φ_leg_* given, a leg’s stance duration (Eq. (J.3)) increases linearly with its length (Eq. (J.4)) and (ii) the observation that the stance durations approach lower limits at the highest speeds compellingly suggest that spinal flexibility is in operation in the gallop of fast runners. Additionally, the anatomical lengths (completely extended or aligned, respectively) of the hindlegs usually slightly exceed those of the forelegs in fast quadrupedal runners [134, 135], which probably contributes to the hindlegs’ longer stance durations and lower leg angles with the substrate at LO, and further prolongs the time frames for accelerating the leg muscles to maximised contraction speeds. To sum up, the resulting functional leg-lengthening mechanism by spinal flexibility facilitates high maximum running speeds through work enhancement during a leg’s stance phase. This mechanism gives the galloping gaits an advantage over symmetrical gaits when it comes to maximum speed performance.

Moreover, like locomotor apparatuses as a whole, spines must meet variable, sometimes contradictory, demands. Spines provide sufficient flexibility but must just as much maintain the body posture against gravity. Particularly in very large animals, the static gravity-support aspect gains importance (see bone allometry above, [7, 75, 105]), which results in comparably stiff lumbar spines [136]. However, reduced spine flexibility reduces the stance length of the legs, particularly when gallop dynamics are deployed. For a given running speed, therefore, the time available for muscle contraction and GRF application is accordingly reduced. The described anatomical restriction is, thus, expected to contribute to the observable decrease in the speed data for animals with large body masses. This tendency has already been incorporated into our current model (see monotonous decrease in cos *φ_leg_* with body size in Fig. 6).

#### Polypedal body plans

In arthropods, a clear gallop gait has so far been observed only in semi-terrestrial ghost crabs that swiftly roam the intertidal zone of their tropical and subtropical shore habitats. Owing to the animals’ short body axis and long (walking) legs, the sideways orientation during locomotion increases the legs’ range of motion and prevents perturbations between them. At maximum speed, however, these crabs use only the longest of their legs, which reduces the number of active legs to two [137, 138]. In contrast to the bodies and muscles of vertebrates, a crab’s body is fairly rigid and its trunk muscles cannot contribute to the generation of propulsive forces. Therefore, the crab’s gallop is essentially dynamically similar to human and passerine skipping [139, 140], i.e., a gait not standing out due to particularly high speeds. Accordingly, the ghost crabs’ maximum running speeds are not exceptional compared to speeds of vertebrate species with similar body masses (Fig. 4); medium-sized animals of the species *Ocypode ceratophthalma*, with a body mass of about 15 g, achieved approximately 2 m s^−1^ [137]. Slightly heavier animals of the species *Ocypode quadrata* (about 50 g) achieved speeds of up to 1.6 m s^−1^ [141]. These are the maximum observed values for all body sizes of the respective species.

As has been recently shown, other polypedal animals like insects and arachnids also deviate from the (trot-like, symmetrical) alternating tripodal or quadrupedal leg coordination scheme at high and maximum running speeds [55, 56]. The cockroach *Nauphoeta cinerea* as well as a trombidid mite dissociated their sets of legs resulting in temporally distributed ground contacts, with the legs’ phase relations becoming more complicated. In principle, this is similar to the sequence of leg contacts in the galloping gaits of quadrupedal vertebrates, as compared to their trotting. Because, essentially, two sets of legs still alternate with each other in these arthropod species, their COMs continue to oscillate with two times the stride frequency, just like in more typical symmetrical gaits. However, if vertical accelerations of the COM in the purely tripodal gait are compared with those in the high-speed metachronal gait (i.e., with temporally dissociated sets of legs) of the cockroach and the mite, the accelerations of the former gait are significantly higher [56]. Simultaneously, the accelerations in the fore-aft direction and absolute running speeds were higher in the latter gait. Similar trends seem to occur in horses [142]. Both findings indicate the superiority of running with dissociated sets of legs when it comes to attaining maximum speed.

Taking into account extremely high relative running speeds, such as those found in tiger beetles [97] and mites [96], and the incompleteness of the available data, dissociated sets of legs might also be used by the fastest of the small arthropods. With duty factors lower than those found in the medium-fast speckled cockroach and a slightly lower deviation from the alternating pattern, the sequenced legs of the alternating sets of legs would be intermitted by aerial phases [104], bringing body dynamics even closer to galloping dynamics and providing further increased stride lengths due to the introduction of a ballistic phase. Such hybrid body dynamics would take advantage of both gallop dynamics with temporally distributed leg force application to the ground and prolonged swing phases, as well as the maintenance of the general control pattern of polypedal running. This general pattern, again, may mitigate fatigue in the single legs, as two sets of legs still alternate with each other: Those muscles in the swinging legs that have mainly supported the weight during their previous ground contact are given prolonged periods to recover their energy reservoirs. Indeed, the required decrease in leg synchronisation within the alternating sets has even been demonstrated for very fast running arthropods such as desert ants and fast-moving spiders with duty factors as low as 0.25 and 0.35, respectively [98, 143]. Unfortunately, the crucial analyses regarding leg coordination and COM oscillations were not included in these studies.

### 4.4. Lizards and lions: a critical analysis of the “general scaling law” [1]

#### Point of departure

Introducing a “general scaling law”, Hirt et al. [1] claimed that their hump-shaped v_*max*_(*M*) curve for fitting the legged speed allometry was obtained by conflating three mass-dependent quantities: (1) the theoretically possible maximum speed v_*max*(*theor*)_ = *a · M^b^*, (2) the acceleration ability *k* = *c · M^d^*^−1^, and (3) the critical time *τ* = *f · M^g^* available for acceleration until the fatigue of white muscle fibres. In their course of parameter identification by fitting, the latter two effects were indistinguishably lumped together as *k · τ* = *c · f · M^d^*^−1+*g*^:= *h · M^i^*. Here, we show that it is absolutely possible to attribute a mechanism or a combination of such to each of these three quantities. In the following, we use literature data and unroll a physiological mechanism in order to scrutinise the mechanism of white muscle fibre fatigue as being an explanation for introducing *τ*. With this, the justification in [1] for *explaining* a potential overall maximum in the legged speed allometry by a data fit that assumes the product of two exponential functions collapses.

**Ad (1)**: Regarding Hirt et al.’s literature sources [25, 144], we note three issues: first, in [25], only work-optimised speeds are presented rather than maximised ones. Second, in the same source, the presented power law was fitted despite a visible hump (saturation of v_*max*_ at higher masses). This hump is not visible in [144] because no animals above 100 kg have been considered [9]. Third, the key figure [144, fig. 6.4] contains an erroneous ordinate scaling. Our model shows that the assumption of a classical (non-humped) allometric relation between *M* and v can be obtained if neither the serial elastic (tendon) nor the EMA properties are changed across scales (see Fig. 2 and App. L, and thick black lines). In the small-animal limit case, our power law slope *b^model^* is approximately 0.32 (see the beginning of Sect.3.3), which is higher than *b^lit,^*^1^ = 0.24 (estimated by [5, fig. 4] including hump data) but lower than *b^lit,^*^2^ = 0.38 ([144] without hump data). Air drag then induces reductions of the slope *b^model^* at larger sizes (thick black line in Fig. 2). The optimised value *b^fit^* = 0.26 obtained in [1] is, thus, reasonable.

**Ad (2)**: For the acceleration ability *k*, [1] reported a literature value of the power law slope *d^lit^* between 0.75 and 0.94, assuming that 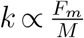 (see Eq.(29) below) directly reflects the muscular drive in the leg. Based on geometric considerations, the muscle force *F_m_* should scale with its CSA Æ_*m*_, i.e., with 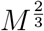 (see our Eq. (E.3)). Relating *F_m_* via EMA-gearing (Ω in Eq. (17)) to its transform *F_grf_*, our model then yields *d^model^* ≈ 0.67 for smaller animals (before the onset of change in EMA) and *d^model^* ≈ 0.67 + 0.25 = 0.92 for larger animals (see Table 2), which is in perfect congruence with the literature. To distinguish this quantity from the fibre fatigue effect proposed by Hirt et al., we examined their cited work and directly calculated the acceleration ability *k* of lizards (*Stellio stellio*) [145] and lions (*Panthera leo*) [146] by using the time course v = v_*max*_ · (1 − *e^k^*^·*t*^) (see Eq.(28) below) of speed from the latter source. The lizard with a mass of 0.035 kg is able to achieve 95% of its maximum speed (3 m s^−1^) within 0.25 s [145, fig. 1], hence *k^lizard^* ≈ 12.0 s^−1^. Further, assuming that a male lion weighs 175 kg and has the ability to achieve 95% of its maximum speed (13.9 m s^−1^) within 4.4 s [146, fig. 5], we obtain *k^lion^* ≈ 0.68 s^−1^. Solving the respective two-point form yields *c^empir^* ≈ 3.9 kg^0.34^ s^−1^ and *d^empir^* ≈ 1 0.34, thus, *d^empir^* ≈ 0.66. This value confirms our above model-based consideration of muscle force scaling, as lions have crouched legs with an EMA clearly lower than that of humans (see Sect. 4.1), probably similar to much smaller animals [10] like lizards.

**Ad (3)**: The crucial part of the “general scaling law” is certainly the emergence of the speed hump, which mathematically necessitates the substitution of the size-dependent parameter *τ* for the independent variable ‘time’ describing the acceleration process [1, eq. 2] (see also Eq. (28) below). Moreover, as insinuated above, the assumption of *τ* being linked to muscle fibre fatigue is doubtful. In addition to a missing mechanistic foundation of this assumption, the confusing term “critical time *τ* available for maximum acceleration” can be found in [1]. On the one hand, this sentence is misleading because the word ‘maximum’ implies an *ongoing* acceleration (variable ‘time’), whereas in their model, *τ* seemingly defines the *end* of the acceleration process (a parameter). On the other hand, acceleration ability had already been introduced in their eq. 3, on the basis of purely mechanical observations (see above: [145, 146]), denoted by *k*. Hence, 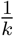 and an exponential time constant *τ_acc_* for acceleration are simply equivalent 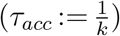: They represent a linear approximation [146, eq. 7] of any mechanical damping that counteracts the maximum possible driving force in the running direction (*F*_◦_ = *F_grf_* (v = 0); see further below).

Many mechanisms possibly causing such mechanical damping are conceivable. In [1], the mechanism of consuming ATP in white fibres and its depletion (there termed “fatigue”), was hypothesised to be the cause of the overall maximum in speed. This hypothesis was mathematically implemented by using *τ* as a limiting factor for acceleration in their data fitting function. In this, they assumed that an increase in *τ* is associated with an increase in muscle mass via the ATP energy storage capacity, which eventually yielded the necessitated power law ansatz (see [1, eq. 4]). That larger animals need more *total* energy (and, thus, muscle mass) than smaller ones to accelerate to their size-dependent maximum speed (kinetic energy: see Eq. (31)) is a mechanical fact, yet not a particular argument for setting an additional limit on *available* acceleration time. In the present muscle-mass-scaling context, the power law slope *g* = *g^lit^* was narrowed down in [1] by literature sources to between 0.76 and 1.27. Concordantly, our model predicts *g^model^* = 1, since 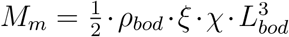 (substitute Eq. (E.3) and Eq. (E.4) into Eq. (J.1)) as well as 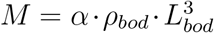, with, hence, *M_m_* ∝ *M*, and, thus, 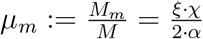 being a mass-independent constant.

Now, comparing the literature data and our model prediction with [1], we see an optimally fitted value *i^fit^* = −0.6 from their table S4, which clearly falls outside its mechanically and physiologically expectable range: *i^lit^* = *d^lit^* − 1 + *g^lit^* ∈ [0.42, 0.93] (for *d^lit^* see **Ad (2)** above). Indeed, a direct consequence of Hirt et al.’s mathematical setup is that *g* = *i* − (*d* − 1) must be negative in order to obtain a hump shape for which then again *i< −b* ≲ −0.3 must be fulfilled (with *b* ≈ 0.3 for the legged speed allometry v_*max*_(*M*) given, see **Ad (1)** above and again Sect. 3.3).

#### A plain model of ATP depletion during acceleration and its time frame depending on size

We now want to ‘repair’ and carve out what *τ* may, in fact, represent by deriving it from a mechanism. For this, we catch up on quantitative energetic considerations, both of metabolic (*E_met_*) and mechanical (*E_mech_*) nature. We clearly distinguish between the time scale for acceleration *τ_acc_* and the time scale for ATP depletion *τ_dep_*. Furthermore, we refrain from using the term *fatigue* because we focus on one specific metabolic process (ATP depletion) rather than taking diverse and interwoven physiological fatigue phenomena into consideration.

We start our ‘repair’ by estimating the metabolic energy release (consumption) from muscles per time:

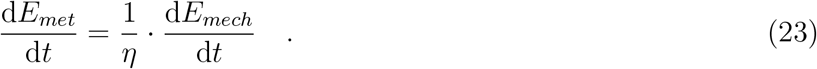

This plain ansatz is similar to what was proposed in [147, eq. 1] and can likewise be divined by comparing fig. 4 with fig. 6 in [148]. In Eq. (23), we further assume that the overall mechanical efficiency *η*, which covers all energetic losses from ATP split to a change in an animal’s forward speed, i.e., a gain in the kinetic energy of its COM, can be written as

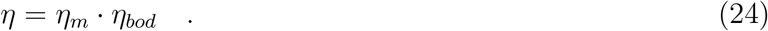

That is, we assume *η* to map a sequence of at least two separable processes: (i) the release of free energy by ATP split, with a skeletal muscle’s mechanical efficiency *η_m_* ≈ 0.4 0.6 [21, 22, 149, 150, 151], and (ii) the result of energy conversions within and around the body, which induce energy dissipation (damping): *η_bod_*. Such damping is caused by air drag and by friction that inevitably accompanies all tissue deformations, which occur, for example, in tendon stretch-recoil dynamics, in motions within and between muscle compartments, in motions of muscles relative to bones and the limbs relative to the trunk, and within the foot and in its sole pads. For our calculations of metabolic energy (ATP) consumption by muscles in accelerated running, with directly coupling consumption to mechanical work generated on the body scale (Eq. (23)), we chose *η_m_* = 0.5, *η_bod_* = 0.5, and thus *η* = 0.25 (Eq. (24)) as a default. A retracing of the latter two values to experimental data is provided in App. K.

In our plain mechano-metabolic model, we use the linearising assumption [146, eqs. 6,7], [152, p. 34]

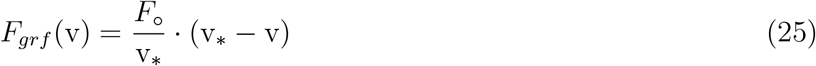

to model the GRF (the component in the running direction), which has also been adopted by [1]. Here, v_∗_ = lim_*t*→∞_ v(*t*) is the final (targeted) running speed, which is theoretically achieved only after an infinite period of acceleration (approached asymptotically: Eq. (28)), and *F*_◦_ = *F_grf_* (v = 0) is the ‘isometric’ (maximum) GRF value. The mechanical power for accelerating the COM in the running direction is

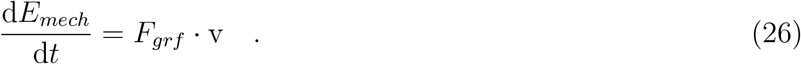

With the assumption Eq. (25), we can now calculate the solution v = v(*t*) to Newton’s equation

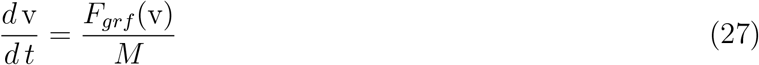

symbolically as [146, eqs. 6,7], [148, eq. 10, fig. 2], [152, p. 34]

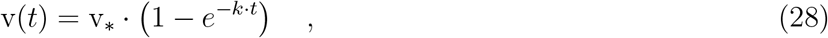

with *k* being the inverse of the *exponential time constant for acceleration*

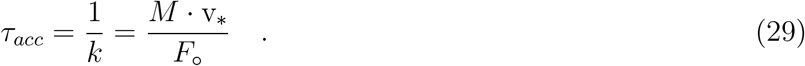

Because we simply assume proportionality between mechanical and metabolic powers (Eq. (23)), it follows that the change Δ *E_mech_*(*t*) in kinetic energy and the metabolic energy Δ *E_met_*(*t*) that is consumed during acceleration from v(*t*=0) = 0 to v(*t*) accumulate with time in proportion to each other:

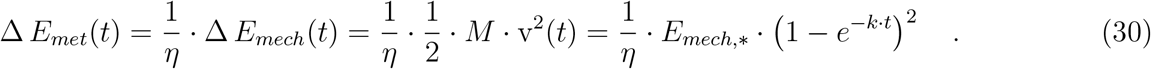

The symbol *E_mech,_*_∗_ in Eq. (30) represents the kinetic energy

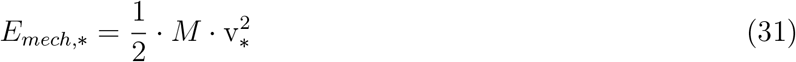

that corresponds to the final running speed v_∗_.

We follow the line of reasoning by [1] and go beyond. The metabolic energy available within a volume 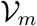 of muscle fibre material is

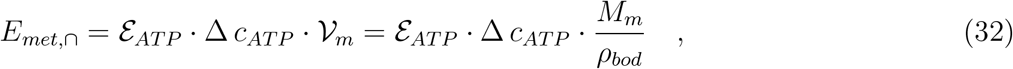

that is, *E_met,_*_∩_ scales linearly with the mass *M_m_* = *μ_m_* · *M* of the fibre volume. In the case of the present mechano-metabolic acceleration model, the values of 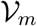 and *M_m_* represent all those muscles that contribute to doing work for acceleration. In the next paragraph, we will outline some considerations regarding the realistic choices of *μ_m_* values. The chemical energy stored per *one* ATP molecule seems to be at least 50 zJ, though 100 zJ [151, table 10] being much more realistic. In other literature, the numbers for the chemical energy *ε_ATP_* stored per *one mole* of ATP molecules have been given as 30 kJ mol^−1^ in older sources [153, 154, 155]. However, about 45 kJ mol^−1^ in fresh [149] or even fatigued [150, 156] muscle seems to be a much firmer (minimum) value, as 60 kJ mol^−1^ (corresponding to 100 zJ per ATP molecule) is a sound value according to more recent, comprehensive, and substantial reviews; see [156, p. 762] and [151, table 10], with the latter having even considered detailed sub-processes. The concentration Δ *c_ATP_* of ATP molecules in the muscle fibres that is available before reaching the fibres’ depletion level is *c_ATP,_*_0_ ≈ 1.2 mol m^−3^ [157, p. 291,305]. Note that we include in Δ *c_ATP_* the amount of ATP (approx. 7 mol m^−3^ [157, p. 305]) plus the amount of PCr (approx. 30 mol m^−3^[157, p. 305]), because PCr almost instantaneously re-builds ATP so long as it exceeds a minimum level of *c_PCr,_*_0_ ≈ 2.5 mol m^−3^ [157, p. 291,305]. In our considerations, we incorporate neither the re-build of ATP by other anaerobic processes nor the replenishment by blood flow. Thus, Δ *c_ATP_* ≈ (7 + 30 1.2 2.5) mol m^−3^ = 33.3 mol m^−3^ accounts for the least amount of available ATP resources.

Furthermore, according to our initial guess of the model parameter values (Table 2) for predicting maximum running speed, the value of the ratio of muscle to body mass in the running model is (Eqs. (5,J.1,J.2,E.3,E.4)) 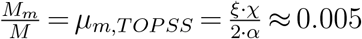 (i.e., 0.5%). This *μ_m,TOPSS_* value well matches *μ_m_* values found empirically for the ankle extensors of non-hopping mammals [158, table 1], which supports the idea that our model muscle is ‘the last one standing’ rather than a lumped muscle. However, during the sequence of steps involved in accelerating to maximum speed, more than one muscle works concentrically within a leg’s stance phase from roughly MS (ISOMS) to LO (TOPSS). The multiple-joint-multiple-muscle deployment in biological legs evidently reflects a mechanical design principle: Both a multi-joint chain and muscular redundancy implement the *spatial distribution of work generation*. Joints in a chain distribute work generation by partitioning actuator displacements in series, while muscles acting in parallel past a joint implement force-sharing among actuators (see also [121, p. 490]). Thus, both design concepts (chain of joints and muscle redundancy) implement the same thing: work-sharing. An approximate count of the muscles working in a real leg is this: In both vertebrate and arthropod legs, there are about three major leg joints in series, often with an additional proximal joint near or within the trunk, as well as at least one minor joint in the distal leg. Thus, at least five active muscles, distributed along the leg, will be synchronously active during stance to contribute to the work required for generating the same axial force as one lumped muscle in one representative joint. In four-legged animals, usually at least two legs do accelerating work in parallel during some period in a gait cycle. Therefore, in a real animal, at least two times five synchronised leg muscles are required to do work. Additionally, assuming some force-sharing by muscle redundancy across some leg joints (including bi-articular muscles, let us say: two per leg), the energy reservoir of accelerating muscle masses would be made of approximately fourteen muscles of the size of an ankle extensor, amounting to a minimum value of *μ_m_* = 14 *μ_m,TOPSS_*. In running with solely one-legged ground contacts, as in human sprinting, both arms and also some muscles in the swing leg will have to do work for purposes of keeping yaw and roll torques low, which, work-wise, may roughly compare with two synchronously acting legs (or maybe more in arthropods). In one-legged running, two legs are still deployed, although not synchronously. In running with whatever number of legs acting temporarily in parallel, trunk masses must support locomotion anyway. Thus, we would expect at least *μ_m_* = 2 · 14 · *μ_m,TOPSS_* = 0.14 as an absolute minimum value for the fraction of the body mass that does muscle work to accelerate the animal’s COM. In fact, muscle mass accounts for 32% of body mass in small mammals and primates, and even for 50% in bovides [158, table 4]. In arthropods, we are not aware of muscle mass counts. Instead, we have found ratios of overall leg to body mass in ants ([159]: 10%) and cockroaches (13%: [160]) However, as will be seen, ATP depletion is way off by orders of magnitude in such small animals. For our reasoning, therefore, only *μ_m_* values of larger animals are of interest. Hence, as a conservative guess (see also [161, suppl. table 1]), we choose *μ_m_* = 0.32 as a default parameter value in our acceleration model.

Eventually, by equating Δ *E_met_*(*t*) (Eq. (30)) and *E_met,_*_∩_ (Eq. (32)), we arrive at the energy balance that determines the instant *t* = *τ_dep_* of the depletion of the metabolic energy reservoir in muscle fibres during acceleration:

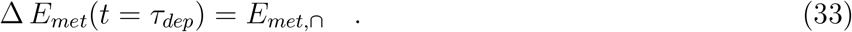

This event can occur only if 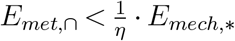; we can then calculate the instant of depletion during acceleration by solving Eq. (33) for

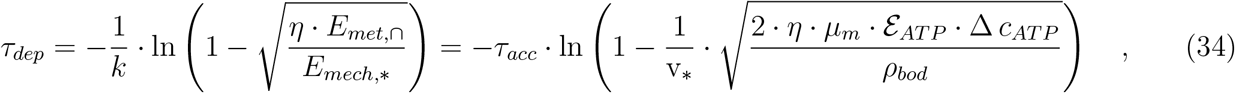

which implies substituting Eqs. (29,30,31,32) into Eq. (33). In the case of 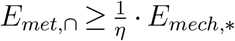, ATP depletion does not play a functional role in accelerating to (almost) v_∗_, and *τ_dep_* does not exist. In such case of non-existence, alternatively expressed, there is only one characteristic time that determines the dynamics of the system—namely, the purely mechanical one: *τ_acc_*. In the first instance, according to our estimate Eq. (34), *τ_dep_* is explicitly independent of *M*. However, there is an implicit *M*-dependency due to both v_∗_(*M*) and *τ_acc_*(*M*) or *k*(*M*), respectively.

#### Predictions of ATP depletion and concluding remarks

Now, after having introduced a mechanistic idea of ATP depletion, we can investigate the quantitative consequences, which are shown in Figs. 10 and 11. In Fig. 11, we have inserted into Eq. (34) for analysis only those two formulations of 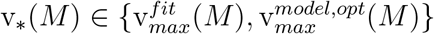 from Fig. 4 that predict at least a whiff of depletion; our standard solution 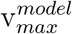 from the initial guess of parameters did not provide any indication of ATP depletion (see Fig. 10). Accordingly, we display in Fig. 10 a corridor of possible 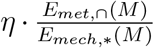 values based on 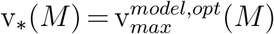. We use *η* = *η_m_ η_bod_* = 0.5 0.5 = 0.25, *μ_m_* = 0.32, *ε_ATP_* = 45 kJ mol^−1^, and Δ *c_ATP_* = 33.3 mol m^−3^ as the default (mean) parameter set and plot a lower boundary and an upper boundary of the corridor by, instead, assuming either *η_bod_* = 0.4 (thus *η* = 0.2) or *ε_ATP_* = 60 kJ mol^−1^, respectively.

**Figure 10:**
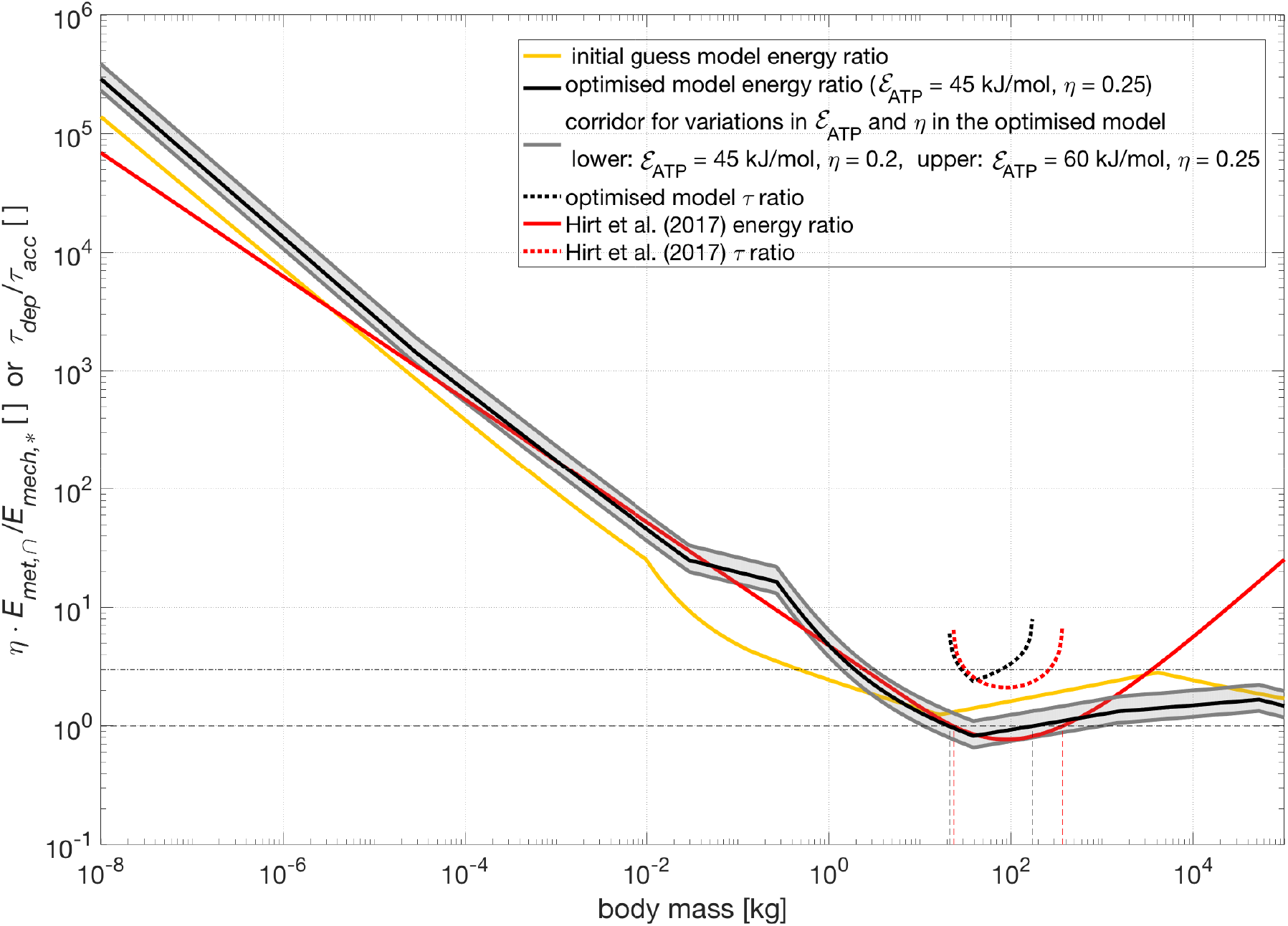
Model predictions of energy consumed during acceleration and instant of ATP depletion vs. body mass *M*. (i): The estimate *η* · *E_met,_*_∩_ of the mechanical energy (solid lines) that can be generated from the reservoir of metabolic (chemical) energy *E_met,_*_∩_ (Eq. (32)) stored in ATP within the whole-body muscle mass, with *η* being the estimated value of the mechanical efficiency for converting *E_met,_*_∩_ into work. The estimate *η* · *E_met,_*_∩_ is plotted as normalised to the kinetic energy *E_mech,_*_∗_ (Eq. (31)) that corresponds to the targeted speed v_∗_ always assumed to equal maximum running speed v_*max*_(*M*). We used the mechano-metabolic parameter values *η* = 0.25, *μ_m_* = 0.32, E*ε_ATP_* = 45 kJ mol^−1^ (i.e., 75 zJ free energy per ATP molecule), and Δ *c_ATP_* = 33.3 mol m^−3^ (see Sect. 4.4) for all three *η E_met,_*_∩_ predictions: v_∗_ = v_*max*_(*M*) was taken from either our initial guess model (solid orange line; in Fig. 4: ‘model without opt’, column three in Table 2), or our optimised v_*max*_ model parameter set (solid black line; in Fig. 4: ‘model opt’, column eight in Table 2), or the fit by [1] (solid red line; in Fig. 4: ‘Hirt et al. (2017)’). (ii): For the same three v_∗_ = v_*max*_(*M*) cases, an estimate of the instant *τ_dep_* (Eq. (34); dotted lines) of depletion of the ATP reservoir *E_met,_*_∩_ during acceleration from v(*t*) = 0 at *t* = 0 to v(*t*) = v(*τ_dep_*) at *t* = *τ_dep_* (event Eq. (33): Δ*E_mech_*(*t* = *τ_dep_*) = *η* · *E_met,_*_∩_). The estimate *τ_dep_* is plo(tted a/s normalise)d to the exponential time constant 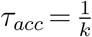 (Eq. 29) for acceleration, that is, 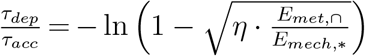 is plotted in case 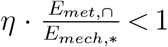 applies at all (*τ_dep_* a real, positive number). Further explanations of the data plotted are given in App. N.

**Figure 11:**
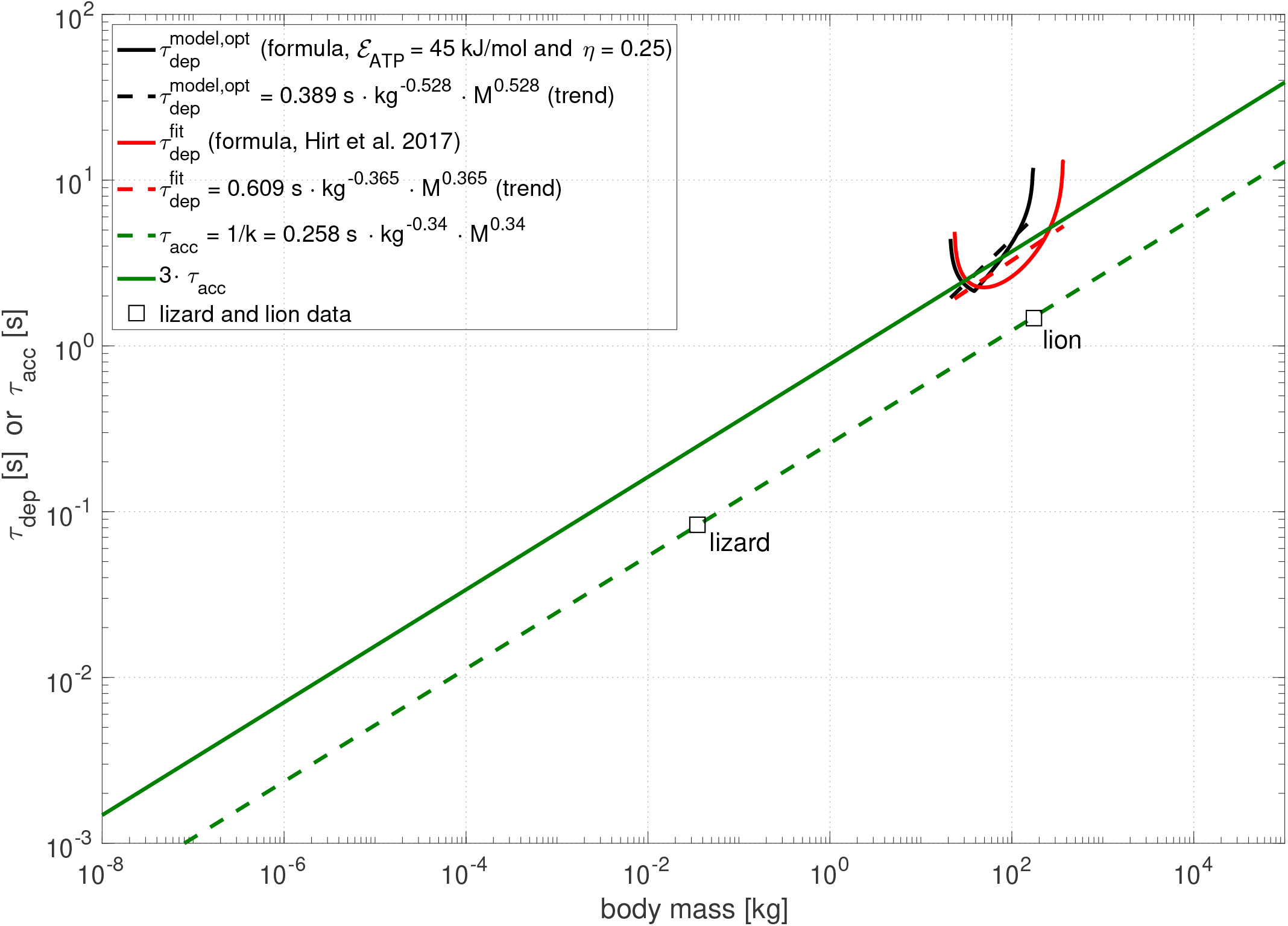
Model predictions of exponential time constant for acceleration and instant of ATP depletion vs. body mass *M*. (i): An estimate of the exponential time constant 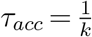 (Eq. 29) and its treble (dashed and solid green lines, respectively) for accelerating from v(*t*) = 0 at *t* = 0 to 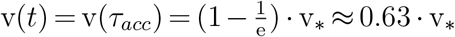 at *t* = *τ_acc_*, with v_∗_ the targeted speed. The parameters of the corresponding power law (Eq. (A.1)) were calculated from two known (*M*, v_∗_) data points (squares): empirical data of lizards [145] and lions [146]. Both v_∗_ values (lizard with *M* = 35 g: 3 m s^−1^, lion with *M* = 175 kg: 14 m s^−1^) in this minimal ‘data set’ are close to our model calculations of v_*max*_ and the v_*max*_ data fit by [1]. (ii; in Fig. 10, plotted as normalised to *τ_acc_*): Model estimates (using v_∗_ = v_*max*_(*M*) of ‘model opt’ and ‘Hirt et al. (2017)’ from Figs. 4,10) of the finite time horizon *τ_dep_* (Eq. (34); curved solid red and black lines) for depletion of the reservoir of metabolic energy *E_met,_*_∩_ (Eq. (32)) stored in ATP within the whole-body muscle mass. This time horizon is finite—and, thus, representable—only in the range of body masses for which the depletion condition 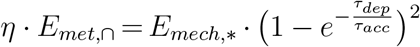 (Eq. (33) with Eqs. (30,31,32)) is fulfilled, with *E_mech,_*_∗_ being the targeted kinetic energy (Eq. (31)) that corresponds to targeted speed v_∗_. A whiff of depletion (i.e., a finite *τ_dep_* value fulfilling *τ_dep_ > τ_acc_*) is predicted by our plain depletion model (Eq. (34)) for the case of depletion occurring *later* than the instant *t* = *τ_acc_*. Since, v(*τ_acc_*) is only 63% of the targeted speed v_∗_, and while the animal will not achieve v_∗_ in this case, it will yet achieve more than 0.63 · v_∗_, 0.86 · v_∗_ for *τ_dep_* = 2 · *τ_acc_*, and 0.95 · v_∗_ for *τ_dep_* = 3 · *τ_acc_*. Using the default values *η* = 0.25, *μ_m_* = 0.32, *ε_ATP_* = 45 kJ mol^−1^, and Δ *c_ATP_* = 33.3 mol m^−3^ (see Sect. 4.4), our model predicts 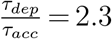 as a minimum, which corresponds to achieving a speed v of at least 0.90 · v_*max*_ in any case. (iii): Power law (Eq. (A.1)) fits (dashed straight red and black lines) to all available (finite) *τ_dep_*(*M*) values.

According to Fig. 10, both our optimised model (column eight in Table 2) and the fit by Hirt et al. [1] yield a *minimum* body mass of around 20 kg. For further increasing body sizes, a slight depletion before achieving v_*max*_(*M*) may appear in outlines. Furthermore and more importantly, a *maximum* body mass (our model: about 180 kg; Hirt et al. [1]: about 370 kg) always exists, which implies that depletion is *not at all* a limiting issue for the largest animals. Essentially, the allometry of the decisive ratio 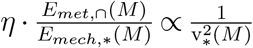 (see Eq. (34)) is nothing more than a flipped and distorted representation of v_∗_(*M*), i.e., practically v_*max*_(*M*)). Evidently, even for a decrease of v_∗_(*M*) = v_*max*_(*M*) in large animals as flat as predicted by our maximum running speed model, *τ_dep_*(*M*) is predicted by Eq. (34) to increase steeper than *τ_acc_*(*M*) with size. For immediate comparison, the ratio 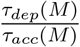 of the instant of depletion and the exponential time constant for acceleration is also plotted in Fig. 10 in the domains of *τ_dep_*(*M*) existence.

Figure 11 then shows the magnitudes of *τ_acc_*(*M*) and *τ_dep_*(*M*) themselves. We emphasise that their ratio is never lower than about 2.3 in our model prediction (see again Fig. 10). Thus, any animal can accelerate to at least 90% of v_∗_ (see Eq. (28) with (1 − exp(−2.3)) ≈ 0.90) before potential depletion. The steep ascents of the *τ_dep_*(*M*) curves at the boundaries of their domains (Fig. 11) occur due to the calculation of *τ_dep_*(*M*) according to Eq. (34). There, they tend to infinity as 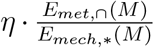 tends to the critical value 1 (Fig. 10). Hence, auxiliary regression curves are also plotted in Fig. 11 to visualise the unobvious mean tendencies. These boundary singularities may be artefacts of linearising the force-velocity relation within the acceleration equation examined in [146, eqs. 6,7]. Alternatively, it seems reasonable to consider the idea substantial that the accelerating drive (model: Eq. (25)) is probably a transformed version of the hyperbolic shape of the muscle’s well-known force-velocity relation [39] (Eq. (E.1)), which is incorporated (Eqs. (2,17,G.1)) into our model calculation of the legged speed allometry. As the main conspicuity, we obtain (on average) a *positive* slope, i.e., *τ_dep_*(*M*) *increases* with size, which directly contradicts the role of ATP depletion in the model setup of Hirt et al., as discussed above.

The whiff of depletion vanishing for larger animals is not only reflected by the sheer existence of only a potential mass range of depletion (20…180 kg or 370 kg, respectively). Already within this range, the trend towards a vanishing proximity to depletion with size is reflected by the mean slopes of the predicted *τ_dep_*(*M*) characteristics (Fig. 11) having at least the same magnitude as that of *τ_acc_*(*M*), both with using our model calculation of maximum speed and Hirt et al.’s curve fit. The slightly stronger whiff predicted with Hirt et al.’s curve fit is due to their v_*max*_(*M*) values being slightly higher than those from our model in exactly the mass range where depletion might occur.

In a nutshell, we find *τ_dep_* > 2.3 · *τ_acc_* for all animals including the largest, which lets them achieve at least 90% of v_∗_ as determined by muscle mechanics and air drag, while noticing, at worst, a whiff of depletion if not attempting to maintain v_∗_ = v_*max*_.

Summarising this section, we state that the idea, exposed by Hirt et al. [1], of ATP depletion explaining the decline of large animals’ maximum running speeds is unfounded. Their idea was led to a contradiction by engaging in a model calculation based on basic mechanics plus the known physiological properties of energy storage in the muscle. We suggest, instead, that a muscle’s own inertia in combination with other physical constraints (repulsion dynamics counteracting gravity and/or limited stance length: App. J; see also Sect. 3.2 and Sect. 4.3) causes the overall running speed limit for legged animals.

### 4.5. A point of view: predictions, explanations, and laws

In our introduction, we emphasised that we aim to provide a mechanistic—and, thus, causal—*explanation* for the legged speed allometry. We have also emphasised that we try to *predict* species-specific deviations from this characteristic. We feel compelled to elucidate the semantics of these introductory statements.

In our view, ‘prediction’ means predicting a *state of a dynamic system* (which may also be a static or steady state), which is represented by *state variables*, in response to a *variation* in the system’s *parameters*. According to this definition of a ‘prediction’, the variation can be realised in the real physical world (e.g., an experiment that can vary material properties rather than just the initial conditions of the state variables) or in a mathematical formulation that is assumed to represent the real system (a model).

This definition of a ‘prediction’ of a system state implies some issues: (i) The purpose of the ‘prediction’ is to understand real-world phenomena. By ‘understanding’, we mean ‘providing a *causal* explanation’ of the real-world system dynamics. (ii) Therefore, ‘causal explanation’ and ‘prediction’ are logically (or, maybe even better, epistemologically) interwoven. (iii) This connection is a consequence of the method of performing a ‘prediction’: A *parameter variation* in the real system or its model, respectively, allows making a statement about how the system *responds* if one of its *characteristic properties* is *changed*. (iv) These characteristics, depending on state variables and system parameters, determine the interactions within the system or with its environment. (v) To make ‘causal explanations’, the deployed model must be of a *mechanistic* type: Based on first principles, laws of nature, design criteria, and characteristic properties (e.g., force laws), it links the values of state variables (in this study, solely body mass) on the *independent preparation* side to corresponding *dependent response* values (in this study, solely maximum running speed) by means of mathematics. (vi) The *comparison* between the ‘prediction’ of a system’s response (represented by its state variables) and real-world measurements is mandatory for the process of ‘understanding’. (vii) Thus, a scientific approach that seeks ‘causal explanations’ by use of a mechanistic model is imperatively bound to formulating the model and, thus, the ‘prediction’ in terms of measurable state variables and, likewise, *principally measurable parameters*. (viii) Any violation of the latter requirement makes disproving the model impossible. (ix) The possibility of model disproof is mandatory for model *validity*.

In an empirical process, the repetitive measurement of system states provides a compilation of state snapshots. From quantitatively *analysing* this compilation, a potential ‘phenomenon’ may be extracted by a sorting process as, e.g., a statistical method. As a result, the ‘phenomenon’ is *described*, in the simplest case of a two-variable analysis, in terms of a fitting curve. Such a fit assigns one (state) variable that represents the *independent* system preparation (e.g., an initial condition, or body mass *M* in this study) in the experiment to one that represents the *dependent* response of the system (v_*max*_(*M*) in this study). A fitting curve may be termed a ‘model’ of the system, with the fit parameters accordingly representing the model parameters.

Once a model, whether mechanistic or not, is established, it can always be used to assign an independent state variable to another, dependent one—the system’s response or output, respectively. A scalar function (response, output) depending on one scalar variable (preparation, input) is the simplest case. For example, if the length of a linear spring is known, its force can be assigned. The stiffness and rest length are the only system parameters. They characterise the spring’s material properties. The spring’s characteristic is a force law in terms of length, stiffness, and rest length; by changing the state variable ‘length’, whether in the real world or in the spring model, the corresponding force value is gained by either interpolation or extrapolation. An ‘interpolation’ or ‘extrapolation’, respectively, is clearly distinct from a ‘prediction’. Both of the former denote the procedure of assigning, with given values of system parameters, independent state variables to dependent response variables. An ‘interpolation’ assigns *within* the range of measured data points, whereas an ‘extrapolation’ assigns *outside*. A ‘prediction’ is performed *by changing an initial value of a system parameter* (stiffness, rest length) and then calculating the system’s response to a given independent state variable (length) preparation just like in an interpolation or extrapolation. Any parameter value adjustment to the measured response data *after* the initial parameter value is fixed turns a ‘prediction’ into a fit. The requirements for ‘predictions’ to allow ‘causal explanations’ have been formulated above.

In empirical, just like mechanistic, approaches, a model is usually formulated as a mathematical ansatz, with the parameter values being gained by a mathematical procedure that fits such a model description to measured data points. In statistics, the model quality is assessed by the sum of deviations (residue) of all measured data points from the model fitted to them: the ‘unexplained variance’. According to the terminology in empirical, statistics-based approaches, the model fit ‘explains’ the measured data cloud, with a certainty given by the residue; what we would call an ‘interpolation or extrapolation’ is termed ‘prediction’ in these approaches.

If a process of a stochastic nature is part of a model, the term ‘prognosis’ is used instead of ‘prediction’. By its inherent properties, the model can then provide the values of probability for the accuracy of the ‘prognosis’ and should be labelled ‘stochastic’ rather than ‘mechanistic’. For a mechanistic model, the uncertainties of its ‘predictions’ are determined by the uncertainties of its input data, initial and boundary conditions, and parameter values.

A law of nature must fulfil a number of fundamental requirements [162]. One of them is that, to be revealed, a law of nature must be guessed in terms of a mathematical formulation that can explain measurable data: a model. Then, ‘predictions’ (according to our view elaborated above) of unmeasured system responses must be made using the model. Eventually, withstanding such repeated ‘prediction’ challenges, the law must be found to be non-rejected again and again. The legged speed allometry may turn out to be a general scaling law. Whether it can fulfil all requirements for a law of nature seems to be an open question.

A structure in the physical world is characterised by material properties, which determine interactions within it and with other structures. The most basic characteristics of a structure are its capacities for absorbing energy (nutrition in living organisms) and data (sensors), and the magnitudes of energy dissipation (heat loss) and internal information gain (local and transient entropy reduction). Regarding an animal that relies on legged locomotion, its own basic properties are the size (mass and characteristic length), leg length, strengths (e.g., stiffness) of its body materials, force and energy capacities of its actuators (muscles), and sensor and data processing (neural system) properties. Physical principles and laws in terms of its own and external properties constrain the structure dynamics. In legged locomotion dynamics, these constraints for mechanical performance are gravity and other external forces due to substrate contacts and airflow, as well as material failure loads and fatigue, characteristic times due to limited body size (and, thus, leg, stance, step, and stride lengths), whole-body inertia combined with leg stiffness, and both leg and muscle inertia combined with the muscle’s force-velocity relation. Subjected to these physical principles, laws, and constraints, the living structures explore the physical world, and evolution finds physiological solutions that enable the self-reproduction of the structures. The legged speed allometry is an expression of such solutions, which, as we are convinced, first and foremost indicates the basic physical shaping rules to which legged locomotion has ever been subjected.

## 5. Conclusion

In this manuscript, we have presented a strictly mathematically formulated, biomechanical model for predicting the maximum speed of any legged animal that runs or hops terrestrially, across the whole range of body sizes found, extant or extinct. We term the relation between maximum legged locomotion speed and body size the *legged speed allometry*. Our model consists of a plain representation of both air drag physics and the geometrical-anatomical structure of a generalised leg that gears the force generated by a Hill-type muscle model into a forward drive on the animal’s COM. Given its body mass *M*, the most prominent scaling factor is the animal’s characteristic length *L_bod_* assumed to be proportional to 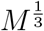, with *L_bod_*(*M*), thus, being the dimension that eventually represents the size of an animal (thinking of its trunk length). The values of all twenty-six relevant anatomical and biomechanical parameters entering our model were directly accessible from the available literature, which provided a reasonable initial guess. Refined values could be assigned to many of the parameters by the least-square-fitting of model solutions to the existing speed data [1] of about 200 species. Balancing the muscle-generated internal leg force with the body-external air drag, whileknowing that, on the one hand, the time required for a muscle to accelerate itself to high concentric contraction rates increases steeply with its mass whereason the other hand, only finite time is available for acceleration (due to the stance duration being limited by other, superposed mechanisms), we are able to predict the overall speed limit for running animals and the corresponding body size. In their paper, Hirt et al. [1] followed a similar goal. However, their result is a mere fit of parameter values that determines a descriptive double exponential function, which lacks a mechanical model foundation originating from first principles as well as a physiology- and structure-based parametrisation. In this respect, we are confident that the here-presented biomechanical model, which can calculate the *legged speed allometry*, including a prediction of the overall speed maximum, and predict the functional effects of diverse variations in body design, is suitable for bridging empirical data and their causal (mechanistic) explanation.

## List of abbreviations

≈: mathematical symbol meaning “approximately”
≳: mathematical symbol meaning “greater than about”
≲: mathematical symbol meaning “less than about”
COM: centre of (body) mass: spatial position
COP: centre of pressure: spatial location on the ground surface
CSA: cross-sectional area
DOF: degree of freedom
EMA: effective mechanical advantage
GRF: ground reaction force (vector: 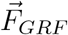)
ISOMS: running system’s state with its crucial muscle being isometric (around MS)
LO: (leg) lift-off: event of end of a leg’s stance phase
MS: (leg) midstance: event of half a leg’s stance duration after TD
TD: (leg) touch-down: event of start of a leg’s stance phase
TOPSS: top speed state of the running system (before, but close to, LO)

## Acknowledgements

We sincerely thank Monica Daley and Aleksandra Birn-Jeffery for them generously providing us with speed data of running birds. Michael Günther would like to thank two unknown DFG reviewers for their esteem, as well as Peter Aerts and Jim Usherwood for their dedication to inspiring discussions during the SEB annual meetings in 2018 and 2019. Furthermore, we would like to thank two unknown peers who had served as referees in a former submission to another journal: one for his concise yet thorough and very supportive reviews, and the other one for likewise supportive two rounds of detailed and in-depth review, which were tremendously helpful in polishing both the methods section and the whole story at many text passages. Those of us who have worked with Reinhard Blickhan love to ‘sign’ that a strong tune of his voice speaks through our words here. Reinhard has passed to us the key to the treasure box that has been filled by great minds in biomechanics.

## APPENDIX

## A. Scaling parameters with body size

In our model, we assume that each model parameter *p_n_* that is required to predict maximum running speed v_*max*_ can be either assumed to have constant values across body sizes or be eventually expressed as dependent on body size represented by body mass *M*. In our model formulation, this is symbolised by *p_n_*(*M*), i.e., the symbol of functional dependency (*M*) is attached to the respective parameter symbols introduced. Some of the parameters depend indirectly on *M* as they are assumed to scale with some multiple of the characteristic body dimension *L_bod_*(*M*) (Eq. (5)): e.g., *Re* (Eq. (B.5)), other lengths (Eq. (E.4)), areas (Eq. (B.3)), or area-dependent parameters (Eq. (E.2)). Some, like the drag coefficient *C_d_* (Eq. (B.4)) or the Reynolds number *Re* (Eq. (B.5)), have been taken or adopted from empirical models. Some are introduced due to considerations based on literature knowledge, like *A_rel_* (see Eq. (E.1) and Eq. (I.6)), Ω (App. C and see Eq. (8), Eq. (17)), or *c_con_* (see Eq. (J.3)). In this case, they are then assumed to scale, as typically done in empirical approaches, according to a power law

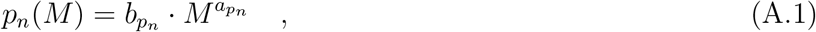

i.e., as straight lines in a log-log plot in which the values of both mass *M* and model parameter *p_n_*(*M*) are reported in terms of their logarithms along the abscissa and ordinate, respectively. Here, we always use the logarithm with base 10 for both axes in our plots. The relations

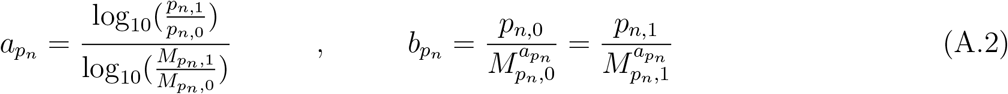

are useful for immediately extracting parameters in Eq. (A.1) from literature, if two data points 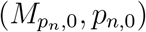 and 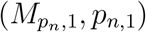 at a lower 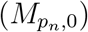 and higher 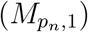 body mass are given.

Some of the parameters do only depend on *M* in a limited region of this whole range. In this paper, we use

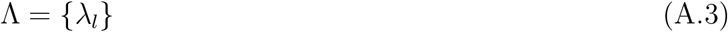

as a symbol for the set of all model parameters. The fact of the model parameter *λ_l_* being a member of the set Λ is then expressed in precise mathematical language as *λ_l_* ∈ Λ. Such concise notation for systematically addressing all model parameters is particularly helpful in applying an analysis of the sensitivity of the predicted model number (the solution: maximum running speed v_*max*_) with respect to any single model parameter (see Sect. 3.4).

In our model, each member *λ_l_* actually represents either the parameter *p_n_* itself if *p_n_* is always assumed to be a constant number (i.e., 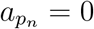 implying 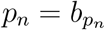) or one of the two constant numbers 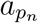 and 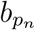 if *p_n_* = *p_n_*(*M*) according to Eq. (A.1) with no boundary values specified. For any *p_n_*(*M*) that becomes a constant at low or high masses, respectively, its lower or upper boundary value, respectively, is added to the set Λ. For capturing an *M*-dependent parameter in our model, between two and four model parameters are, thus, introduced. A synopsis of how many parameters are in fact used in our model is given in the **List of Abbreviations and Symbols (Table 1)**.

The choices of 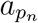 and 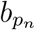 values—plus *p_n,_*_0_, *p_n,_*_1_ where applicable—for the size-dependent parameters *A_rel_*(*M*), Ω(*M*), and cos *φ_leg_*(*M*) (and two assumed size-independent: *B_rel_*, cos *φ_grf_*) in our model are summarised in App. D, and the choice of all model parameter values is compiled in Table 2.

## B. Parametrisation of air drag: A reduced model of body-external pressure and friction forces

Fluid dynamics is a demanding field of ongoing research. A classical textbook for an introduction to the field has been written by G. K. Batchelor [163]. For the purpose of our reductionist running model, we have broken down fluid dynamics to modelling the net drag force by airflow around the body as

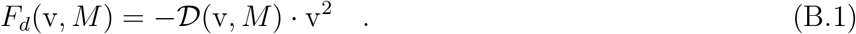

In Eq. (B.1), v is the running speed (forward or horizontal component, respectively, of the COM velocity) and

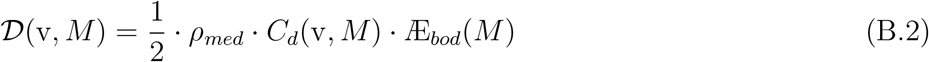

is the coefficient of damping for drag forces due to both pressure (normal to the body surface) and friction (tangential), which are exerted by medium (gas or fluid) flow on a body that moves with velocity v relative to the medium. In Eq. (B.2), *ρ_med_*, *C_d_*, and Æ_*bod*_ are the mass density of the dragging medium (here, air: *ρ_med_* = 1.2 kg m^−3^), the drag coefficient, and the projection of the body surface onto the plane perpendicular to the running direction, i.e., its cross-sectional area (CSA), respectively. In our model, the latter CSA scales as

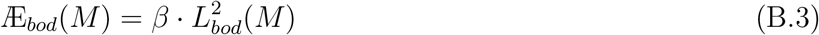

with body size. The initial guess 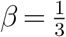 comes from the rough estimate that the area Æ_*bod*_ (Eq. (B.3)) of an animal facing airflow is approximately trunk length times a third of trunk length, which approximately reflects human-like proportions.

The drag coefficient

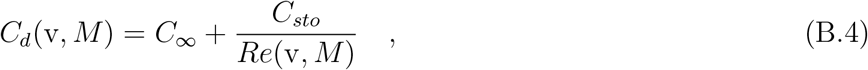

a proportionality factor in Eq. (B.2) and, thus, for the drag force *F_d_* (Eq. (B.1)), depends on the geometric shape of the body surface and the character of the flow of the medium around. The flow in incompressible media is characterised by the Reynolds number

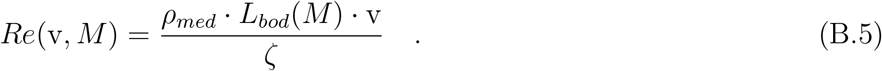

The assumption of the medium to be *incompressible* implies that the body’s speed v relative to the medium is neglectable as compared with the medium’s speed of sound (about 300 m s ^−1^ on earth). The Reynolds number *Re* can be physically interpreted as the ratio between the magnitude of a partial medium volume’s linear momentum (i.e., the flowing volume’s inertia) and the magnitude of the same volume’s linear momentum that is transported to the surrounding medium by pressure and friction forces, with the latter being characterised by a velocity gradient perpendicular to the direction of the inert linear momentum. As the body is the cause of such dragging transport (displacement) of medium volume, this physical idea of the Reynolds number can be immediately applied to the finite volume of the body, which must continuously sweep away medium volumes of its own dimensions. That is why the body length dimension *L_bod_* occurs in Eq. (B.5).

To find solutions v = v_*max*_ to Eq. (2) in closed form without changing the overall shape and asymptotic course of *C_d_* as a function of velocity, we used the ansatz according to Eq. (B.4). This ansatz neglects the 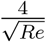 addend in the common empirical *Re* formula given by [164]. As *Re* (Eq. (B.5)) scales linearly with running speed v, *C_d_* depends in inversely linear proportion on v at low Reynolds numbers *Re*, or running speeds v, respectively, i.e., in the Stokes (friction) flow regime. Equation (B.5) as employed in our model is the simplest, empirically based approximation for *Re* possible. We used the value *ζ* = 1.8 · 10^−5^ kg m^−1^ s^−1^ for the dynamic viscosity of air.

At very low values of the Reynolds number below about *Re* ≈ 1, the medium flow around the body is laminar and stationary [165, sect. 12.4]. For values in the approximate range *Re* ≈ 1…30, the medium flow around the body is still laminar but periodical, i.e., with stationary vortices attached behind the body. In this range, drag (Eq. (B.2)) due to Stokes friction force, which goes in linear proportion to velocity v (see Eq. (B.2) with the right hand addend in Eq. (B.4) dominating), starts to gradually transform into the combination of friction and pressure drag in the Newtonian flow regime, in which the net drag force (Eq. (B.2)) goes quadratically with velocity v (see again Eq. (B.2) with the left hand addend in Eq. (B.4) dominating). In the Newton regime (Fig. 7: *M* ≳ 0.01 g), *C_d_ ≈ C*_∞_ becomes about a constant, which applies up to very high *Re* values at which further flow complexities occur. In our model, we used the values *C*_∞_ = 0.4 (Newton) and *C_sto_* = 24 (Stokes), which apply to a sphere. At values of about *Re* ≈ 40 (low-velocity region of the Newtonian flow regime), the flow is still regular, with vortices of a size similar to the body’s regularly and alternatingly detaching from the body into a ‘Kármán wake’.

The physical cause of vortex generation is that the tangential velocity of a medium particle that comes into contact with the body surface must be zero (relative to the surface), whereas particles in a distance from the surface move (at running speed) relative to the body. As a consequence, a (thin compared with the body dimensions) boundary layer builds up alongside the body surface, in which a velocity gradient is enforced in the medium. At the flow-exposed tip of the body, there is laminar flow in the boundary layer, at least up to the critical value *Re_c_* (see below). This (laminar) boundary layer grows thicker alongside the body the further away from the tip. Now, the thicker the boundary layer, the more likely do the medium’s viscosity and its particles’ inertia cause small-scale turbulences as a consequence of particle movement fluctuations (e.g., thermal) perpendicular to the flow strata, which are arranged tangential to the body and ideally parallel in the laminar boundary layer. This boundary layer part further away from the body tip is accordingly labelled ‘turbulent’. From the small-scale turbulent processes within the turbulent boundary layer part, bigger-scale vortices can grow by diffusion into the surrounding medium if physical conditions, as, e.g., the local ratio of inertia to friction forces in the flowing medium, allow the non-linear medium-medium-surface interactions to let turbulent medium volumes ‘organise’ themselves into bigger-scale structures. The higher *Re*, the more kinetic energy is stored per volume of the surrounding medium in turbulences outside the boundary layer, and the higher the local magnitudes of particles’ inertia forces, as well as the friction forces acting between them, which both somehow ‘fight against each other’. Accordingly, the flow structures around the body and in its wake change with increasing *Re* numbers.

If *Re* numbers increase beyond 100—or few times 100, depending on how the body length dimension is fixed—such local turbulent structures that are generated in the turbulent boundary layer also increasingly enter into the wake behind the body. The regular vortices in the (Kárman) wake are increasingly accompanied and superposed by small-scale turbulences (‘irregularities’), and the turbulent boundary layer grows more and more to the body tip. For *Re* numbers still higher than the critical value *Re_c_* ≈ 3…5 · 10^5^ (larger animals: Fig. 7), the laminar part of the boundary layer has eventually vanished, and the flow in the wake has narrowed to a fluidised bed that is entirely small-scale turbulent, without remainders of regular (bigger-scale) vortices. At even higher numbers (*Re* > 10^7^), new oscillatory phenomena on the basis of turbulence emerge. As not making a crucial difference for v_*max*_(*M*) calculations in the regime beyond *Re_c_*, we have still used *C_d_* = *C*_∞_ [165, sect. 12.4] here as a reasonable first guess.

## C. Axial joint alignment: The effective mechanical advantage (EMA)

The “effective mechanical advantage” [58]

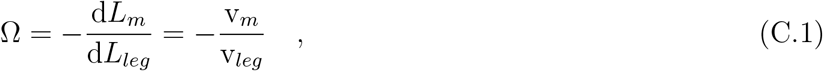

for a lumped extensor muscle *m* active at TOPSS has been factored into our model. The EMA transforms both a muscle’s length change (Eqs. (C.1,7,8)) and its force (Eqs. (14,17)) into the respective leg variables, with the latter determining COM dynamics (Eqs. (1,2)). The minus sign in Eq. (C.1) is due to muscle shortening (v_*m*_ < 0) being linked to leg lengthening (v_*leg*_ > 0) and vice versa. The EMA can be purely geometrically defined (Fig. 1): the leg length (*L_leg_*) change d*L_leg_* (> 0 at TOPSS: leg lengthening) per muscle length (*L_m_*) change d*L_m_* (< 0 at TOPSS: muscle shortening). The (functional) leg length is thereby defined as the distance between the most distal leg point COP (where GRF acts: may be anatomically approximated by a hoof, paw, or toe), and the most proximal leg point COM (where the leg does its axial work: may be anatomically approximated by a bony part usually most adjacent to the COM).

This combination of EMA and functional leg concepts moreover incorporates muscles and joints in a natural and mechanically consistent way. Any mechanical (usually rotational) degree of freedom can then be defined as a property ‘joint’ of a functional leg if such a joint can (geometrically) contribute to a respective leg length change. Accordingly, any muscle that crosses (actuates) a joint in a functional leg also becomes a property of the functional leg: basically, a source or drain of mechanical work. This integrated EMA-leg-joint-muscle concept makes it easy to get an illustrative view of it: Fig. 1.

Quantitatively, when any two adjacent body segments *i* and *k* within a multi-joint leg rotate relative to each other, as measured by a change of their relative orientation (joint angle) *φ_j_*_(*i,k*)_, with *j*(*i, k*) indicating this joint degree of freedom (DOF), the corresponding angular change d*φ_j_*_(*i,k*)_ concurs with the length changes

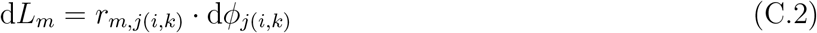

of all *M*(*j*) muscles *m* = 1,…, *M* (*j*) that cross joint DOF *j*(*i, k*) as well as the leg length change

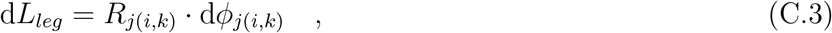

with *r_m,j_*_(*i,k*)_ and *R_j_*_(*i,k*)_ being lever arms (Fig. 1). Furthermore, the introductory EMA definition Eq. (C.1) also applies to any single muscle *m* of which the length change d*L_m_* is due to an angular change d*φ_j_*_(*i,k*)_ in *solely one* selected joint DOF *j*(*i, k*) as assumed in Eq. (C.2). By now substituting Eqs. (C.2,C.3) into Eq. (C.1) and therewith eliminating the angular change d*φ_j_*_(*i,k*)_ itself, we find

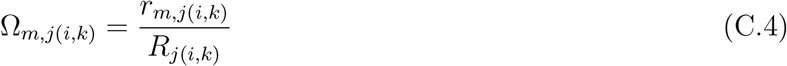

for the muscle *m* crossing joint DOF *j*. For a multi-articular muscle *m*, each *r_m,j_*_(*i,k*)_ is generally a function of the joint angles *φ_j_* of *all* those joint DOFs *j* that the muscle crosses.

Note that introducing just one, lumped muscle into our model allowed to omit any index to the symbol Ω in Eqs. (C.1,7,8,14,17). The explicit minus signs in these equations come from our model-specific definition of a lumped *extensor* muscle, together with the combination of signs chosen for concentric contraction velocity (v_*m*_ < 0) and corresponding contractile force (*F_m_* > 0) in the muscle’s force-velocity relation (see its various forms in Eqs. (I.1,I.2,I.3,I.4,E.1)). In the general consideration of length change transformations leading to Eqs. (C.1,C.4,C.5,7,8), specific choices to define joint angles as well as structurally determined relations between length changes in muscles and their effects on joint angle and leg length changes can be implicitly factored in to any leg model by choosing proper numerical values of the lever arms *r_m,j_*_(*i,k*)_ and *Rj*(*i,k*).

Angular change contributions d*φ_j_*_(*i,k*)_ of multiple joint DOFs add independently to leg length change:

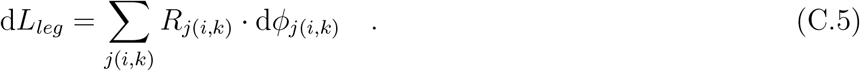

If a rotational centre of a joint DOF *j* crosses the leg axis, the numerical value of its geometrically defined lever arm *R*_*j*(*i,k*)_ changes sign together with the corresponding joint torque 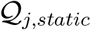 that would be required for static equilibrium in the leg. In this static case, the sum 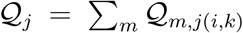 of the joint torque contributions 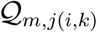 of the single muscles equals 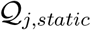.

In [58], a mean EMA estimate Ω_*j*(*i,k*)_ = ∑_*m*_ *w*_*m,j*(*i,k*)_ · Ω_*m,j*(*i,k*)_—summing over the contributions according to Eq. (C.4) of all extensor muscles *m* that cross the joint DOF *j*(*i, k*)—was introduced as an empirical value being representative for one joint in a leg. The summation over just the extensor muscles implied that all lever arms and, thus, Ω_*m,j*(*i,k*)_ were positive numbers. The weight *w_m,j_*_(*i,k*)_ for each muscle was its cross-sectional area (CSA) Æ_*m,j*(*i,k*)_, divided by the sum ∑_*m*_Æ_*m,j*(*i,k*)_ of these muscle CSAs, times a scaling factor in common for reproducing the experimentally determined joint torque. The use of this mean EMA estimate in [58, 74, 90] was, first of all, for describing the overall transmission of the forces of all (extensor) muscles that cross a selected leg joint to the corresponding joint torque 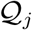. The other way round, the joint torque is a transformed measure of axial leg force *F_leg_* (see Eq. (14) with Eq. (9)) in nearly static situations 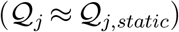. Further, the EMA was then employed for quantifying the found tendency of alignment of each leg joint to the leg axis (reduction in the *R_j_*_(*i,k*)_, i.e., increase in EMA: Eq. (C.4)) with increasing body size [58, 74, 75, 166].

Note that the concept EMA was introduced in [58] by estimating the joints’ lever arms *R_j_*_(*i,k*)_ as the perpendicular distances of the joints’ axes perpendicular to the GRF vector 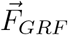, rather than to the functional leg’s axis. In contrast, we have here suggested to start from defining EMA solely by leg anatomy and geometry, and the single functional leg axis as defined by its respective COP and whole-body COM. Whereas the magnitude 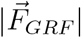 of the GRF vector and the axial leg force *F_leg_* do not differ substantially (see Sect. 2.5)even if there is significant misalignment of leg axis and 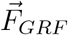, the lever arms *R_j_*_(*i,k*)_ may very well differ (Fig. 1), particularly the proximal ones. Notwithstanding this methodical issue, it was convincingly argued in [58] that the found increase in joint alignment with 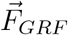 (straightly quantified by EMA) along with body size can be very likely traced back to the design criterion of maintaining maximum stress in the leg bones across a wide range of body sizes. Whether evolution, in fact, favoured joint alignment to 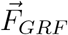 or to the functional leg axis from COP to COM may still be an open question. Being certainly a crucial evolutionary design concept anyway, enhanced joint alignment (increase in EMA)—whether to functional leg axis or 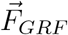—at a given level of required (axial) leg force reduces the statically required joint torques, with this, the isometric muscle force demands, and, thus, the metabolic energy consumption required for body posture maintenance and roaming locomotion, as metabolic consumption is dominated by the demand for active muscle force generation, essentially, supporting the weight [167, 168, 169].

## D. Choice of values for M-dependent parameters

The literature-based initial guesses of all model parameter values, be them given here or at other text positions in this paper, are compiled in the third column of Table 2 (‘initial guess’ *λ_l_*).

## D.1. A_rel_(M) and B_rel_

In our model, muscle mechanics is completely described by a hyperbolic force-velocity relation (App. E, see particularly Eq. (I.4)). We can use this relation as an empirical force law that applies also if the muscle has a tendon in series to the fibre material [40, fig. 5]. This observation is backed by a theoretical consideration: The functional form of a relation that varies and plots the values of an inert mass (a suspended load)—which is dynamically lifted against gravity by a pre-loaded elastic spring—versus the respective velocities maximally achieved during lift is very similar to a muscle’s hyperbolic force-velocity relation [170, appendix]. The crucial difference of a muscle-tendon complex to a muscle that solely consist of fibre material is that the value of the maximum concentric contraction velocity *V_m,max_* (Eq. (I.5)) of the approximating hyperbola approaches infinity with adding more and more elastic material in series (e.g., tendons with fixed CSA and increasing lengths). Here, we modelled this by simply letting values of *A_rel_* (Eq. (I.6)) approach very low numbers: As an initial guess, we assume *A_rel_* = 0.05 [37] below *M* 10 g, with the muscles being assumed to have no tendons in such small animals [72, 171], and decreasing values down to *A_rel_* = 0.001 at *M* ≈ 100 kg, which yields the values

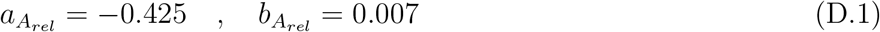

for the power law (Eq. (A.1)) parameters. Both boundary values are assumed to also apply as constants in the respective *M*-regions outside the transition interval (see Fig. 5).

The value *A_rel_* = 0.05 for pure fibre material seems to be very low as compared with, e.g., Hill’s original value of approximately 0.25 for frog fibres [39]. However, his experiments, and many other experiments since, were characterising isotonic (or isokinetic) *steady-state* contractions. It has been shown [36], however, that *V_m,max_* values are approximately eight times higher in the *non-steady-state* contractions that immediately follow rapid steps in force than in the (isotonic) steady-state contractions that finally follow these step responses, after the latter having faded away within few milliseconds. Moreover, in isometric *non-steady-state* contractions during increasing activity, values of approximately *A_rel_* = 0.05 have been found [37].

Note that we have used, as an initial guess, a constant value *B_rel_* = 5.0 s^−1^ for all body sizes, and look again at Eq. (I.5) to see how *V_m,max_* depends on parameters *L_opt_* (Eq. (E.4)), *B_rel_* (Eq. (I.7)), and *A_rel_* (Eq. (I.6)).

## D.2. Ω(M)

Scaling of the ‘effective mechanical advantage’ (EMA, see App. C; EMA symbol is Ω) is done using literature data [58, 74, 75]

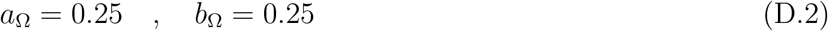

as the power law (Eq. (A.1)) parameter values that apply for mammals with body masses higher than *M* = 1.3 · 10^−1^ kg, with continuously increasing values up to an upper boundary at *M* = 4.1 · 10^3^ kg (see Fig. 5). That is, we have extrapolated the published power law [58, 74, 75] up to elephant-size animals, assuming Ω(*M* = 4100 kg) = 2.0. For masses lower than *M* = 1.3 · 10^−1^ kg, the constant value Ω(*M* = 130 g) = 0.14 is assumed.

## D.3. cosφ_leg_(M) and cosφ_grf_

The cosine cos *φ_leg_* of the angle *φ_leg_* that quantifies the kinematic constraint between COP and COM positions (i.e., leg axis, see Fig. 1) and running direction is assumed to change across the complete range of modelled body sizes. The angle is assumed to be, on average, flatter for smaller animals and steeper for larger ones. As an initial guess, we used *φ_leg_* = 40° as a first data point representing average animals of the size of the smallest mammals (Etruscan shrew with *M* ≈ 2 g), although there are some indications [67] that this may not be flat enough to represent a spider of the same size. As a second data point, we assumed that the average leg angle for animals of body mass *M* ≈ 100 kg may be somewhat between values for human runners (*φ_leg_* ≈ 55…60°) [92]) and racing greyhounds (*φ_leg_* ≈ 34°). The latter value was estimated by putting together length data from [95, fig. 2a] and leg configuration at LO plotted in [94, fig. 7,top]. Such leg angle values may also be found in big cats. As human body design is less representative in the animal kingdom, we, thus, estimated a second data point with a value of *φ_leg_* ≈ 45° being about the mean value of humans and greyhounds for an average animal of body mass *M* ≈ 100 kg. These data yield the parameter values

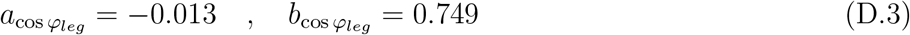

for the power law (Eq. (A.1)) and predict angle values of *φ_leg_* ≈ 18° for animals of the size of mites (*M* = 10^−8^ kg) and *φ_leg_* ≈ 50° for animals of the size of the heaviest dinosaurs (*M* = 10^5^ kg) (Fig. 6).

Like cos *φ_leg_*, the cosine cos *φ_grf_* of the angle *φ_grf_* of the GRF vector with respect to the running direction may change across the complete range of modelled body sizes. As our initial guess from literature, we assume, however, a constant *φ_grf_* ≈ 65° for all sizes (Fig. 6). Literature data on as much as accelerating spiders [67], running greyhounds and cheetahs [77], as well as human sprinters, both when accelerating and running at maximum speed [89], support this initial guess. This is not surprising as basic physical boundary conditions should be the reason: The ratio between the GRF component that a leg exerts tangentially to the running surface (*F_grf_*) and the normal component (*F_n_*) is restricted (*F_grf_ < ς* · *F_n_*) by the coefficient of static friction (*ς*), which is usually larger than the coefficient of kinetic friction (*ς_k_ < ς*) occurring if the leg slips on the surface. In horses competing during polo games [172], a static coefficient *ς* ≈ 0.6 has been found for hoof–polo-grass-field interaction. A ratio 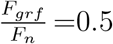 a little below such probably near-slipping-limit conditions corresponds to *φ_grf_* = 63.4° (≈ 65°) and cos *φ_grf_* = 0.447, respectively, which we, thus, do not only use as an initial guess but also as a constant parameter value throughout in our model.

## E. Parametrisation of muscle force: Hill’s hyperbolic force-velocity relation

The body-internal driving and dragging forces generated by the lumped leg muscle are assumed to be integrally characterised by the hyperbolic force-velocity relation as empirically formulated by A.V. Hill [39, 173] (see Eq. (I.1) and its particular form Eq. (I.4)):

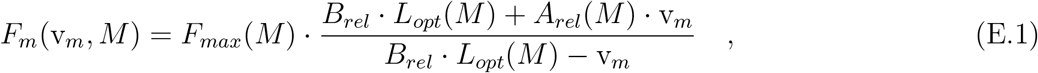

with v_*m*_ < 0 the concentric contraction velocity of the muscle, *F_max_* its maximum isometric force (i.e., the driving part of muscle force; we assume the crucial muscles to be *fully* activated), *L_opt_* the optimal length of the bulk of active muscle fibres, and 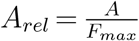, 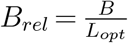 (values: App. D.1) the normalised asymptotes of the rectangular hyperbola, with *A* and *B* commonly being termed ‘Hill parameters’. The Hill parameters represent active and passive visco-elastic properties [38] and, thus, contain the dragging part of muscle force. In our model, we assume

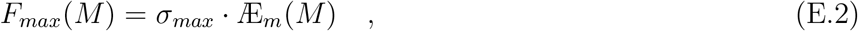

to scale with the muscle’s CSA

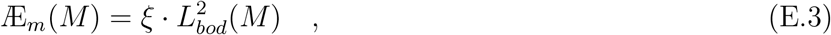

and optimal fibre material length

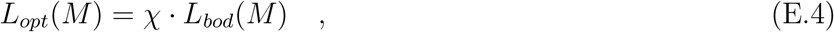

with our initial guess of the maximum isometric stress *σ_max_* = 2.5 · 10^5^ N m^−2^ (somewhere between the values given in these sources: [71, p. 145], [174, table 9.7], [175, p. 518] [176, 177, 73, 178], [179, suppl. mat.]) in vertebrate skeletal muscles, and the proportionality factors *ξ* = 0.0075 and *χ* = 0.15. The *ξ* value together with the chosen *σ_max_* value means that the modelled leg generates an isometric axial leg force value of one and a half times bodyweight in a human, with thereby implying a realistic human-like value Ω ≈ 0.7 [90] (see Table 3) for the ‘effective mechanical advantage’ (EMA: see App. C; EMA symbol is Ω; values: App. D.2). The chosen *χ* value means that the length of the (modelled) active leg fibres is assumed to be 15% of the trunk length.

## F. The projection of GRF vector 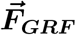 onto running direction

In a real multiple-leg animal, more than one leg may be on the ground at the same time, generating interlocking forces (“interaction forces” [102]) between the legs [103], with the net COP position depending on their load distribution. Likewise, due to a respectively extended contact area, the single leg’s COP may move even while just one paw or foot is on the ground. Furthermore, inertia forces generally occur within a leg, and joint torques may be exerted on the proximal anatomical leg by muscles originating close to the trunk, with the COM usually being located a distance away from any anatomical or functional joint. As a consequence, the GRF vector 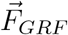 does generally not point in the same direction as the leg axis (Fig. 1), independent of movement direction and leg axis likewise non-aligning (model parameter cos *φ_leg_*, Eq. (6)). To account for the scalar product between GRF and COM velocity determining the work done by the GRF on the COM, we introduce the cosine cos *φ_grf_* as another model parameter (remember, Ω symbolises EMA, and see Eqs. (11), (12), (14, rightmost equality), (15), and (16)):

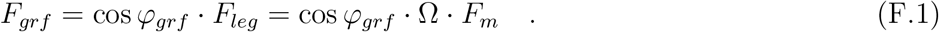

In other words, the parameter cos *φ_grf_* is the projection of 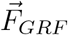 (and, thus, approximately the leg force Eq. (12)) onto the running (forward) direction, which quantifies in some sense the effectivity of a leg regarding running speed.

We have started modelling the force generation by the leg with the idea in mind that the magnitude 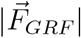 of the GRF vector is generally a good estimate of the force magnitude *F_leg_* generated along the leg axis (Eqs. (11,12)). Accordingly (Eq. (F.1)), the leg is the model structure that *primarily causes* GRF, apart from superposing inertia effects. Based on this, we use, on the one hand, the magnitude *F_leg_* of the axial leg force as one crucial multiplier for the forward-driving 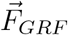 component *F_grf_* (Eq. (15)) in Eq. (F.1), and introduce just one additional parameter cos *φ_grf_* as a first approximation of the fact that, on the other hand, any anatomical definitions of leg axes, the functional leg axis (line from the leg’s COP to the COM), and the leg’s GRF vector are usually not at all aligned due to the physical effects mentioned above.

Notwithstanding, we have refrained from introducing a second angle-dependent multiplier when merging Eq. (11) and Eq. (15) into Eq. (16), which may seem an obvious one, namely, cos(*φ_grf_* − *φ_leg_*) occurring in Eq. (11), i.e., the ratio of the projection of 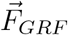 onto the leg axis, which we have defined as the ‘axial leg force’ *F_leg_*, and 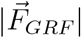 (Fig. 1). This multiplier would always effectively reduce EMA values as introduced by pure geometry (compare thick dashed green with solid blue lines in Fig. 1). However, due to (i) uncertainty whether this multiplier always properly represents the cause-effect chain from muscle drive to GRF generation, while inertia effects and proximal torques on the anatomical leg are present, which are already approximated at this stage of our model by one additional multiplier cos *φ_grf_*, (ii) more uncertainty about the multifaceted influence of the actual positions of all leg joints on a net EMA value, which might not properly be taken into account by exactly the cos(*φ_grf_* − *φ_leg_*) multiplier, and (iii) an effective reduction of EMA by cos(*φ_grf_ − φ_leg_*) not being higher than approximately 20% in a wide size range (Fig. 6), we have neglected this potential multiplier for the time being.

## G. The equilibrium between muscle force and air drag, and its solution (v_*feq*_(M)): Maximum running speed up to a size limit

By now substituting Eq. (8) into Eq. (E.1), the resulting *F_m_*(v_*m*_(v, *M*), *M*) then into Eq. (17), and again the resulting *F_grf_* (v, *M*) together with Eq. (B.1) into the basic force equilibrium Eq. (2), the latter explicitly writes

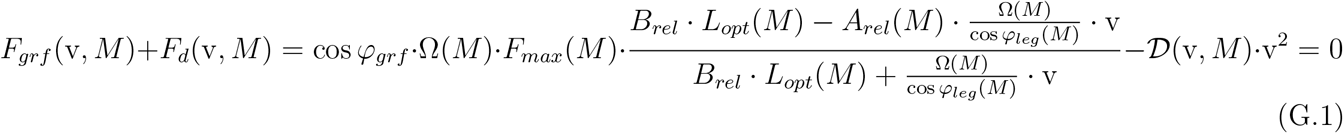

Further on, substituting 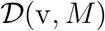 in Eq. (G.1) by Eq. (B.2), in the resulting expression then *C_d_*(v, *M*) by Eq. (B.4), and therein eventually *Re*(v, *M*) by Eq. (B.5), yields the force equilibrium Eq. (2) or Eq. (G.1), respectively, written solely in terms of the model parameters representing fixed numbers (structural properties), the body mass *M* representing an independent variable, and the resulting dependent variable v that we are searching for. By now multiplying, after all these rearrangements, Eq. (G.1) by the denominator of the first addend in Eq. (G.1) (i.e., *F_grf_* modelled as a transformed muscle force *F_m_*(v_*m*_(v, *M*), *M*)) and having taken notice that *Re*(v, *M*) is linearly proportional to v (Eq. (B.5)), it becomes obvious that Eq. (G.1) as a specific form of the force equilibrium Eq. (2) is a cubic equation in terms of the unknown variable *z* = v:

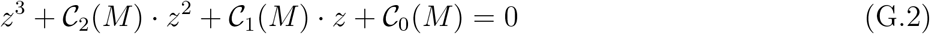

with the coefficients

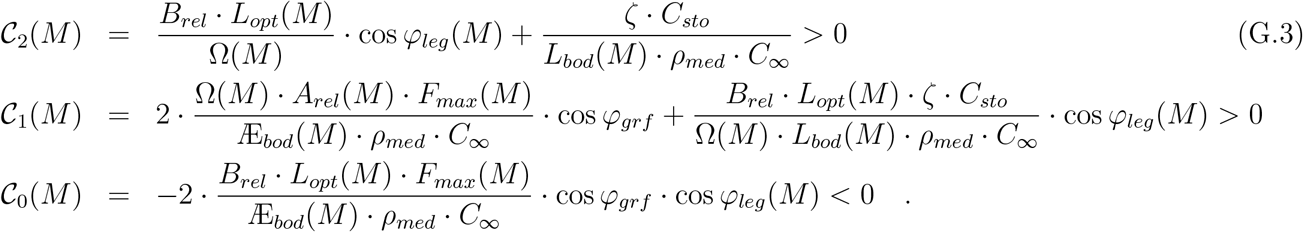

For all body masses *M*, the only real-numbered (and positive) solutions v of Eq. (G.1) are the ones with the coefficients 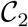, 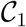, and 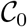 (Eqs. (G.3)) fulfilling *D* > 0 and *P* > 0 (Eq. (H.4); see App. H). At a given body mass *M*, maximum running speed v_*max*_(*M*) as calculated by our model from fulfilment of the force equilibrium Eq. (2) in its specific form Eq. (G.1), thus, would be the value v = v_*feq*_(*M*) of this real-numbered solution (Eq. (H.4)). Therefore, if no other constraint—like potentially Eq. (18) (see App. J)—is more speed-limiting than the force equilibrium Eq. (2), then we predict v_*feq*_(*M*) to be the natural value for maximum running speed: v_*max*_(*M*) = v_*feq*_(*M*).

## H. The solutions of interest to our basic equations Eq. (2) or Eq. (G.1), respectively, as well as Eq. (18) or Eq. (J.3), respectively: Solving the cubic equation Eq. (G.2) with the coefficients of either Eqs. (G.3) or Eqs. (J.9), respectively

The mathematical formulation in this section, in particular the concise, explicit presentations Eq. (H.4) and Eq. (H.6) of solutions to the cubic equation, has been adopted from https://de.wikipedia.org/wiki/Kubische_Gle G. Cardano [180] was the first to derive the symbolic solutions to the cubic equation, and the key for the following formulation of the concise solution classification and presentation has been provided by G.C. Holmes [181].

The three possible solutions to the cubic equation Eq. (G.2) are characterised by the discriminant

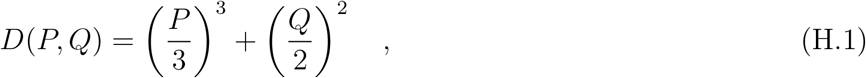

with the auxiliary numbers *P* and *Q* expressed in terms of the coefficients 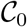, 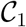, and 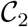 of the cubic equation Eq. (G.2):

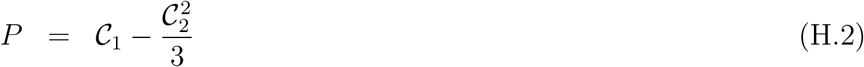

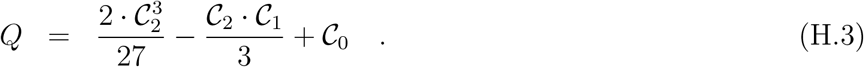

In case *D* ≤ 0 there are three real-numbered solutions (roots), otherwise only one root is real-numbered whereas the other two are complex-numbered. The solution of interest to calculate v_*max*_ from the force equilibrium (Eq. (G.1)) with the coefficients 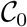, 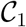, 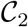 written down in Eqs. (G.3)) is given by the case *D* > 0 and *P* > 0:

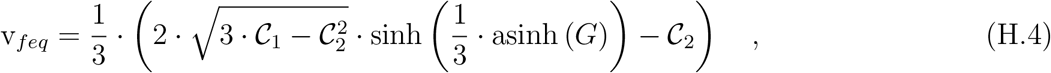

using another auxiliary number

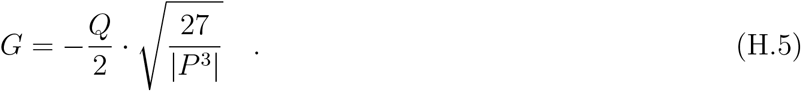

The solution of interest to calculate v_*max*_ from the muscle’s settle time becoming equal to half the leg’s stance duration (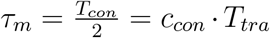: Eq. (18) with the coefficients 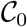, 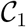, 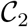 written down in Eqs. (J.9)) is given by the case *D* > 0 and *P* < 0:

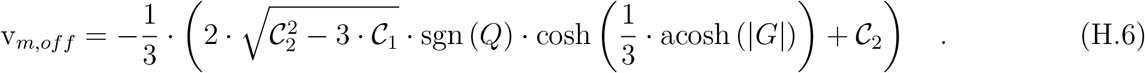

The kinematic transformation of inverting Eq. (8) eventually provides an alternative solution (Eq. (20))—to solving the force equilibrium Eq. (2)) in its particular form Eq. (H.4)—

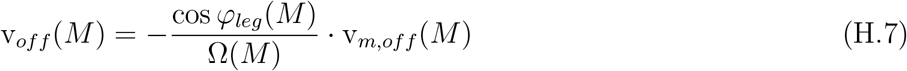

for maximum running speed v_*max*_, which is calculated from the condition Eq. (18) by particularly solving Eq. (H.6).

## I. The hyperbolic Hill relation and a muscle’s time to settle to a new force level

The force-velocity relation of a muscle is well-described by a rectangular hyperbola

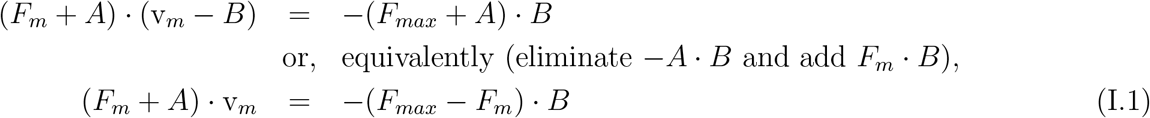

as first formulated by A.V. Hill [39] for frog muscle fibre contractions (pull: *F_m_* ≥ 0) in concentric (shortening: v_*m*_ 0) steady-state conditions. Since then, it has been shown to apply for single fibres as well as for bulks of muscle fibres and whole muscles including tendons in many experimental situations, even in non-steady-state conditions [36]. For our model calculations here, deviations from a single, exact hyperbola as, e.g., nearby the isometric condition [182, 183, 184] were neglected. The hyperbolic force-velocity relation is defined by three parameters: *F_max_*, *A*, and *B*. The parameter *F_max_* is the muscle’s (maximum, as a start [39],) isometric force (i.e., v_*m*_(*F_m_* = *F_max_*) = 0), and *A* and *B* are the magnitudes (positive numbers) *A* = − lim_v*m*→−∞_ *F_m_* and *B* = lim_*Fm*→+∞_ v_*m*_ of the rectangular force and velocity asymptotes, as introduced in Hill’s original formulation [39]. In our model, we use the hyperbolic force-velocity relation as a function

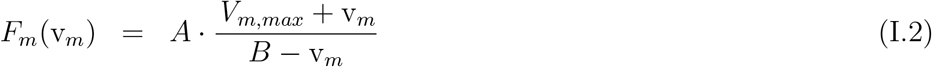

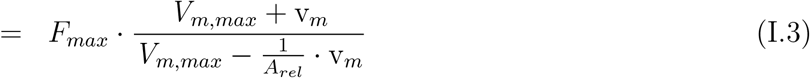

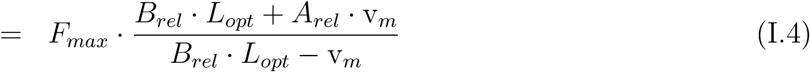

solved for the force *F_m_* as the dependent variable, with the contraction velocity v_*m*_ interpreted as the independent variable. In these formulations,

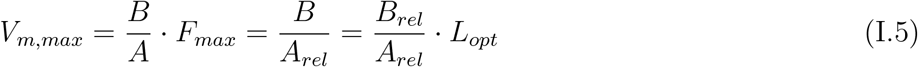

is the magnitude of the concentric contraction velocity for vanishing force (i.e., v_*m*_(*F_m_* = 0) = −*V_m,max_*) and

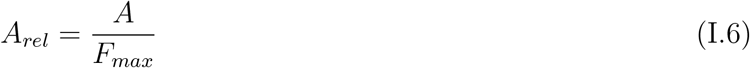

the normalised force asymptote. For also linking the Hill relation to muscle anatomy (thus, the muscle’s absolute geometrical dimensions), the velocity asymptote *B* and, therefore, *V_m,max_* (Eq. (I.5)) can be normalised to the optimal length *L_opt_* of a single fibre or the bulk of the fibre material (muscle belly), respectively:

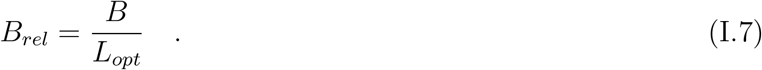

Note that

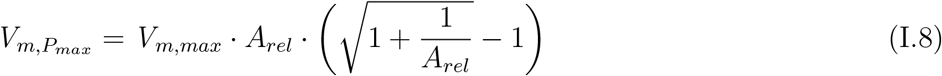

is the magnitude of the concentric contraction velocity v_*m*_ = −*V_m,Pmax_* at which the muscle generates maximum mechanical power *P* (v_*m*_) = −*F_m_*(v_*m*_) v_*m*_.

For further considerations, it is helpful and convenient to (i) note that the force-velocity relation fulfils

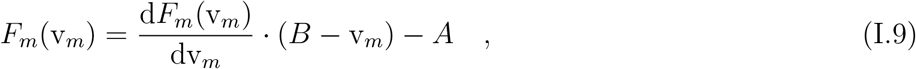

(ii) introduce the force-velocity relation in its normalised form (simply divide Eq. (I.3) by *F_max_*)

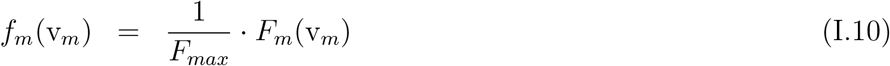

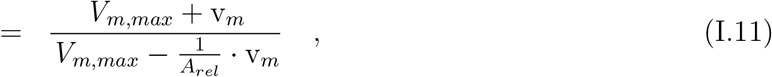

and (iii) calculate the latter’s slope:

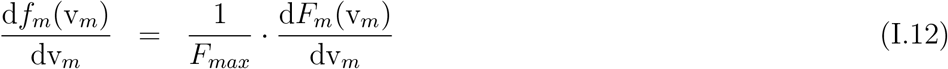

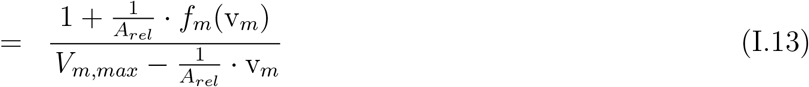

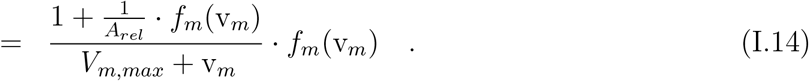

If the mechanical output of a muscle is characterised by the force-velocity relation Eq. (I.1) or Eqs. (I.2,I.3,I.4,I.11) respectively, then the dynamics to accelerate an arbitrary mass 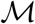 from initial (*t* = 0) rest (v = 0)—at which the isometric muscle force *F_max_* and the sum of all other external forces *F* (*t*) acting on the mass 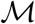 equilibrate (*H · F_max_* = *F* (*t* = 0))—to a lower value of external forces *H · F_m_* = *F* (*t* > 0) ≥ 0 is determined by Newton’s second law:

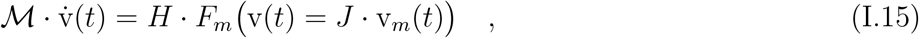

with the dot on top of the v symbol representing the time derivative of the velocity, i.e., 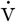 is the acceleration of 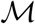. The symbol *H* represents a leverage between muscle force and force acting on 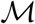, and the symbol *J* is an arbitrary but fixed transformation (constraint) *J* between the mass’ (v) and the muscle’s (v_*m*_) velocities. In this study, we interpret Eq. (I.15) to predict the dynamics of *the muscle accelerating its own mass*, which implies |*H*| = |*J*| = 1. It has been shown [185] that half of the muscle mass 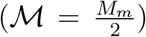 is a good approximation for the inertia that a muscle’s force output has to overcome when accelerating itself. With taking also the definitions (signs) of muscular force (*F_m_* > 0) and velocity (v_*m*_ < 0) during concentric contractions into account, which eventually yields *H* = −1 and *J* = 1, interpreting Eq. (I.15) as the dynamics that describes the muscle accelerating its own mass allows rewriting Eq. (I.15)—to gain Eq. (I.17), substitute Eq. (I.9) into Eq. (I.16), take notice of Eq. (I.12), and collect terms without the multiplier 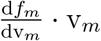 into *g_m_*(v_*m*_)—this way

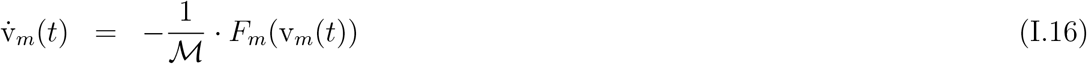

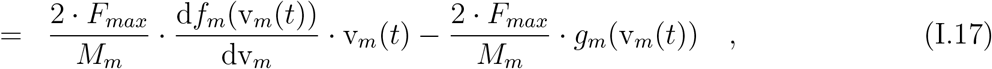

with the function

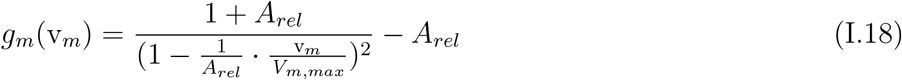

describing the intermediate force set points during the muscle’s settling dynamics Eq. (I.17) to the new force level *F_m_*(*t* → ∞) with the respective velocity v_*m*_(*t* → ∞). The multiplier

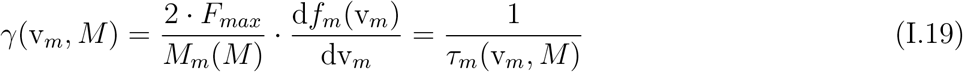

to v_*m*_(*t*) in the left addend to the right hand side in the dynamics Eq. (I.17) is the inverse of the time 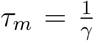 [185, appendix] during which the muscle mass *M_m_*(*M*) (Eq. (J.1)) settles to the new force level *F_m_*(v_*m*_) that corresponds to contraction velocity v_*m*_.

## J. A work restriction beyond muscular dissipation and air drag: Premature work termination due to muscle inertia and finite stance duration; a solution (v_*off*_) for maximum running speed above a size limit

In our model, the muscle mass *M_m_* is assumed to scale according to

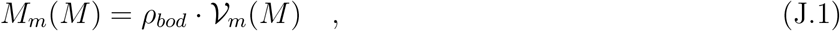

with the muscle volume

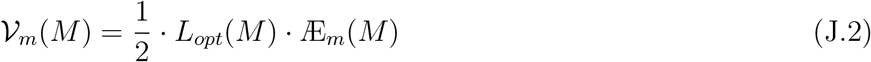

calculated from the muscle’s CSA (Eq. (E.3)) times optimal length of the bulk of fibre material (Eq. (E.4)) times a geometrical factor of 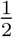, which is a reasonable approximation for an ellipsoid.

The time *τ_m_* (see Eq. (I.19) together with Eq. (I.14)) of a muscle settling from an initially generated to a different, newly demanded force level—i.e., the settle time for the muscle accelerating its own mass— depends strongly on both the demanded force *F_m_* or velocity v_*m*_, respectively, and on the muscle mass *M_m_*, which all scale with body size *M*: *τ_m_* = *τ_m_*(v_*m*_, *M*). The size dependencies of the two extreme cases *τ_m,min_*(*M*) = *τ_m_*(v_*m*_ = 0, *M*) and *τ_m,max_*(*M*) = *τ_m_*(v_*m*_ = −*V_m,max_, M*), respectively, are plotted in Fig. 3. Settle times are shortest in the isometric condition (*F_m_* = *F_max_*: *τ_m,min_*(*M*)) and increase monotonically with decreasing 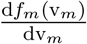 when approaching the unloaded condition (*F_m_* = 0: *τ_m,max_*(*M*)). Therefore, when the muscle accelerates through states from nearby the isometric condition at ISOMS to low force levels at TOPSS, the duration for the whole acceleration process is dominated by the very settle time that is associated with the final, low force level at TOPSS, since this is the longest one. Because the muscle can never fully achieve v_*m*_ = −*V_m,max_* due to external drag forces, we will always find *τ_m,min_ < τ_m,TOPSS_ < τ_m,max_* at maximum running speed v_*max*_ = v_*feq*_ as predicted by the force equilibrium (Eq. (G.1)). For the example of our model parameters taken as an initial guess form literature (column three in Table 2), Fig. 3 shows that the *τ_m,max_* line crosses our model estimate 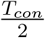 for half of the stance duration *T_con_* at a body mass of approximately *M* = 0.2 kg. For eventually capturing the condition of the running system, in which the muscle’s settle time equals half the stance duration, we first of all estimate the latter from our model parameters as

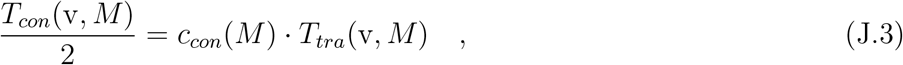

which requires to introduce just one additional, empirical parameter *c_con_*(*M*), and estimate the ‘travel time’

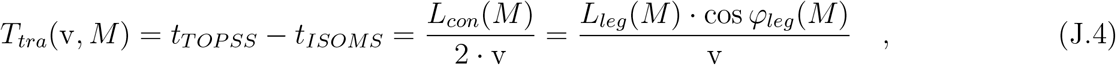

that is, the time period (*t_TOPSS_* − *t_ISOMS_*) it takes the COM approximately to move from the event ISOMS (assumption: *φ_leg_*|_*ISOMS*_ ≈ 90°, i.e., around MS) to the event TOPSS (model: *φ_leg_*|_*TOPSS*_ = *φ_leg_*(*M*)), with the leg length being approximated by a characteristic value *L_leg_*(*M*) (Eq. (J.6)) and the product

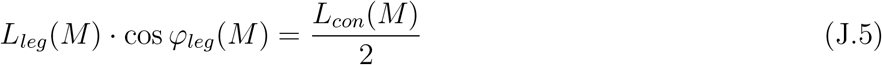

being our rough estimate of half of the stance length *L_con_*. Yet, the muscle’s settle time *τ_m_* becomes v_*max*_-limiting only at body masses higher than a critical value (Fig. 2: kink in v_*max*_ at *M* = 16.5 kg; Fig. 3: corresponding kink in *T_con_*). How in particular the finite duration *τ_m_* of accelerating muscle mass constrains maximum running speed more than the force equilibrium (Eq. (2)) is detailed in the following.

We start with rewriting our just-introduced empirical estimation for half the stance duration (Eq. (J.3)) by first substituting the ‘travel time’ *T_tra_*(v, *M*) (Eq. (J.4)) into Eq. (J.3) and, second, the simple approximation of a (characteristic) leg length (compare Fig. 1)

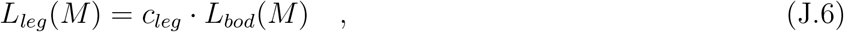

which yields

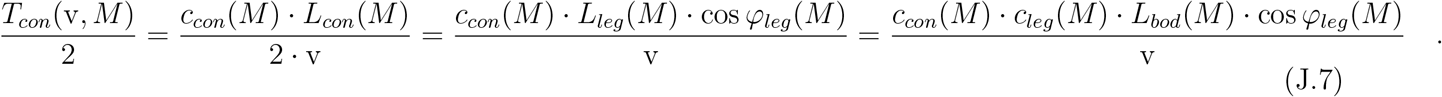

Eventually substituting Eq. (8) solved for v = v(v_*m*_, *M*) into Eq. (J.7) yields half the stance duration in terms of muscle contraction velocity v_*m*_:

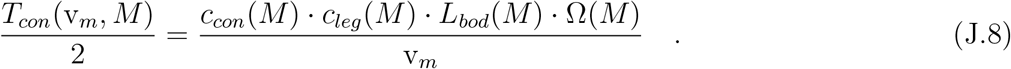

Without any frills, we chose, as an initial guess, the parameter value *c_leg_* = 1, which means that leg length in our scaling model would be always simply the same as characteristic body length (thinking of trunk length, see Sect. 2.2). Furthermore, we assumed *c_con_* = 0.75, not only as an initial guess but throughout. This is, because it turned out that this *c_con_* value can both roughly reproduce measured minimum stance durations of approximately 80 ms in maximum-performance human sprinters [78, 89, 32] of size *M* ≈ 80 kg and stance durations of approximately 10 ms in mites [59] of *M* ≈ 10^−8^ kg, which run at speeds of v_*max*_ ≈ 0.08 m s^−1^, with these speeds estimated by correcting down to 37° C and assuming that body, leg, and stance lengths are about the same (approximately 1 mm). Furthermore, a comparison of the green line in Fig. 3 to [30, fig. 5] gives evidence that our resulting estimate of *T_con_*(*M*) well matches measured data in the range *M* = 10^−1^…10^2^ kg.

Note that the solution v_*feq*_ to the force equilibrium Eq. (G.1) does not depend on leg length *L_leg_* at all (see Eqs. (G.3)), whereas the solution v_*off*_ to the constraint of the time available (stance duration) for accelerated muscle contraction Eq. (18) (denominated as *duration constraint*) does very well (see Eqs. (J.9)).

The muscle’s settle time *τ_m_* becomes v_*max*_-limiting if it exceeds half the stance duration 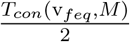 (Eq. (J.3)) that is predicted for the maximum running speed v = v_*max*_ = v_*feq*_ by the force equilibrium (Eq. (G.1)) between muscle and drag forces. From one perspective, this time limit due to muscle inertia can be explained this way: (i) The duration *T_tra_* (Eq. (J.4); ‘travel time’) of the leg axis rotating from approximately vertical orientation at ISOMS to an orientation according to cos *φ_leg_* at TOPSS, during which the COM accordingly travels on average by speed v, and which, therefore, continuously demands the corresponding contraction velocity 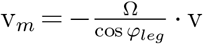 from the muscle, is assumed to be a proportionate estimate of half the stance duration 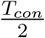 (Eq. (J.3)). (ii) If 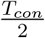 is not long enough to let the extensor muscle accelerate itself within its settle time *τ_m_* (Eq. (I.19)) up to the very value 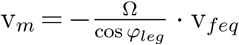 that corresponds to the state at which the (low values of) geared muscle force (GRF) of a single driving leg and the counteracting drag force equal (Eq. (G.1) fulfilled), muscle acceleration and, thus, work release is prematurely terminated. In other words, this fact may be expressed as: “*The animal would simply run too fast at* v = v_*feq*_ *for its crucial, potentially driving muscle(s) to accelerate in due time (until inescapable LO) and make the leg’s extension catch up with the COM movement*.” From another perspective, for the muscle and, thus, the leg to generate axial work until force equilibrium, it is required to always maintain the kinematic constraint between COP and COM (leg lengthening rate).

LO of a single leg can be enforced by mechanisms *other* than achieving the equilibrium of GRF and drag force components in running (*forward*) direction according to Eq. (2) (TOPSS), be it all the same TOPSS due to a single leg or a set of legs. In the latter case, TOPSS is caused by superposition of GRF contributions from several legs that interact among themselves. One alternative mechanism that may enforce LO before achieving TOPSS is the bouncing movement *perpendicular* to running direction, which is induced by the COM interplaying with gravity and the leg’s compliant, repulsive force-length characteristic [26, 27, 50, 114, 186, 187, 188]. Another basic time limit that potentially enforces LO comes from anatomy plus physiology: The leg length limits the stance length, which in turn limits the stance duration (Eqs. (J.3,J.4)), as the speed v is limited anyway by the actuators’ work capacities (Eqs. (1,2)). In any case, enforced LO entails termination of muscle work, which we then term ‘premature work termination’. Accordingly, an animal can only run with the speed v_*max*_ = v_*off*_ that corresponds to the value to which the inert muscle can accelerate itself during approximately half a stance duration, with the latter being potentially determined elsewhere.

In summary, by using the representation of half the stance duration Eq. (J.8) as a function of muscle contraction velocity v_*m*_, the condition of premature termination of muscle acceleration and, thus, work release can be accordingly written as Eq. (18):

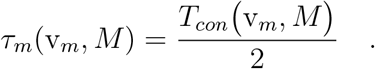

Like the force equilibrium Eq. (G.1) for a single functional leg (Fig. 1), the duration constraint Eq. (18) yields a cubic equation (Eq. (G.2)), however, here in terms of *z* = v_*m*_ for reasons of simplicity of the coefficients

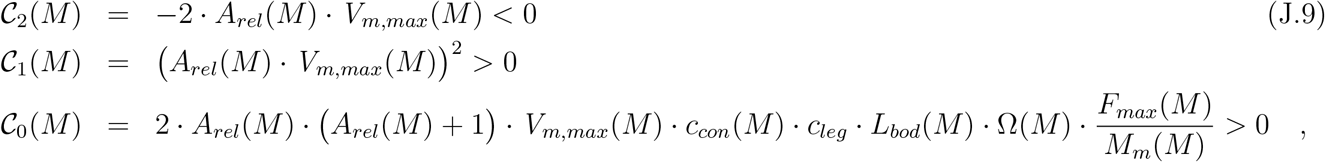

rather than in terms of *z* = v as in the case of the single leg’s force equilibrium Eq. (G.1). For all body masses *M*, the only real-numbered (and negative) solutions v_*m*_ of Eq. (18) are the ones with the coefficients 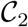, 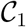, and 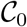 (Eqs. (J.9)) fulfilling *D* > 0 and *P* < 0 (Eq. (H.6): see App. H). At a given body mass *M*, maximum running speed v_*max*_(*M*) as calculated by our model from fulfilment of the duration constraint Eq. (18), thus, would be the transform (see Eq. (8)) v_*off*_ (*M*) (Eq. (H.7)) of this real-numbered solution v_*m,off*_ (Eq. (H.6)) to Eq. (18).

## K. Mechanical efficiency in muscle-driven legged running

A classical study [46] on human sprint running provides a first rough guess of a value for *η_bod_*, which is one of two multipliers in our model ansatz for the overall mechanical efficiency (Eq. (24)) of converting (with efficiency *η_m_*) free energy contained in an ATP molecule to mechanical work that the whole body of an animal does for accelerating itself in running direction. Cavagna et al. [46, fig. 2] suggested that, at maximum speed, about 20% of the mechanical power a human sprinter generates is dissipated by air drag [189]. Attributing, the same magnitude (20%) of energy loss to body-internal dissipation, we end up with our rough guess of *η_bod_* ≈ 0.6, thus, *η* = *η_m_ · η_bod_* ≈ 0.5 0.6 = 0.3.

Oxygen uptake is a reliably proportional measure of metabolic energy consumption during steady state locomotion in the aerobic regime. Applying this method, some early reliable values of experimentally determined overall mechanical efficiency in human sprint [190], steady-state walking [191], and running [192] have been in the range *η* = 0.23…0.25. Although stating differently there, a later study [193] of running at different speeds in the whole range up to sprinting, confirmed *η* = 0.2 0.33 (by re-interpreting their fig. 3,right,bottom with a more appropriate *η*-definition as external work divided by the sum of consumed metabolic energy and internal work). In fact directly based on the work of these predecessors, others [194] distinguished contributions of positive and negative external work in the running cycle to the metabolic costs, then also the costs of internal work [195], and applied a simplified version of the phenomenological relations from [194, 195] to locomotion at extreme slopes [196]. They all found overall efficiencies again in the range *η* = 0.22 0.24, partly based on assuming that *η* = 0.25 is generally a good estimate for phases of doing positive work. Therefrom, a phenomenological model [148] of metabolic costs in accelerated human running was developed, and a mean value *η* = 0.29 for 4 s of acceleration calculated (see their table 3 with assuming metabolic power of 25 W kg^−1^).

All these human data can be cross-checked by other direct measurements [169] of metabolic costs (oxygen uptake), in combination with stance durations, in an assortment of animals that all ran or hopped in a range of steady-state speeds. Kram & Taylor [169] found

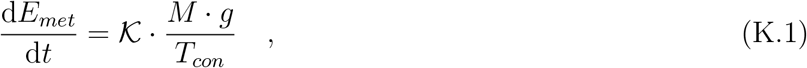

with *T_con_* (see, e.g., Eq. (J.7)) the stance duration of a leg, *g* the gravitational acceleration, and 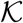 a phenomenological multiplier of unit J N^−1^. In legged locomotion, the work is always done by performing consecutive steps, and we are here anyway only interested in metabolic-consumption-to-work balances across a sequence of steps. The appropriate energetic measure is then power *integrated* over a sequence of steps, which can be dismembered into finite amounts of energy consumed or generated during each single stance phase, that is, a sum of stepwise contributions with a mean power value for each step. In accordance with this, 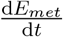 in Eq. (23) can well be replaced by the ratio 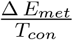 of the average amount Δ *E_met_* of metabolic energy consumed during a stance phase and the stance duration. Correspondingly, the right hand side in our ansatz Eq. (23) can be interpreted as a differential representative 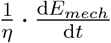 of the right hand side in Eq. (K.1), with Kram & Taylor’s 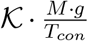, thus, representing a number in proportion to mean mechanical power released during a stance phase. In a nutshell, equating the right hand side in Eq. (23) with the right hand side in Eq. (K.1) and writing both of them in terms of average numbers for a stance phase, we can express the change in kinetic energy (work done)

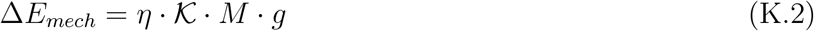

during *T_con_* in terms of parameters in [169]. Interestingly, Kram & Taylor [169] found in particular that 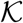 in their Eq. (K.1) is almost independent of speed *and* animal size. The idea behind our ansatz Eq. (23), thus, seems to be that the work per stance phase, at least in steady locomotion, is assumed to scale linearly with body mass *M*, which may be then again interpreted as scaling in fact with muscle mass *M_m_* = *μ_m_* · *M* (see also below), because the muscle to body mass ratio *μ_m_* is very similar [158, table 4] for the assortment of animals examined in [169]. Consequently, the mean mechanical power during one stance phase would then also scale approximately linearly with *M* (and potentially *M_m_*), and in inverse proportion to stance duration *T_con_*.

We aim here at finding the connection of *η* with *both* the parameters in [169] (particularly 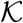) *and* those of our exponential acceleration model introduced in Sect. 4.4 (Eq. (28)), for the purpose of gaining a further guess of *η* from animal locomotion data. To this end, we predict Δ*E_mech_* in Eq. (K.2) by linearising the exponential function in v(*t*) = v_∗_ · (1 − *e*^−*k*·*t*^) (Eq. (28)) as *e*^−*k*·*t*^ ≈ (1 − *k* · *t*), which approximates the time course of COM velocity v(*t*) in the early acceleration phase (i.e., nearby *t* = 0) simply as v(*t*) = v_∗_ · *k* · *t*, and calculating therewith the change in kinetic energy between *t* = *t*_1_ and *t* = *t*_2_ > *t*_1_ as

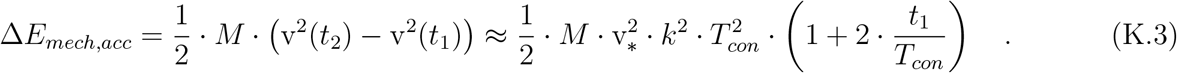

The term 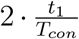 is a first-order error term due to the linearisation of v(*t*) in tandem with the *e*^−*k*·*t*^ function, which vanishes for *t*_1_ → 0. Hence, for *t*_1_ = 0, Eq. (K.3) provides an approximation of the change in energy

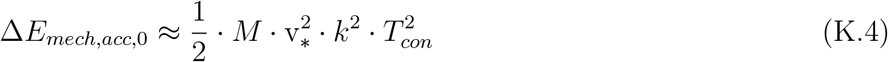

during the first step (more exact: stance phase) in acceleration. Equation (K.4) is a rough guess of how the work done during a stance phase scales with basic mechanical parameters, and it can now serve to estimate the relation between *η* and 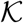. According to Eq. (29), the inverse 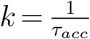 of the exponential time constant *τ_acc_* for COM acceleration depends on mass *M*, maximum forward-driving GRF component *F*_◦_ (force at v = 0, i.e., also nearby *t* = 0), and final speed v_∗_. With this, equating the work per stance phase as inferred from [169] (Eq. (K.2)) with the rough guess of Eq. (K.4), which has been derived from the mechanical part (right hand side in Eq. (30)) of our mechano-metabolic model for acceleration, yields the estimate

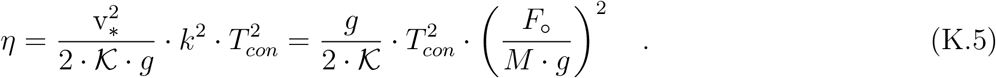

The rightmost expression in Eq. (K.5) is gained from eliminating v_∗_ by use of Eq. (29). Knowing 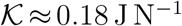 [169] and *g* = 9.81 m s^−2^, we can now estimate by Eq. (K.5) the value *η* ≈ 0.24 of the mechanical efficiency in the early acceleration phase of a human sprinter who performs at *T_con_* = 0.094 s [89, table 1] and an approximative value *F*_◦_ ≈ *M g* [89, fig. 1] of the peak force in running direction. From this, we conclude that the overall mechanical efficiency *η* is very similar in both steady-state and accelerated running conditions, and *η* = 0.25 is a good guess for estimating ATP depletion during acceleration according to Sect. 4.4.

## L. Further explanations to Fig. 2

The duration constraint Eq. (18) generally determines a running speed value v = v_*off*_ (*M*) at which *τ_m_*(v_*m*_(v, *M*), *M*) equals 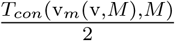. The v_*off*_ (*M*) solution that fulfils the condition 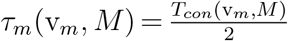 is depicted in Fig. 2 by the very thick green line. The v_*off*_ (*M*) course that is predicted for masses lower than the crossing point (at *M* ≈ 0.1 kg; overlaid yellow line) with the no-air-drag v_*feq*_ solution (thin dark blue line) exploits, in a mathematical sense, the fact that Hill’s hyperbolic muscle force-velocity relation (Eq. (I.1) or (E.1), respectively) allows (at least, mathematically) assigning solutions of *negative* muscle force values (*F_m_* < 0) that correspond to concentric contraction velocities *faster* than *V_m,max_* (v_*m*_ < −*V_m,max_*). Here, *τ_m_*(v_*m*_, *M*) > *τ_m,max_*(*M*) applies (the region to the left of the light blue line in Fig. 3), with very low 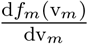 values. Such solutions imply that the muscle is *pushed* by an external force, rather than it does work on the COM. Accordingly, this high-speed solution branch is just academic—all the more so because the force equilibrium is generally more speed-limiting in this size range.

Both kinks in the course v_*off*_(*M*) are due to kinks in Ω(*M*) (see Fig. 5). For understanding the dependency of v_*off*_ on EMA Ω, we refer to Eq. (20) and remind of maximum muscle contraction velocity *V_m,max_* (Eq. (I.5)) generally increasing, *ceteris paribus*, along with *L_opt_* and, thus, body size. The second kink in the v_*off*_ solution at about 4 · 10^3^ kg is, therefore, due to the increase in Ω(*M*) terminating here in the ‘initial guess’ parameter set, i.e., Ω(*M*) becoming a constant (*M*_Ω,1_: Table 2) again. In this, we have already extrapolated the tendency of increasing Ω(*M*) beyond the upper mass case (horses) included in the data of [58, 74, 75]. Ω values for (extinct) larger animals are currently not available.

To see the impact of some crucial parameters on the solution v_*feq*_ to the force equilibrium, we have additionally plotted the cases of solely neglecting serial elasticity (‘no tendon’: for all sizes, the value *A_rel_* = *A_rel,_*_0_ of the smallest animals) for both neglecting air drag (‘no drag, no tendon’: thin violet line) and realistically taking it into account (thick violet line), as well as the two cases, with air drag (‘…, Ω_,0_’: thick black line) and neglecting it (‘…, Ω_,0_’: thin black line), respectively, in which *both* major determinants of small animal design are maintained throughout the size range: animals deploying just muscle fibres, i.e., having no tendons (*A_rel_* = *A_rel,_*_0_), as well as all animals assumed to having crouched legs (the value Ω = Ω_,0_ of the smallest animals).

## M. Further explanations to Fig. 4

In Fig. 4, note that the rightmost kink at *M*_Ω,1_ in any v_*off*_ (*M*) solution, which is due to Ω(*M*) increasing above *M* = *M*_Ω,0_ and becoming a constant at *M*_Ω,1_ again (for the ‘initial guess’: see Fig. 5), has been shifted in the optimal parameter sets to even higher *M*_Ω,1_ values (see Table 2) as compared with the ‘initial guess’ parameter set (orange line: ‘model without opt’).

Further note, that making the four mite data points in Fig. 4 comparable with mammalian conditions was done by using an approximately linear [96] dependency of v_*max*_ on temperature, with a *Q*_10_-factor of about 1.6 [59, 96]: *Parateneriffia spp.* (the smallest) with *M* = 3.0 · 10^−8^ kg [96] was corrected to v_*max*_ = 0.08 m s^−1^ as well as *Paratarsotomus macropalpis* with *M* = 1.8 · 10^−7^ kg [96] and *M* = 2.7 · 10^−7^ kg [59], respectively, were both corrected to v_*max*_ = 0.09 m s^−1^. Beyond [1], we have added a fourth data point, representing mites [55], to the experimental data cloud plotted in Fig. 4: The mites’ body mass *M* = 6.0 · 10^−8^ kg was calculated solving Eq. (5) for *M* with *L_bod_* = 0.8 mm known, and we corrected, as in the above cases, speed v_*max*_ = 0.042 m s^−1^ that was measured at about 25° C (cold-light illuminated in the lab) to an extrapolated value at 37° C: v_*max*_ = 0.08 m s^−1^. The iris, dash-dotted line depicts the power law (Eq. (A.1)) fit to data of solely those running birds (iris squares, triangles, and asterisks) that have been added beyond the data provided by [1]. Most of the additional bird data points have been generously made available by Monica Daley & Aleksandra Birn-Jeffery [197] (squares). One data point (triangle: very low speed, a kiwi) has been taken from [198, fig. 3], another four (triangles: ostrich, rhea, cassowary, emu) from their introduction (no references). One roadrunner point (asterisk) has been added from literature [79], and three further points (asterisks) from internet sources (two roadrunners: https://en.wikipedia.org/wiki/Roadrunner and https://www.youtube.com/watch?v=RU_7nZbII6o as well as a kiwi: https://a-z-animals.com/animals/kiwi/).

## N. Further explanations to Fig. 10

The time constant 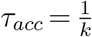 (Eq. 29), plotted versus body mass *M* in Fig. 10, is the time when 63% of the targeted speed v_∗_, which is only exponentially approached during acceleration, is achieved: 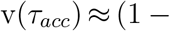 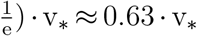. At 2 · *τ_acc_* and 3 · *τ_acc_*, v ≈ 0.86 · v_∗_ and v ≈ 0.95 · v_∗_, respectively, are achieved. We further assume that maximum running speed v_*max*_, as taken from either our model or the data fit by [1] (see Fig. 4), reasonably approximates the targeted speed value v_∗_. The case 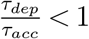 would indicate depletion to occur before *τ_acc_*, with then depletion possibly impeding the animal to achieve v = v_*max*_. By 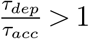 a finite time *τ_dep_* of depletion is predicted, which occurs *later* than *τ_acc_*. The condition 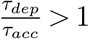, thus, means that the animal can accelerate, without being impeded at all until the estimated instant of depletion at *τ_dep_*, to a running speed higher than v = 0.63 · v_∗_. The body mass values at which the depletion condition *η* · *E_met,_*_∩_ = *E_mech,_*_∗_ is exactly fulfilled, and which, thus, enclose the size range within that depletion may occur, are indicated in Fig. 10 by dashed vertical lines for two calculated solutions v_*max*_(*M*) as well as for the data fit by [1]. Here, at least v = 0.90 v_*max*_ is achieved according to the predictions with our optimised model in the *M* range 20…180 kg. Outside this range, depletion is predicted to not interfere at all during acceleration to v_*max*_(*M*). With v_*max*_(*M*) taken from our initial guess model or with more recent estimations of 100 zJ free energy per ATP molecule [151, table 10] (upper boundary: *ε_ATP_* = 60 kJ mol^−1^), we would even predict depletion to not occur at all.

